# Brain functional characterization of response-code conflict in dual-tasking and its modulation by age

**DOI:** 10.1101/2023.03.21.533669

**Authors:** Lya K. Paas Oliveros, Edna C. Cieslik, Aleks Pieczykolan, Rachel N. Pläschke, Simon B. Eickhoff, Robert Langner

**Affiliations:** Institute of Neuroscience and Medicine (INM-7: Brain and Behaviour), Forschungszentrum Jülich, Jülich, Germany; Institute of Systems Neuroscience, Heinrich Heine University Düsseldorf, Düsseldorf, Germany; RFH – University of Applied Sciences, Cologne, Germany

**Keywords:** multitasking, aging, functional magnetic resonance imaging, individual differences, executive functions, effective connectivity, cognitive action control

## Abstract

Crosstalk between conflicting response codes contributes to interference in dual-tasking, an effect exacerbated in advanced age. Here, we investigated (1) brain activity correlates of such response-code conflicts, (2) activity modulations by individual dual-task performance and related cognitive abilities, (3) task-modulated connectivity within the task network, and (4)age-related differences in all these aspects. Young and older adults underwent fMRI while responding to the pitch of tones through spatially mapped speeded button presses with one or two hands concurrently. Using opposing stimulus–response mappings between hands, we induced conflict between simultaneously activated response codes. These response-code conflicts elicited activation in key regions of the multiple-demand network. Older adults showed non-compensatory hyperactivity in left superior frontal gyrus, and higher left intraparietal sulcus activity associated with lower attentional performance. While motor and parietal areas of the conflict-related network were modulated by attentional and task-switching abilities, efficient conflict resolution in dual-tasking was linked to suppressing visual cortex activity. Finally, connectivity between premotor or parietal seed regions and the conflict-sensitive network was neither conflict-specific nor age-sensitive. Overall, resolving dual-task response-code conflict recruited substantial parts of the multiple-demand network, whose activity and coupling, however, were only little affected by individual differences in task performance or age.

## Introduction

It is widely known that dual-tasking (i.e., performing two tasks simultaneously or in close succession) leads to interference, as reflected in speed and accuracy costs (Koch et al. 2018). As a process of higher-order action control, dual-tasking entangles other cognitive constructs such as attentional control, working memory, or inhibition, which are intertwined with the umbrella terms “cognitive control” and “executive functions” (Miyake et al. 2000; Himi et al. 2019; Saylik et al. 2022). However, the specific cognitive and neural mechanisms behind dual-task costs are not fully grasped yet and may be manifold. Previous studies have identified different sources of interference, such as a structural response selection bottleneck (Fagot and Pashler 1992; Pashler 1994), limited parallel processing capacity (Watter and Logan 2006; Miller et al. 2009; Koch et al. 2018), or crosstalk, which will be the focus of the present study (Pieczykolan and Huestegge 2018; Naefgen et al. 2022; Weller et al. 2022; Paas Oliveros et al. 2023). Crosstalk refers to the non-intentional information transmission between processing streams of different (sub)tasks (Navon and Miller 1987). It has been suggested that this mutual interference becomes more likely if tasks share physical features or conceptual dimensions, such as overlapping or conflicting response alternatives (Navon and Miller 1987; Paas Oliveros et al. 2023).

Dual-tasking difficulties are usually exacerbated with advanced age (Hartley 2001; Verhaeghen et al. 2003; Janczyk et al. 2018). Prominent cognitive theories have explained the age-related decline in dual-tasking through a wide range of mechanisms, such as the semanticization of cognition with a marked decline in cognitive control abilities but an accumulation and preservation of crystallized knowledge (Park and Reuter-Lorenz 2009; Spreng and Turner 2019), or a generalized slowing of information processing (Madden et al. 1992; Salthouse 1996). Other theories highlight an impairment in task-set shielding or attentional resource allocation due to an inhibitory deficit, leading to deterioration in scheduling attention across different task channels and to distraction among parallel processing streams (Hartley 2001; Mayr 2001; Mayr and Liebscher 2001; Hein and Schubert 2004; Maquestiaux and Ruthruff 2021), or process-specific changes, such as response-code confusability (Paas Oliveros et al. 2023).

Several neuroimaging studies have identified links between dual-tasking and brain activity in a fronto-parietal network, with a specific role for the dorsolateral prefrontal cortex (dlPFC) in the coordination of concurrent processes (Stelzel et al. 2006; Szameitat et al. 2006; Stelzel et al. 2008; Stelzel et al. 2009; Deprez et al. 2013; Al-Hashimi et al. 2015). Converging evidence shows that this network involves brain regions overlapping with the multiple-demand network (MDN; Duncan 2010; Camilleri et al. 2018), including the bilateral anterior insula (aI), dorsal premotor cortex (dPMC), anterior intraparietal sulcus (aIPS), and left inferior frontal sulcus and gyrus (IFS and IFG) (Worringer et al. 2019). The MDN has been implicated in goal-directed behavior, working memory, vigilant attention, and inhibitory control (Duncan 2010; Rottschy et al. 2012; Langner and Eickhoff 2013; Cieslik et al. 2015; Müller et al. 2015). Based on previous research, Worringer and colleagues (2019) proposed a neurocognitive processing model linking the brain regions found in their neuroimaging meta-analysis to different cognitive subprocesses involved in dual-tasking, such as attention shifting, working memory, stimulus–response (S-R) mapping, action planning, movement sequencing, and reorienting motor attention.

Here, we were particularly interested in investigating response-related dual-task interference elicited through mutually incongruent spatial response codes. This contrasts with earlier neuroimaging studies on dual-tasking, which have so far focused on multimodal input-related interference or interference at the central response selection stage (e.g., using the psychological refractory period paradigm). Therefore, dual-task crosstalk has remained understudied at the level of responses. Moreover, previous studies often reported relatively broad dual-versus single-task contrasts, which are not optimal for isolating specific interference processes but may englobe a myriad of higher cognitive functions associated with activity in MDN regions. In contrast, our approach allows investigating response-related dual-task crosstalk and specific behavioral and neural subprocesses therein without being confounded by potential stimulus-related interference, task order, or temporal overlap manipulations used in more traditional dual-task settings (Weller et al. 2022; Paas Oliveros et al. 2023).

Given that age-related deterioration in dual-task performance is a well-documented phenomenon, both in research and geriatric care, understanding the neural mechanisms behind this decline and its different facets is relevant for promoting and preserving older adults’ health. So far, however, the age-related neural changes explicitly associated with dual-task response-related crosstalk have remained largely unexplored, despite the dramatic impact the deficits at the response level can have on older adults’ everyday life (e.g., controlling devices, walking while talking, or risk of falling). In contrast, most neuroimaging studies investigating age differences in dual-tasking paired cognitive with motor tasks (van Impe et al. 2011; Papegaaij et al. 2017; Li et al. 2018).

Such studies have provided varied evidence of age differences in task-related brain activity, in which older (vs. young) adults sometimes showed reduced and sometimes increased brain activity (Grady 2012). Neurocognitive aging models have often interpreted hypoactivations among older adults as reflecting brain atrophy and cognitive deficits that come with age. On the other hand, hyperactivations in advanced age have been interpreted as either (beneficial) compensatory over-recruitment or (detrimental) neural dedifferentiation (Grady 2012; Spreng and Turner 2019). These two mechanisms have been most often discussed when assessing age differences during dual-task situations (Clapp et al. 2011; van Impe et al. 2011; Ohsugi et al. 2013; Chmielewski et al. 2014; Li et al. 2018; Thönes et al. 2018). The compensatory model predicts age-related brain hyperactivations in specific brain areas (especially prefrontal and/or contralateral regions) to counteract structural, functional, or cognitive decline in later life and enable successful task performance (Reuter-Lorenz and Cappell 2008; Park and Reuter-Lorenz 2009; Cabeza et al. 2018; Li et al. 2018; Heckner et al. 2021). For example, age-related prefrontal hyperactivations have been associated with a compensatory mechanism during motor control in dual-tasking, but it has been reported that older adults additionally recruit sensorimotor cortical areas (Seidler et al. 2010). In contrast, the dedifferentiation account assumes a loss of neural specificity when dealing with a given task, as reflected in widespread brain activation patterns, including task-irrelevant brain areas, such as more diffuse activation of visual regions among older adults (Park et al. 2001; Voss et al. 2008). This view holds that such hyperactivations in older adults should not be associated with better performance.

Thus, the first aim of this study was to study the neural correlates of response-code conflict in dual-tasking and their age-related differences. We hypothesized that if the above-mentioned neuroimaging meta-analysis (Worringer et al. 2019) reflected a network generally involved in solving dual-task interference, dual-task response-code conflict arising from mutually incongruent response codes in two concurrent tasks should recruit similar brain regions. If, however, the brain resolves different types of dual-task interference differently (i.e., in a conflict-specific manner), a less domain-general network would be expected to be recruited for our type of response-related dual-task crosstalk. Concerning age differences, it is important to emphasize that compensation and dedifferentiation models are not mutually exclusive. One could, for example, hypothesize that increased dual-task interference in advanced age (e.g., due to task-shielding deficits) is compensated by additional activity in action control regions, such as prefrontal and contralateral hand-specific premotor areas, whereas age-related reduced efficiency of sensory and motor activity in dual-tasking may lead to stronger crosstalk and thus, to dedifferentiation or more diffuse activation patterns in sensory and motor networks.

In a second step, we investigated how task brain activity was modulated by individual differences in dual-task performance itself and other cognitive abilities. As mentioned earlier, dual-tasking implicates several cognitive processes (Worringer et al. 2019) and shares variance with other subdomains of executive functioning (Himi et al. 2019; Saylik et al. 2022; Szameitat and Brunel Students 2022) to activate and maintain two task sets in parallel, control attentional processes and coordinate multiple actions. Furthermore, dual-task interference is affected by inter-individual differences and diverse contextual strategies to accomplish the tasks (Lehle and Hübner 2009; Hoffmann et al. 2020). Therefore, studying the associations between dual-task brain activity and related cognitive domains can offer insights into characterizing the cognitive and neural mechanisms behind dual-task crosstalk and their age-related changes (Miyake et al. 2000; Himi et al. 2019; Saylik et al. 2022) as well as the inter-individual differences and strategies implemented to cope with response-related dual-task crosstalk.

Multiple studies have analyzed the brain commonalities underlying cognitive demands of concurrent processes in dual-tasking and divided attention, highlighting the involvement of inferior and middle frontal gyrus, as well as IPS (Herath et al. 2001; Newman et al. 2007; Vohn et al. 2007). In addition, a recent study showed that left middle and superior frontal gyrus (MFG and SFG), medial frontal cortices, and inferior and superior parietal lobules are commonly recruited during tasks taxing one of four major executive functions (updating, inhibition, switching, and dual-tasking; Saylik et al. 2022). Furthermore, better working memory has been strongly associated with improved dual-task performance and, in combination, both cognitive abilities recruit dlPFC and parietal regions (Klingberg 2000; Deprez et al. 2013; Nijboer et al. 2014; Heinzel et al. 2017). Finally, the previously mentioned meta-analysis not only found brain regions uniquely associated with dual-tasking (bilateral dPMC, frontal operculum (fO), aIPS, and left IFS and IFG) but also brain areas that shared functional activity with task-switching performance (i.e., consecutively alternating between different tasks), comprising bilateral IPS, left dPMC, and right aI (Worringer et al. 2019). Thus, the similarity and difference in activation patterns across the intertwined cognitive processes in dual-tasking are likely related to behavioral interference resulting from specific task interactions in several brain regions (Nijboer et al. 2014). And this probably differs across age groups, since all these cognitive processes are known to decline with age (Craik and Salthouse 2008; Salthouse 2010; Cabeza et al. 2018; Spreng and Turner 2019).

Considering the previous observations, our second aim was to investigate how conflict-related brain activity in young and older adults is linked to individual dual-task performance and various related cognitive abilities. Specific sub-goals included: (1) to examine how much group-level response crosstalk effects in brain activity were modulated by individual dual-task ability and (2) other cognitive abilities associated with dual-tasking, as well as (3) to investigate age-dependent differences therein. We inferred that if conflict-sensitive brain areas, such as frontal or superior parietal regions, shared variance with other cognitive processes associated with dual-tasking, such as working memory or attentional processes, it would hint to brain activity non-specific to response-code conflict and possible inter-individual differences in cognitive strategies among participants to deal with crosstalk effects. However, if the task-related regions selectively co-varied only with dual-task ability, this would reflect a process-specific involvement of brain regions.

Besides regional brain activity, the question arises whether the integration and dynamic coordination between different parts of the task network is relevant for efficient dual-task performance. In fact, it has been established that functional connectivity patterns change with aging, potentially leading to cognitive deficits (Craik and Salthouse 2008; Li et al. 2018; Spreng and Turner 2019). For example, resting-state functional connectivity studies suggest that age-related reductions in interregional coupling between dlPFC, PMC, aI, and parietal regions may go along with a decline in cognitive action control (Langner et al. 2015). However, to our knowledge, previous research has not yet explored the task-related connectivity changes during response-code dual-task crosstalk and their modulation by age.

Somewhat related to this question, Breukelaar and colleagues (2018) assessed the relationship between changes in task-modulated connectivity and performance in tasks taxing cognitive control abilities over two years. The authors found that a longitudinal increase in task-modulated connectivity between left IPS, bilateral dlPFC, and anterior cingulate cortex (ACC) during a working memory paradigm was associated with improved executive functions over the two years. Another study found increased coupling between anterior prefrontal cortex and precuneus for a prospective memory task among young adults and decreased coupling for older adults, possibly reflecting age differences in top-down attentional monitoring processes (Lamichhane et al. 2018). In contrast, a more recent study on associative memory, including dual-task conditions with a tone detection task, found increased monitoring-related connectivity between three seed regions (ACC, left and right dlPFC) and bilateral occipital cortex and left IPS but neither task (single vs. dual) nor age effects (Horne et al. 2021). Thus, our third and final goal for this study was to explore whether task-modulated connectivity between dual-task conflict-sensitive brain regions changes depending on the dual-task conflict level. Again, we aimed to explore age-related differences in the synchronization of task-related brain areas.

In summary, our objectives were, first, to study brain activity correlates of response-code conflict and their age-related differences. Second, we aimed to investigate how this conflict-related brain activity and age differences therein are linked to individual dual-task performance and various related abilities. Our third and final aim was to explore whether highly conflict-sensitive brain regions change their connectivity to other task-relevant brain areas depending on the level of dual-task difficulty and whether this is affected by age.

## Materials and methods

### Participants

We recruited 51 young (age range: 20 – 35 years) and 51 older adults (age range: 50 –71 years) via advertisements and personal contacts. Participants received monetary compensation (45 – 60 EUR) for their participation in a larger project in which they completed an extensive battery of cognitive tasks, psychometric questionnaires outside the scanner, and various MRI sequences in two sessions. All participants had normal or corrected-to-normal vision and reported no history or presence of psychiatric or neurological disorders. All older participants passed the DemTect screening for dementia (Kessler et al. 2000), indicating they did not suffer from clinically relevant cognitive impairment (cut-off 13/18 points). The study complied with the Declaration of Helsinki and was approved by the local ethics committee of the RWTH Aachen University Hospital. All participants gave written informed consent before the experiment.

We excluded 21 participants committing ≥ 50% errors in at least one experimental condition from further analyses, attesting to the task’s difficulty despite the practice blocks presented outside the scanner. In addition, two participants were excluded because of low fMRI data quality. This left us with 43 young (*M_age_* = 25.6 years, *SD_age_* = 3.4 years; 22 females) and 36 older adults (*M_age_* = 61.9 years, *SD_age_* = 5.5 years; 15 females) for our analyses. In this analysis sample, 27 young and 30 older adults were right-handed, according to the Edinburgh Handedness Inventory (Oldfield 1971). For further descriptive statistics of the sample, see **Table 1.**

**Table 1.**
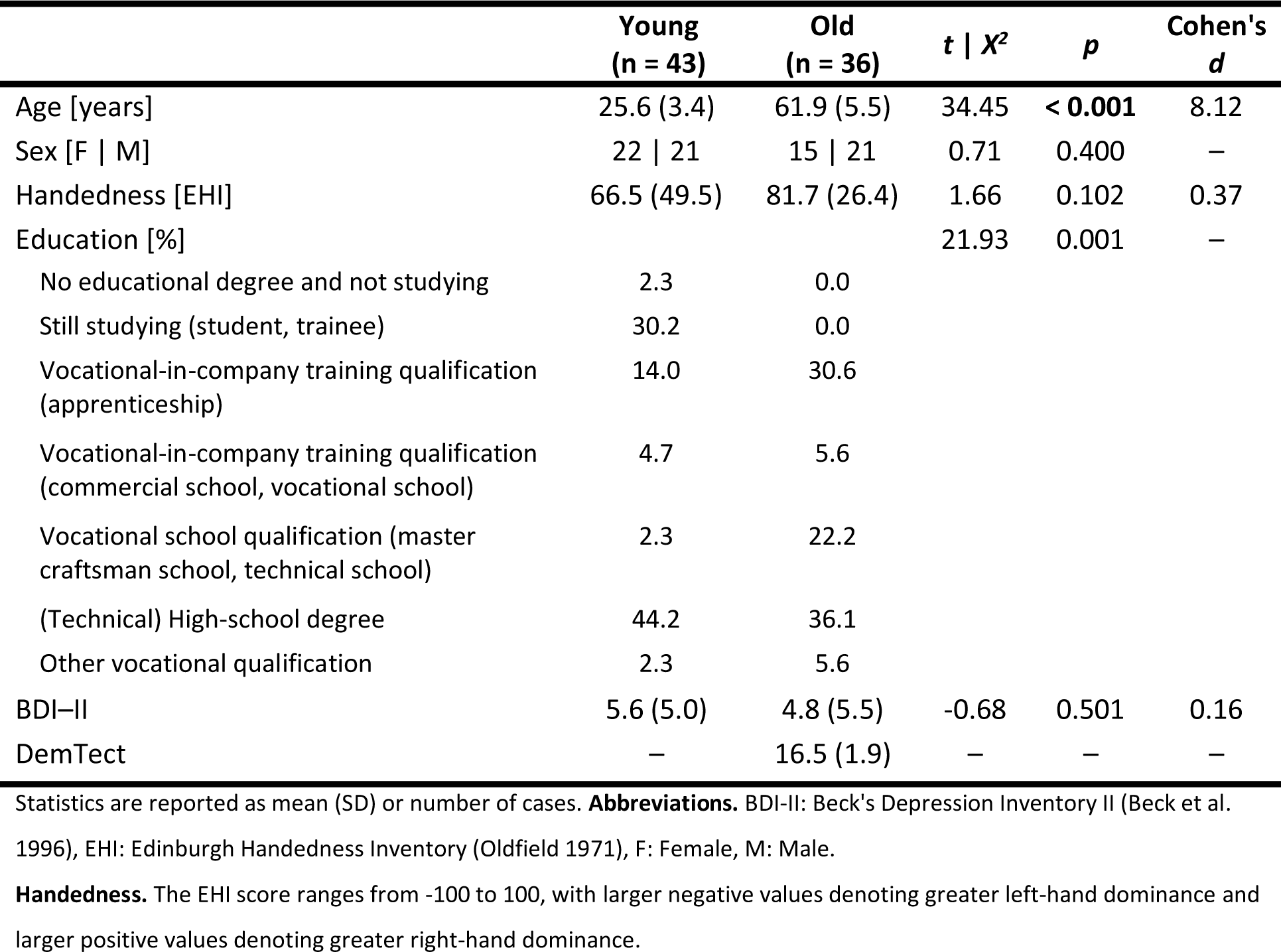
Sociodemographic information for the two subsamples.

### Psychometric assessment

Participants completed computerized cognitive tasks assessing other facets of dual-tasking outside the scanner. They sat in a quiet room in front of a 17.3-inch laptop (HP ProBook 470 G4; temporal resolution: 60 Hz; spatial resolution: 1920 × 1080 pixels) at a distance of about 50 cm, wearing over-ear headphones (Sennheiser HD 201). The covariates were collected as described below (see the **Supplementary Material** for a detailed description of each subtest and **Supplementary Figure S1** for an inter-correlation matrix between all behavioral scores across age groups and per age group).

1. *Crossmodal attention:* Crossmodal attention was assessed via the selective and focused attention subtests (WAF-S and WAF-F, respectively) from the Perception and attention functions battery from the Schuhfried Vienna Test System (https://marketplace.schuhfried.com/en/waf; visuo-auditory crossmodal test form S1). Since we aimed to assess both facets of attention jointly, we computed a compound score termed *crossmodal attention* by averaging mean reaction time in both tests per participant. Thus, higher scores reflect lower performance.
2. *Working memory*: Visuo-spatial working memory span was assessed via the computerized version of the Corsi block-tapping task (forward and backward versions) from the Schuhfried Vienna Test System (https://marketplace.schuhfried.com/en/corsi; test forms S1 and S5). We derived a compound score by averaging the number of correctly tapped sequences in both tasks; thus, higher values reflect higher performance.
3. *Global task-switching costs:* An alternative-runs task-switching paradigm (Stoet 2010; https://www.psytoolkit.org/experiment-library/taskswitching.html) was used to specifically assess global task-switching costs (also termed *mixing costs*). They reflect increased processing demands associated with maintaining and juggling two task configurations in mixed blocks, as compared to pure blocks (Rogers and Monsell 1995; Wylie and Allport 2000). We obtained global task-switching costs by subtracting mean reaction time (RT) in single-task trials from that in task-repeat trials, and thus, higher values or costs reflect lower levels of performance.

### Experimental protocol

We recorded brain activity while participants completed a single-stimulus onset dual-task paradigm (Weller et al. 2022; Paas Oliveros et al. 2023) during the second session of the study. In the task, participants were presented with high- or low-pitched pure sine tones (1000 or 500 Hz), to which they were instructed to respond by pressing one of two vertically arranged response keys with their index finger (high-positioned key) or thumb (low-positioned key; see **Figure 1**). Participants were asked to respond as fast and accurately as possible. In single-response blocks, responses were given with one hand (left or right), while in dual-response blocks, responses were given with both hands concurrently. Across blocks of trials, we manipulated the response mapping by varying S-R compatibility: Responses were to be executed towards either the same or opposite spatial location as implied by the pitch, inducing compatible (e.g., low pitch – low-positioned response key) or incompatible (e.g., low pitch – high-positioned response key) S-R mappings (Rusconi et al. 2006). This, in turn, allowed us to induce response-code conflicts in dual-response blocks: The response codes of either hand were either mutually congruent (R-R congruency: Both concurrent responses were either S-R compatible or S-R incompatible) or mutually incongruent (R-R incongruency: One hand’s response was S-R compatible, while the other hand’s one was S-R incompatible, e.g., combining a low pitch – low button compatible S-R pair with a concurrent low pitch – high button incompatible one). As a result, the paradigm included eight experimental conditions (see **Figure 1A**), but to reduce the complexity of the paradigm, each participant was presented with only seven experimental conditions (i.e., only one of the two R-R incongruent conditions, either condition 7 or 8, counterbalanced across the sample; see **Figure 1A**).

**Figure 1.**
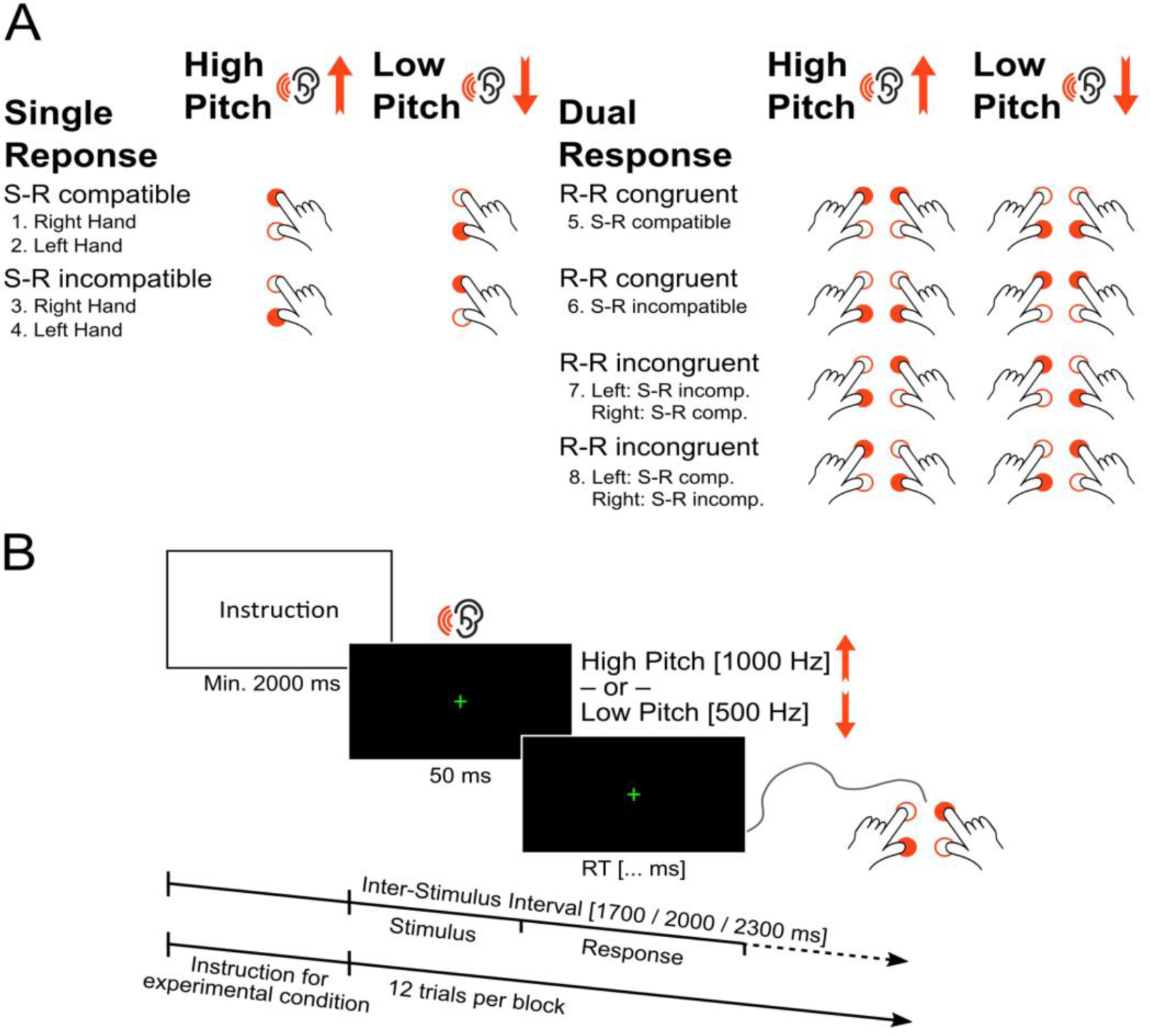
Spatial auditory–manual single-stimulus onset dual-task paradigm (Paas Oliveros et al. 2023). **A.** The eight experimental conditions of the dual-task paradigm according to the auditory stimulus, the number of responses, S-R compatibility, and R-R congruency. S-R compatibility was manipulated by having responses in the same or opposite spatial location implied by the pitch, leading to either compatible (e.g., low pitch – low response) or incompatible (e.g., low pitch – high response) S-R mappings. In dual-response conditions, responses were either mutually R-R congruent (i.e., both responses were S-R compatible or S-R incompatible) or incongruent (response-code conflict, e.g., low pitch – low response S-R compatible mapping combined with a concurrent low pitch – high response S-R incompatible one). **B.** Trial structure for an experimental condition. **Abbreviations.** S-R: Stimulus– response, R-R: Response–response.

Each experimental block started with a visual instruction of the hand(s) to be used and the S-R mapping(s) to be applied according to the specific experimental condition. Thus, experimental conditions were varied between blocks and were presented in pseudo-randomized order to control for potential task-sequence effects. Participants were instructed to maintain fixation on the centrally presented cross throughout the experimental blocks separated by resting breaks (13.5 or 14.0 s, pseudo-randomly varied across blocks). The visual instruction was displayed for a minimum of 2 s and until terminated by the participant via button press, followed by a post-instruction time interval of 1.0 or 1.5 s (varying pseudo-randomly between blocks). Each block comprised 12 trials (stimulus presentation: 50 ms, mean inter-stimulus interval: 2000 ms, pseudo-randomly varying between 1700, 2000, and 2300 ms, see **Figure 1B**). For familiarization, participants performed three blocks (cf. conditions 1, 2, and 5 in **Figure 1A**) outside the scanner. Inside the scanner, participants completed four blocks of each experimental condition, giving a total of 28 blocks with 48 trials overall per experimental condition for statistical analysis (see **Figure 1**).

Auditory stimuli were presented via single-use foam earplugs connected to a pneumatic audio system from allMRI GmbH (Nordheim, Baden-Württemberg, Germany, www.allmri.com). Participants executed motor responses using an MRI-compatible LUMItouch response box (Photon Control Inc., Burnaby, BC, Canada, www.photoncontrol.com), recorded via the Celeritas Fiber Optic Response System Operator (Psychology Software Tools, Sharpsburg, PA, www.pstnet.com). The instructions and the fixation cross were displayed on a 30” Apple Cinema HD screen (76.2 cm [diagonal], resolution: 1,280 pixels × 768 pixels, frame rate: 60 Hz). The display was located 2.55 m behind the scanner’s isocenter and was viewed via a mirror installed on the head coil. The experiment was controlled with Presentation® (Version 14.1, Neurobehavioral Systems, Inc., Berkeley, CA, www.neurobs.com) running under Microsoft Windows 7®.

### MRI data acquisition

Whole-brain images were acquired using a 3T Siemens MAGNETOM® Tim-TRIO scanner (Siemens Medical Systems, Erlangen, Germany) and a 12-channel head matrix coil. Fieldmap data were obtained using a B0 T2*-weighted gradient-echo sequence (TR: 34 ms, TE: 2.30/4.88 ms, FOV: 200 mm, voxel size: 3.1 × 3.1 × 3.1 mm^3^) before the functional sequence to correct the functional images for magnetic field inhomogeneity during preprocessing. Functional images were acquired using a T2*-weighted gradient-echo echo-planar imaging (EPI) sequence (TR: 2200 ms, TE: 30 ms, flip angle: 80°, FOV: 200 mm, voxel size: 3.1 × 3.1 × 3.1 mm^3^, in-plane matrix size: 64 × 64). Each volume had 36 axial slices (ascending series and interleaved multi-slice mode). Overall, 450 – 650 volumes were acquired per participant, depending on individual task performance speed.

### Data analysis

#### Behavioral data

In our single-stimulus onset dual-task paradigm, responses divergent to the instructed condition were considered error trials, as well as anticipatory responses (RT < 150 ms). Overly quick or slow correct responses were identified by RTs more than three times the standard deviation below or above the individual mean RT of the respective condition and were replaced with the given cut-off value. After that, the final RT averages for each experimental condition were calculated. For R-R congruent conditions (requiring either two S-R compatible or two incompatible responses; cf. conditions 5 and 6 in **Figure 1A**), we assumed a single joint task representation (Fagot and Pashler 1992; Paas Oliveros et al. 2023); thus, RT was averaged across both responses. In contrast, R-R incongruent conditions always contained both a compatible and an incompatible response (e.g., low pitch - low response S-R mapping combined with a concurrent low pitch – high response key), which were analyzed separately. Finally, dual-task speed costs were obtained by computing the difference in mean RT between dual- and single-task conditions of the same S-R compatibility level.

Error rate (ER) was calculated for each condition by adding the amount of omission and commission errors and dividing the sum by the number of total trials per condition. As mentioned above, participants with ER > 50% in at least one condition were excluded from further analysis. Dual-task accuracy costs were computed by subtracting the ER of single-task conditions from analogous dual-task conditions. For R-R incongruent trials, the ERs of the two related single-task conditions (i.e., S-R compatible and S-R incompatible) were subtracted separately, yielding dual-task accuracy costs for either S-R compatibility level under conditions of R-R incongruency.

We aimed to assess the effects of our experimental manipulations on dual-task costs with a measure that captures both performance facets, speed and accuracy, at the same time and accounts for possible speed–accuracy trade-offs. To this end, we implemented the recently proposed balanced integration score (BIS; Liesefeld et al. 2015; Liesefeld and Janczyk 2019), which is calculated by subtracting the standardized mean RT from the corresponding standardized mean proportion of correct responses (see the **Supplementary Material** for details on the BIS calculation). Higher absolute BIS values represent faster and more accurate performance. Analogous to RT, we derived dual-task performance costs by subtracting single-from dual-task BIS values according to the same S-R compatibility level.

The statistical analyses of the behavioral data focused on three dependent variables: Dual-task costs on BIS, RT, and ER. Each dependent variable was submitted to a three-way 2 × 2 × 2 mixed ANOVA. The factorial model included age group (young vs. older adults) as between-subject factor and S-R compatibility (compatible vs. incompatible) and R-R congruency (congruent vs. incongruent) as within-subject factors. Analyses were performed with SPSS 24.0 (IBM Corporation, Armonk, NY) and R version 3.6.1 (RStudio, Inc., Boston, MA), and statistical significance level was set to *p* < 0.05.

#### Functional MRI preprocessing

The functional MRI images were preprocessed with an in-house pipeline using SPM12 (Statistical Parametric Mapping, version 12, Wellcome Department of Imaging Neuroscience, London, UK, https://www.fil.ion.ucl.ac.uk/spm/) as implemented in MATLAB, version 9.4.0 (R2018a, The MathWorks Inc., Natick, MA) and FSL (http://fsl.fmrib.ox.ac.uk/fsl/fslwiki/FSL).

For each participant, four dummy functional volumes (EPI) preceded the actual acquisition to allow for magnetic field saturation and were discarded prior to further analyses (McRobbie et al. 2006). Next, using SPM12, motion correction was done via an affine two-pass registration that spatially realigned all EPI volumes to the first and subsequently to the mean image, followed by distortion correction (“unwarping”) using the voxel-displacement map obtained from the magnitude and phase differences recorded in the fieldmap. Then, slice-time correction was done using FSL. The mean EPI image of each subject was used for spatial normalization to MNI space using the *unified segmentation* approach (Ashburner and Friston 2005). The resulting deformation parameters were then applied to all other EPI volumes. Finally, resampling to a voxel size of 2 × 2 × 2 mm^3^ (seventh-degree B-spline interpolation) and spatial smoothing with an isotropic 8-mm full-width-at-half-maximum Gaussian kernel were done.

#### Task-related brain activity

All statistical analyses were conducted within a mass-univariate framework where the evoked hemodynamic response at each voxel was independently estimated using a generalized linear model (GLM). At the single-subject level, regressors for an event-related model of experimental effects were defined according to the three experimental factors number of responses (single vs. dual), S-R compatibility (SRC vs. SRI), and R-R congruency (in dual-response conditions, RRC vs. RRI). For each experimental condition, we set up separate regressors for trials with high vs. low response locations, respectively. This resulted in 10 task regressors: Single_SRC_ low (1) or high (2) response location; Single_SRI_ low (3) or high (4); Dual_SRC-RRC_ low (5) or high (6); Dual_SRI-RRC_ low (7) or high (8); and Dual_RRI_ low (9) or high (10). For R-R incongruent trials, this division was based on the S-R compatible response. All task regressors were accompanied by a parametric modulator for the hand(s) used for responding (left = −1, right = 1, or both = 0). We implemented a varying-epoch model, in which events were locked to the tone onset, and their duration was defined by the reaction time (second response in dual trials). The resulting epoch functions and their parametric modulations were convolved with a canonical hemodynamic response function (HRF) and its temporal and dispersion derivatives to account for variability in onset time and width of the BOLD signal change, respectively. We included further regressors of no-interest for error trials (fixed duration of 0.7 s), the times during which the instruction for each block was presented, and the six head-movement parameters estimated during the rigid-body spatial realignment (i.e., translation and rotation relative to the X, Y, Z axes). After each voxel’s time series was high-pass filtered at a cut-off period of 128 s to remove low-frequency signal drifts and corrected for dependent observations according to an autoregressive first-order correlation structure, we calculated parameter estimates of the HRF regressors from the least-mean-squares fit of the model to the time series.

Random-effects analysis was used to examine the contrasts between the single- and dual-response conditions and their interaction with age at the group level. To this end, a sum contrast for each of the five experimental conditions of interest (Single_SRC_, Single_SRI_, Dual_SRC-RRC_, Dual_SRI-RRC_, Dual_RRI_) was created for every subject. For example, Single_SRC_ was created by the sum contrast combining the two Single_SRC_ regressors (low and high tones). Next, the five regressors were entered into a group-level model with two age groups, in which we assumed unequal variance between subjects. This resulted in a second-level design matrix with ten regressors. We were interested in the increase and decrease in brain activity related to the following contrasts: (1) Dual R-R congruent vs. single-task trials (dual-response execution), (2) dual R-R incongruent vs. single-task trials (dual-tasking), and (3) dual R-R incongruent vs. dual R-R congruent trials (dual-task response-code conflict). These contrasts were assessed via strict minimum conjunctions with the corresponding main effect of interest to restrict differences to those voxels that also showed a significant activation or deactivation relative to the implicit baseline. Thus, for example, we computed [Dual_RRC_ > Single] ∩ [Dual_RRC_ +] as well as [Dual_RRC_ < Single] ∩ [Dual_RRC_ –] across both age groups to assess increased as well as decreased brain activity during dual-response execution. Because a major interest of this study lay in examining brain activity linked to processing R-R incongruency (i.e., dual-task conflict), we defined a task mask based on the conjunction assessing this process (i.e., [Dual_RRI_ > Dual_RRC_] ∩ [Dual_RRI_ +]).

To analyze the age-related differences in brain activity associated with each of the three processes of interest (i.e., the contrasts mentioned above), we assessed the interaction of each contrast with age via strict minimum conjunctions with the corresponding contrast of each age group, separately. For example, we computed [YA: Dual_RRC_ > Single, OA: Dual_RRC_ < Single] ∩ [YA: Dual_RRC_ > Single] to assess increased activity among young (vs. older) adults during dual-response execution, and [YA: Dual_RRC_ < Single, OA: Dual_RRC_ > Single] ∩ [OA: Dual_RRC_ > Single] for older (vs. young) adults’ increased activity. All activations are reported at a cluster-level family-wise error (cFWE)-corrected threshold of *p* < 0.05 (cluster-forming threshold at voxel level: *p* < 0.001, *k* ≥ 224 voxels).

All results were anatomically labeled by reference to probabilistic cytoarchitectonic maps of the human brain using the SPM Anatomy Toolbox Version 3 (Eickhoff et al. 2005; Eickhoff et al. 2007, https://www.fz-juelich.de/en/inm/inm-7/resources/jubrain-anatomy-toolbox/jubrain-toolbox) and visualized with Surf Ice (https://www.nitrc.org/projects/surfice/, version 1.0.20211006) and MRIcroGL (https://www.nitrc.org/projects/mricrogl, version 1.2.20211007).

#### Covariance analysis

We computed four separate group-level models to investigate whether and how brain activity evoked during dual-task response-code conflict is linked to dual-task performance and other cognitive abilities related to dual-tasking and how this may differ by age. Each of the four models was identical to the GLM described above but included a behavioral score of interest as a covariate: (1) Dual-task performance (mean absolute BIS score in the S-R compatible, R-R incongruent condition), (2) crossmodal attention, (3) working memory, and (4) global task-switching costs. We focused on the S-R compatible response in R-R incongruent dual-task conditions because compatible responses suffered the highest dual-task costs and crosstalk effects with this sample and in a previous study (Paas Oliveros et al. 2022). As opposed to dual-task performance, the last three behavioral scores were obtained through a psychometric assessment outside the scanner. Crossmodal attention, working memory, and task switching represent different cognitive abilities putatively involved in efficient dual-tasking. Thus, aiming to explain the neural results more comprehensively, we assessed the variance shared with those cognitive abilities. The design matrix for each of the four covariance analyses included ten task and ten covariance regressors. We analyzed the positive and negative linear correlations between the covariate of interest and R-R incongruency-related brain activity (Dual_RRI_ main effect of the covariance regressors) as well as their age differences. Second-level statistics were only assessed for voxels surviving the contrast Dual_RRI_ > Dual_RRC_ (inclusive mask at *p* < 0.001). Since, in these analyses, we were interested in the covariance of any brain activity globally linked to dual-task response-code conflict, we used a slightly less restrictive mask than the task mask described above.

#### Generalized psychophysiological interaction (gPPI) analysis

To investigate whether task-relevant brain areas change their connectivity to other brain areas depending on dual-response conditions, we employed generalized psychophysiological interaction (gPPI) analyses using the gPPI toolbox, version 13.1 (https://www.nitrc.org/projects/gppi) in SPM12. PPI models measure task-modulated synchronization between a predefined seed region and one or more brain areas by exploring the physiological response (HRF-convolved BOLD signal) in one brain region in terms of the context-dependent response of another region (O’Reilly et al. 2012). We implemented gPPI because it extends the standard SPM PPI approach to experimental designs with more than two conditions (McLaren et al. 2012). More specifically, we defined two seed regions by extracting the peak coordinates that showed the strongest brain activation difference during dual-task response-code conflict, using the conjunction [Dual_RRI_ > Dual_RRC_] ∩ [Dual_RRI_ +] across both age groups. A 6-mm sphere was created around each peak, and a gPPI analysis was computed for each region separately.

The BOLD signal time series for each seed region was deconvolved to model the time series of the neuronal signal. These time series were entered into the same first-level analysis described previously (see “Task-related brain activity subsection”). The following regressors were computed for each participant: (1) The same ten “psychological” regressors of the model testing for task-related brain activity, including hand as a parametric modulator, i.e., stick functions at trial onsets convolved with the canonical HRF; (2) a “physiological” regressor formed by the first eigenvariate of the seed region time series; (3) the PPI regressors for each experimental condition created by multiplying (interaction) the deconvolved BOLD signal of the seed region with the condition onsets and convolving the product with the canonical HRF (McLaren et al. 2012); (4) the regressors of no interest formed by the error trials (fixed duration of 0.7 s), the epochs during which the instruction for each block was presented, and the six head motion parameters.

For each participant, the following contrasts of interest were created with the PPI regressors: (1) Single, (2) Dual_RRC_, (3) Dual_RRI_, (4) Dual_SRC-RRC_ > Single_SRC_, (5) Dual_SRI-RRC_ > Single_SRI_, and (6) Dual_RRI_ > Dual_RRC_. The last three contrasts were weighted by the number of trials in each condition. We then computed three separate group-level models for either seed region by passing the estimates of the (1) Dual_RRI_ > Dual_RRC_, (2) Dual_RRC_, or (3) Dual_RRI_ PPI contrasts, respectively, and we inspected the main positive and negative effects of each PPI contrast through a conjunction across age groups (e.g., [Young: Dual_RRI_ > Dual_RRC_] ∩ [Old: Dual_RRI_ > Dual_RRC_]). Second-level statistics were only assessed at voxels contained within the task mask (i.e., [Dual_RRI_ > Dual_RRC_] ∩ [Dual_RRI_ +], inclusively masked at *p* < 0.001). We first inspected connectivity for each seed during response-code conflict (i.e., Dual_RRI_ > Dual_RRC_ PPI contrast for each seed). Next, since we were interested in inspecting the connectivity commonalities in both dual R-R congruent and incongruent conditions, for each seed, we combined the main-effect results from the Dual_RRC_ (i.e., [Young: Dual_RRC_] ∩ [Old: Dual_RRC_]) and Dual_RRI_ (i.e., [Young: Dual_RRI_] ∩ [Old: Dual_RRI_]) PPI regressors derived from separate GLMs using the Image Calculation minimal conjunction function in SPM (ImCalc, http://tools.robjellis.net; Expression: min(Dual_RRC_, Dual_RRI_)). Additionally, we computed the differences between the two age groups for each model to analyze age-related hyper- and hypoconnectivity patterns in each condition.

## Results

### Behavioral data

**Figure 2** and **Table 2** show the dual-task costs on the BIS as well as dual-task costs separately for response speed and accuracy (see **Supplementary Table S1** for the mean absolute single- and dual-task performance score for each age group).

**Figure 2.**
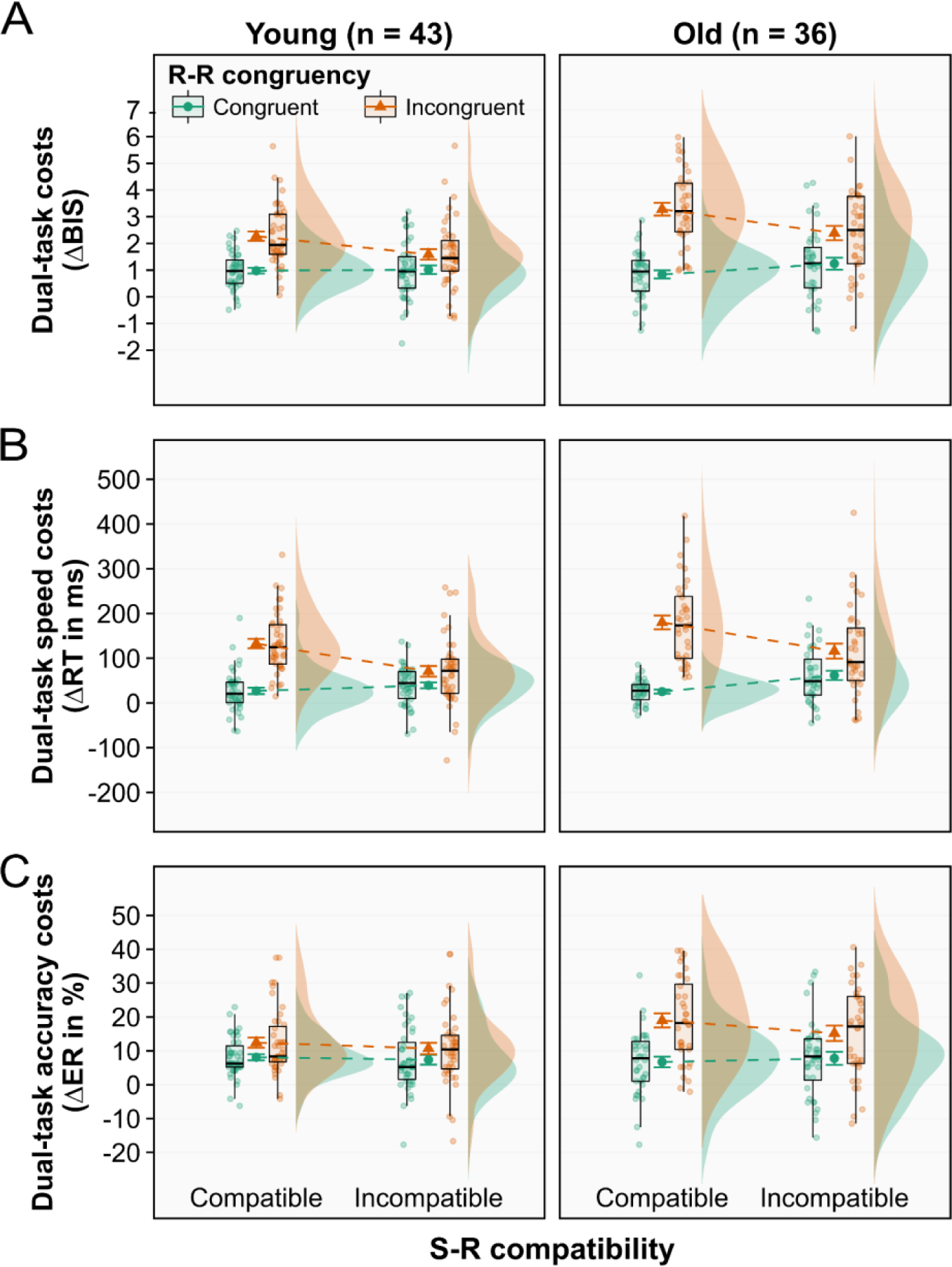
Dual-task costs on **(A)** the balanced integration score, **(B)** reaction time, and **(C)** error rate according to age group, S-R compatibility, and R-R congruency. Dual-task costs were obtained through the difference in mean RT between analogous dual- and single-task conditions. Error bars represent S.E.M. **Abbreviations.** BIS: Balanced integration score, ER: Error rate, RT: Reaction time, S-R: Stimulus–response, R-R: Response–response.

**Table 2.** Descriptive statistics of the behavioral data for each age group.

| Behavioral Score | Young (n = 43) |  | Old (n = 36) |  |
| --- | --- | --- | --- | --- |
|  | *M* | *SD* | *M* | *SD* |
| *Dual-task costs (ΔBIS)* |  |  |  |  |
| SRC-RRC | 0.98 | 0.69 | 0.84 | 0.91 |
| SRI-RRC | 1.01 | 1.06 | 1.24 | 1.34 |
| SRC-RRI | 2.27 | 1.14 | 3.28 | 1.42 |
| SRI-RRI | 1.59 | 1.29 | 2.39 | 1.61 |
| *Dual-task costs on reaction time (ms)* |  |  |  |  |
| SRC-RRC | 26.85 | 46.08 | 25.39 | 26.28 |
| SRI-RRC | 39.27 | 43.90 | 61.68 | 62.29 |
| SRC-RRI | 132.61 | 68.07 | 180.13 | 92.16 |
| SRI-RRI | 70.73 | 77.91 | 116.02 | 100.82 |
| *Dual-task costs on error rate (%)* |  |  |  |  |
| SRC-RRC | 8.12 | 5.93 | 6.71 | 9.52 |
| SRI-RRC | 7.36 | 9.36 | 7.78 | 11.58 |
| SRC-RRI | 12.43 | 9.80 | 18.98 | 12.50 |
| SRI-RRI | 10.71 | 11.04 | 15.19 | 13.51 |
| Crossmodal attention [ms]<sup>a</sup> | 356.77 | 53.95 | 408.89 | 86.12 |
| Working memory [#]<sup>b</sup> | 11.34 | 2.43 | 8.32 | 1.87 |
| Global task-switching costs [ms]<sup>c</sup> | 113.06 | 162.42 | 222.71 | 214.47 |

**Abbreviations.** BIS: Balanced integration score, RRC: Response–response congruent, RRI: Response–response incongruent, SRC: Stimulus–response compatible, SRI: Stimulus–response incompatible.
<sup>a</sup> Measured via the compound speed score derived from the crossmodal (visuo-auditory) selective and focused attention tests of the Vienna Test System.
<sup>b</sup> Measured via the compound score of mean number of correctly tapped sequences in the forward and backward versions of the Corsi block-tapping test of the Vienna Test System.
<sup>c</sup> Measured via the mean reaction time difference between mixed-task repeat trials and single-task trials.

For the dual-task costs on the BIS (see **Figure 2A** and **Supplementary Table S2** for the detailed statistics), the mixed three-way ANOVA revealed a significant effect of age, *F*(1,77) = 6.46, *p* = 0.013, 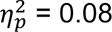, indicating higher dual-task costs in older adults (*M* ± *SD*: 1.93 ± 1.64) than in young ones (1.46 ± 1.18). There were also significant main effects of S-R compatibility, *F*(1,77) = 6.65, *p* = 0.012, 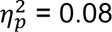, and R-R congruency, *F*(1,77) = 84.29, *p* < 0.001, 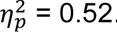. However, these main effects were qualified by significant interactions between S-R compatibility and R-R congruency, *F*(1,77) = 52.01, *p* < 0.001, 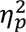 = 0.40, and between R-R congruency and age, *F*(1,77) = 8.37, *p* = 0.005, 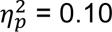. In particular, the former interaction revealed that the S-R compatibility effect strongly depended on the R-R congruency level: When the response codes for both hands were *congruent* with each other (R-R congruent trials), dual-task costs did not differ between S-R compatible and incompatible responses in a pairwise comparison (*p* = 0.13). In contrast, when both response codes were *incongruent* with each other, dual-task costs were significantly higher for S-R compatible responses (mean BIS costs: 2.78 ± 1.28) than for incompatible ones (1.99 ± 1.45, *p* < 0.001). Furthermore, the interaction between R-R congruency and age revealed that in R-R incongruent trials, dual-task costs were significantly higher for older adults (2.83 ± 1.57) than for younger ones (1.93 ± 1.26, *p* = 0.002), but this was not the case for R-R congruent trials (*p* = 0.81). While we did not find a significant two-way interaction between S-R compatibility and age, *F*(1,77) = 0.13, *p* = 0.72, 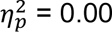, we observed a three-way interaction between S-R compatibility, R-R congruency and age, *F*(1,77) = 4.31, *p* = 0.041, 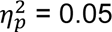, indicating that the S-R compatibility × R-R congruency interaction described above significantly differed between young and older adults. In particular, it revealed that the larger difference in dual-task BIS costs between S-R compatible and incompatible trials that comes with R-R incongruency was accentuated in older (vs. young) adults.

Dual-task speed and accuracy costs (see **Figure 2B-C** and **Table 2**) followed the same pattern of results (see **Supplementary Table S2** for the detailed ANOVA statistics). However, when analyzed separately, there was no three-way interaction (*p* = 0.196 and *p* = 0.095, respectively). Additionally, the main effects of age and S-R compatibility were not significant for ER (*p* = 0.125 and *p* = 0.171, respectively). Nevertheless, the overall consistency in the result patterns supports the validity of the behavioral effects and suggests that the BIS could be an informative and sensitive alternative for assessing dual-task costs, accounting well for inter-individual differences in speed–accuracy trade-offs.

To assess whether the observed age effects on dual-task performance reflect detrimental domain-specific processes or could be explained by an age-related generalized slowing (Salthouse 1996), we implemented the Brinley correction (Brinley 1965; Cerella 1994) for RT and re-assessed the age-corrected three-way ANOVA on dual-task speed costs (see **Supplementary Material** for details on the statistical analysis). The age differences in dual-task speed costs in R-R incongruent trials turned insignificant after the correction.

We additionally tested whether the observed effects on dual-task speed costs persisted after accounting for response grouping in dual-response trials (Pashler 1994; Miller and Ulrich 2008). Crucially, after removing strongly synchronized responses (see **Supplementary Material** for details on the statistical analysis), the dual-task cost asymmetry for S-R compatibility under R-R incongruency remained significant; however, the age main effect and the age × R-R congruency interaction did not. We conclude that beyond response grouping, participants engaged another mechanism to cope with crosstalk under response-code conflict conditions (see Paas Oliveros et al. 2022). On the other hand, considering the three-way interaction effect on dual-task BIS costs but the disappearance of age effects when accounting for generalized slowing and response grouping on dual-task speed costs, the results remain inconclusive as to whether the age-related differences observed do result from process-specific difficulties with solving response-code conflicts in advanced age or can (largely) be explained by generalized slowing or less efficient response grouping in the elderly.

In summary, replicating our earlier findings (Paas Oliveros et al. 2023), we found that across age groups, both responses suffered more costs in dual-task conditions when being based on mutually incongruent (vs. congruent) response codes; however, the detrimental impact of R-R incongruency was disproportionately greater on S-R compatible responses than on S-R incompatible ones, even after removing highly synchronized responses. Furthermore, the interaction between R-R congruency and age revealed that the detrimental effect of incongruent response codes was exacerbated in older (vs. young) adults, pointing towards deficits in multiple-action control, partially explained by generalized slowing and response grouping. Following these observations, we examined the association of dual-tasking and response-related interference with brain activity and connectivity, as well as their age-related differences.

### Neuroimaging data

#### Task-related brain activity

Brain activity related to dual-response execution, as compared to single-hand responding, was investigated via two separate conjunctions: [Dual_RRC_ > Single] ∩ [Dual_RRC_ +] and [Dual_RRC_ < Single] ∩ [Dual_RRC_ –]. These analyses revealed stronger activations in a large set of regions (see **Figure 3A** and **Supplementary Table S5**), including bilateral primary motor (M1) and dorsal premotor cortex (dPMC), supplementary motor area (SMA), somatosensory areas, superior parietal lobe (SPL), cerebellum, thalamus, and putamen as well as right inferior frontal gyrus (IFG) and central insula. Conversely, reduced activity was identified mainly in bilateral occipital areas, medial frontal pole, anterior cingulate cortex (ACC), and right hippocampus.

**Figure 3.**
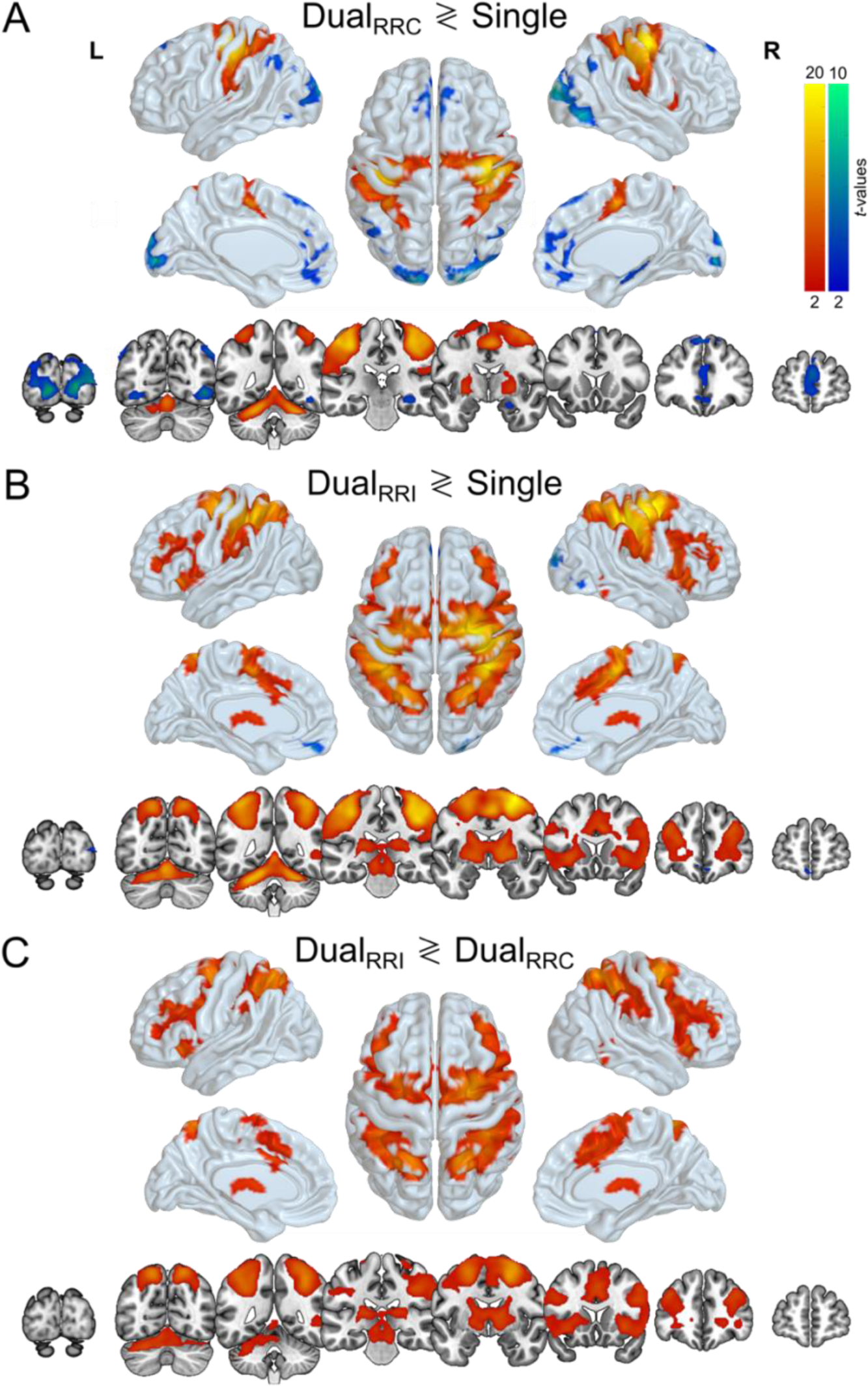
Task-specific brain activity. **(A)** Dual-response execution network obtained by contrasting dual R-R congruent vs. single-task conditions (Dual_RRC_ ≷ Single). **(B)** Dual-task network obtained by contrasting dual R-R incongruent vs. single-task conditions (Dual_RRI_ ≷ Single). **(C)** Dual-task response-code conflict network computed by contrasting dual-task R-R incongruent vs. dual R-R congruent conditions (Dual_RRI_ ≷ Dual_RRC_). Hot colors represent activations (i.e., “greater than” contrasts), and cool colors represent deactivations (i.e., “smaller than” contrasts). All activations were significant at cluster-level *p* < 0.05 (cFWE-corrected) with a cluster-forming threshold of *p* < 0.001 at voxel level. **Abbreviations.** RRC: Response–response congruent, RRI: Response–response incongruent.

Next, we tested for brain activity associated with R-R incongruent dual-tasking at large via the conjunctions [Dual_RRI_ > Single] ∩ [Dual_RRI_ +] and [Dual_RRI_ < Single] ∩ [Dual_RRI_ –], respectively. We found increased activations in an extensive fronto-parieto-insular network during dual-tasking (see **Figure 3B** and **Supplementary Table S5**), including bilateral motor and premotor areas (M1, dPMC, SMA, preSMA), somatosensory areas, dorsolateral prefrontal cortex (dlPFC), superior frontal gyrus (SFG), midcingulate cortex (MCC), SPL, supramarginal gyrus, insular cortex, thalamus, and cerebellum, as well as a cluster in right inferior temporal gyrus. Decreased activity was identified in right occipital areas, bilateral medial frontal pole, and paracingulate gyrus.

Brain activity specifically associated with response-code conflict in dual-tasking was again tested via two separate conjunctions: [Dual_RRI_ > Dual_RRC_] ∩ [Dual_RRI_ +] and [Dual_RRI_ < Dual_RRC_] ∩ [Dual_RRI_ –]. These analyses demonstrated that a large fronto-parieto-insular network also was more strongly activated during R-R incongruent than congruent dual-tasking (see **Figure 3C** and **Supplementary Table S5**). Still, this network was more restricted than the previously observed one more generally related to (incongruent) dual-tasking relative to single-tasking (cf. **Figure 3B**), especially sparing primary motor and somatosensory areas. The network covered bilateral premotor areas (dPMC, SMA, preSMA), dlPFC, SFG, MCC, SPL, supramarginal gyrus, insular cortex, thalamus, and left cerebellum. Here, no significant deactivations were identified. For details on age group-specific increases and decreases in brain activity for each contrast of interest (i.e., dual-response execution, dual-tasking at large, and dual-task response-code conflict), the reader is referred to **Supplementary Figure S5** and **Table S6**.

Examining age-related differences did not yield a statistically significant interaction between type of response execution (dual congruent vs. single) and age. In contrast, when assessing dual-tasking at large (dual incongruent vs. single), we found a significant interaction with age (see **Figure 4A**), in which young participants showed stronger dual-task-related activity in right M1, SFG, SPL, and supramarginal gyrus, relative to older participants. In contrast, older (vs. young) adults showed higher dual-task-driven activation SMA and bilateral prefrontal areas (including ACC and paracingulate gyrus MFG, SFG and frontal pole) as well as left putamen and pallidum. Crucially, a significant interaction with age revealed greater activity in left superior frontal gyrus for older (vs. young) adults during dual-task response-code conflict (dual incongruent vs. congruent; see **Figure 4B**). In summary, dual-task response-code conflict was associated with increased brain activity in a large fronto-parieto-insular network, sparing primary motor and somatosensory areas engaged in dual (vs. single) conditions at large. Moreover, older (vs. young) adults showed hyperactivity in left superior frontal gyrus when facing spatially incongruent mapping rules to be applied concurrently.

**Figure 4.**
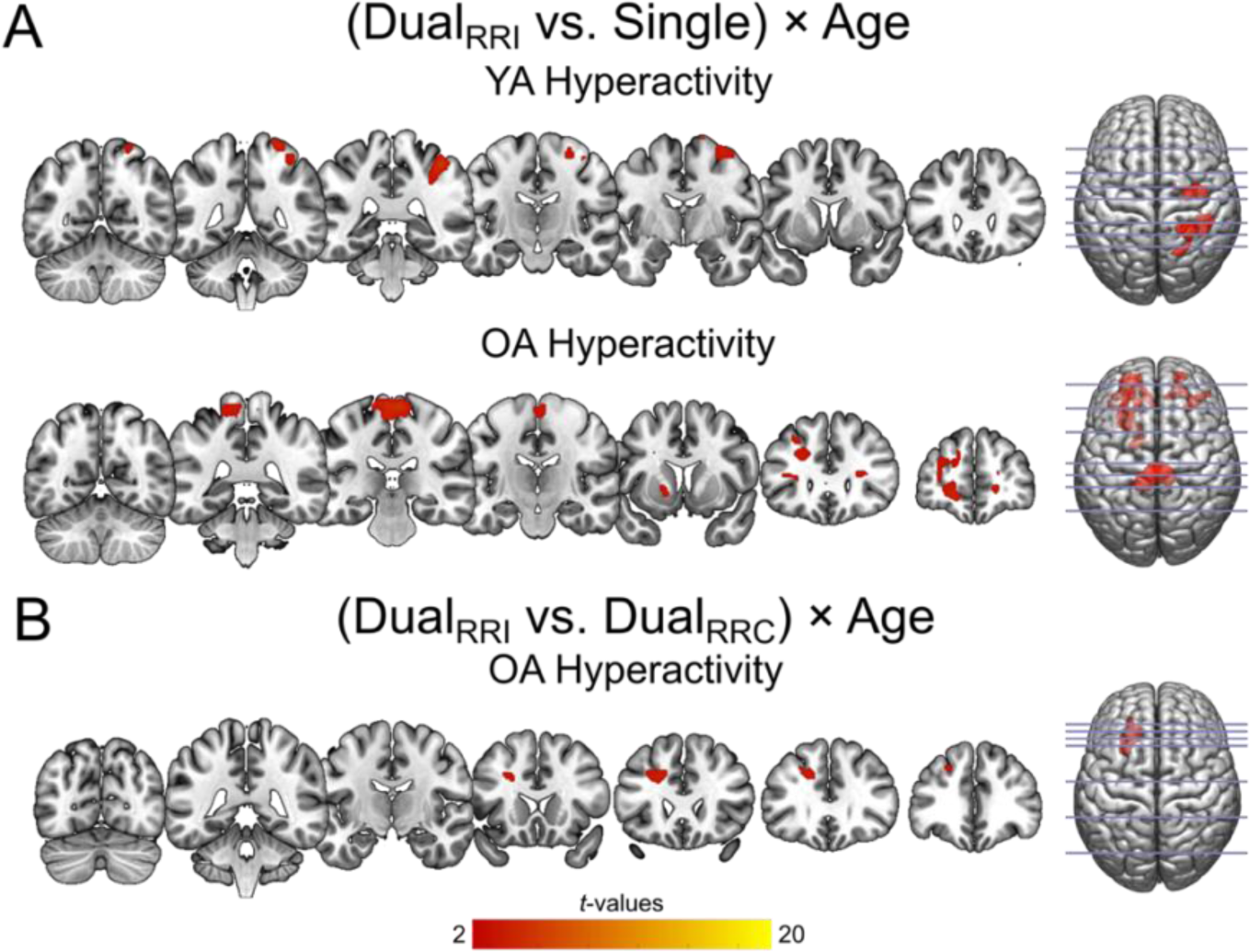
Statistical interaction between **(A)** the dual-task network and age, as well as **(B)** the dual-task response-code conflict network and age. There was no significant interaction between the dual-response execution network and age. Hot colors represent activations (i.e., “greater than” contrasts), and cool colors represent deactivations (i.e., “smaller than” contrasts). All activations were significant at cluster-level *p* < 0.05 (cFWE-corrected) with a cluster-forming threshold of *p* < 0.001 at voxel level. **Abbreviations.** OA: Older adults, RRC: Response– response congruent, RRI: Response–response incongruent, YA: Young adults.

#### Covariance analysis

With the next set of analyses, we aimed to better characterize dual-task conflict-related brain activity by assessing which brain regions shared variance with individual dual-task performance and related cognitive abilities (see **Supplementary Figure S1** for correlations with the behavioral scores). Across separate GLMs, one for each of the four covariates, we tested the positive and negative effects of the covariate regressors for the R-R incongruent condition (Dual_RRI_ × COV +/–, inclusively masked by Dual_RRI_ > Dual_RRC_).

First, we assessed individual dual-task performance through the BIS score, jointly reflecting speed and accuracy for the S-R compatible response in R-R incongruent trials (i.e., those responses where dual-task costs were highest; see **Figure 2**). Our results showed that better R-R incongruent dual-task performance (higher BIS) was linked to decreased activity in bilateral cuneus and lingual gyrus (Dual_RRI_ × BIS_Dual SRC-RRI_ –; see **Figure 5A**). The second model showed that with lower crossmodal attentional performance (i.e., longer RT), dual-task conflict-related brain activity decreased particularly in right thalamus and globus pallidus during R-R incongruency (Dual_RRI_ × Att –; see **Figure 5B**). Finally, in the third model, higher global task-switching costs were associated with decreased activity in bilateral SPL, intraparietal sulcus (IPS), and right cerebellum (Dual_RRI_ × gTSC –; see **Figure 5C**). Conflict-related brain activity was not significantly modulated by working memory performance, and we did not find any positive correlation with the covariates of interest. For more detailed results from the covariance analyses across age groups, please see **Supplementary Table S9**.

**Figure 5.**
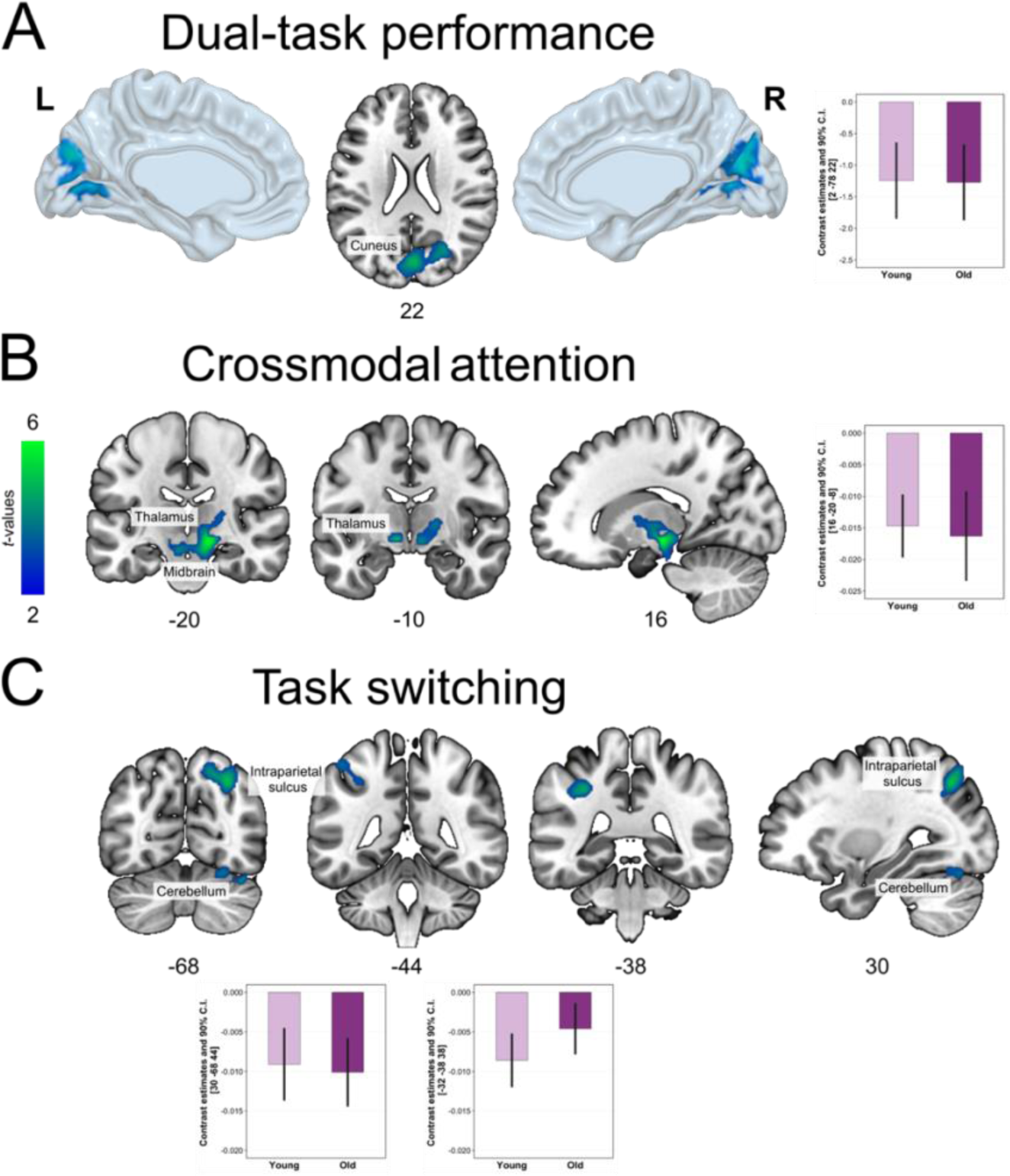
Brain activity in the conflict-related network showing a significant negative correlation with the covariates during R-R incongruency, assessed through separate models for each covariate. Reduced activity in the illustrated brain areas was associated with **(A)** better dual-task performance (i.e., higher balanced integration score [BIS] for the S-R compatible response in R-R incongruent trials), **(B)** lower crossmodal attentional performance (i.e., longer RT), and **(C)** lower task-switching performance (i.e., higher global task-switching costs) during R-R incongruent dual-tasking. All effects were significant at cluster-level *p* < 0.05 (cFWE-corrected) with a cluster-forming threshold of *p* < 0.001 at voxel level.

When investigating age differences in covariate effects, we only found significant results for crossmodal attention performance: In older (vs. young) adults, lower crossmodal attentional performance (i.e., longer RT) was more strongly associated with increased brain activity in left IPS, right fusiform gyrus, and cerebellum during R-R incongruency (Dual_RRI_ × Att: OA > YA, see **Figure 6**). No significant age differences were found in the modulation of conflict-related brain activity by dual-task performance, working memory, or global task-switching costs. For more detailed age-related results from the covariance analyses, please refer to **Supplementary Table S10**. To summarize, while motor and parietal areas of the conflict-related network were modulated by attentional and task-switching abilities, better conflict resolution in dual-tasking was linked to lower visual cortex activity. In addition, older (vs. young) adults showed higher left parietal and right cerebellar activity associated with lower attentional performance.

**Figure 6.**
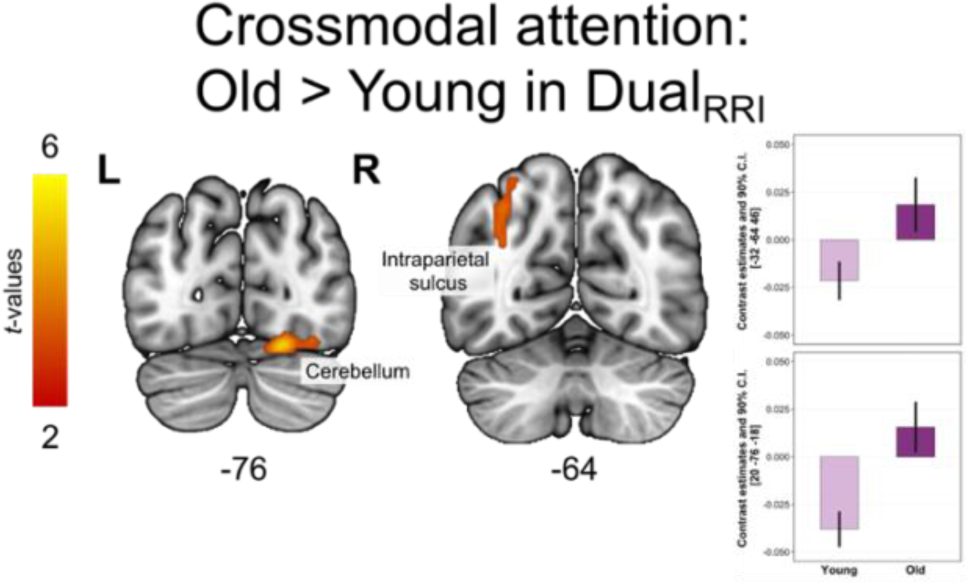
Brain activity in the conflict-related network showing significant age differences (Old > Young) in its correlation with crossmodal attention during R-R incongruent (RRI) dual-tasking. In older (vs. young) adults, lower crossmodal attentional performance (i.e., longer RT) was more strongly associated with increased brain activity in the illustrated brain areas during R-R incongruency. All differences were significant at cluster-level *p* < 0.05 (cFWE-corrected) with a cluster-forming threshold of *p* < 0.001 at voxel level.

#### gPPI analysis

For the gPPI analyses, we selected as seeds the two strongest activation peaks specifically associated with dual-task response-code conflict ([Dual_RRI_ > Dual_RRC_] ∩ [Dual_RRI_+]), which were located in right dPMC [24, −6, 54] and right SPL [20, −60, 62]. By means of the gPPI analyses, we aimed to identify brain areas of the dual-task conflict-sensitive network in which the connectivity with the seeds was most strongly altered during (1) dual-task response-code conflict, (2) dual R-R congruent, and (3) incongruent conditions. Hereby, we assessed commonalities in connectivity across both dual R-R congruent and incongruent conditions.

The gPPI analyses revealed no significant task-modulatory effect on connectivity of dual-task response-code conflict (Dual_RRI_ > Dual_RRC_ PPI regressor) for either seed region, nor age-related differences thereof. In contrast, we observed that during R-R congruent and incongruent conditions, connectivity increased between the right dPMC and the following areas of the task network: bilateral M1, preSMA, ACC, IFG, insula, SPL, IPS, and precuneus, as well as left MFG (see **Figure 7A**). Analogously, for the right SPL seed, we observed an increase in synchronization with bilateral IPS, precuneus, and angular gyrus, preSMA, ACC, thalamus, pallidum, and putamen, as well as right dPMC, frontal areas, and left cerebellum (see **Figure 7B**). There was no significant decrease in task-related connectivity for either seed. The results from both seeds overlapped in areas of bilateral parietal (SPL, IPS, precuneus, and supramarginal gyrus) and frontal (dPMC, SMA, ACC, and SFG) cortex, as well as the anterior insular cortex and right operculum (see **Figure 7C**). Nevertheless, it is noteworthy that visually comparing the results from both seeds, right dPMC tended to show increased connectivity to fronto-parieto-insular areas. In contrast, right SPL additionally increased its connectivity to subcortical regions and the cerebellum, parietal areas, and ipsilateral frontal regions. See **Supplementary Table S11** for detailed results on the connectivity commonalities in both dual R-R congruent and incongruent conditions for each seed across both age groups, and **Supplementary Table S12** for results for each condition, i.e., dual R-R congruent and incongruent conditions separately.

**Figure 7.**
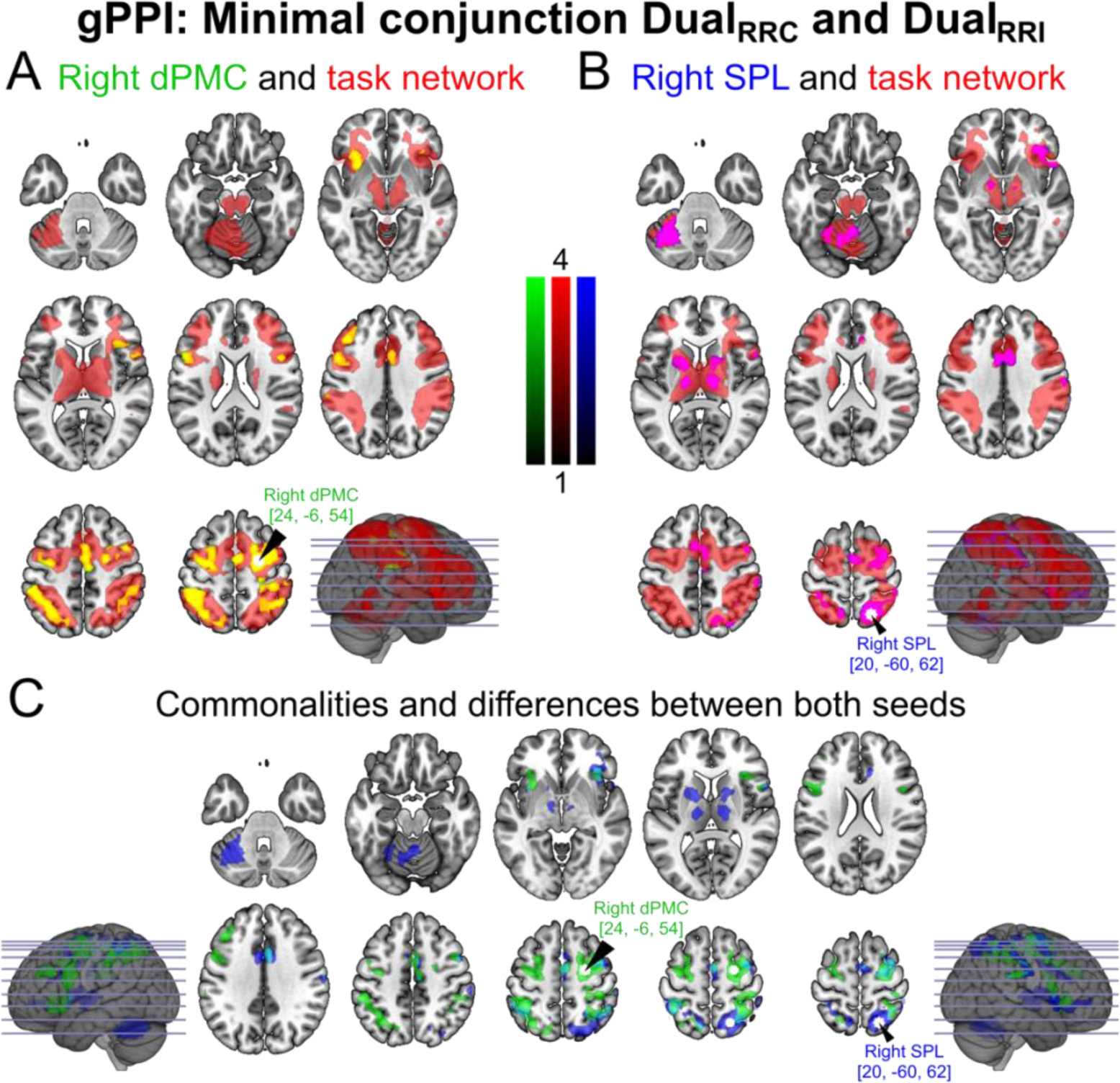
Results of the generalized PPI analysis testing for changes in the connectivity of right **(A)** dPMC and **(B)** SPL, each combining the connectivity results from the Dual_RRI_ (vs. baseline) and Dual_RRC_ (vs. baseline) PPI regressors. Each contrast was estimated through the conjunction of both age groups in separate models and masked with dual-task conflict-sensitive network (Dual_RRI_ + > Dual_RRC_; see **Figure 3C**). Effects in **(A)** and **(B)** are overlaid on the dual-task conflict-related network. **(C)** Overlay of the results across both gPPI analyses (right dPMC in green and SPL in blue) displaying commonalities (in turquoise) and differences in the connectivity patterns. All effects were significant at cluster-level *p* < 0.05 (cFWE-corrected) with a cluster-forming threshold of *p* < 0.001 at voxel level. **Abbreviations.** dPMC: Dorsal premotor cortex, gPPI: Generalized psychophysiological interaction, SPL: Superior parietal lobule, RRC: Response–response congruent, RRI: Response–response incongruent.

When assessing age differences in task-related connectivity for the right dPMC seed in dual R-R congruent (i.e., [Dual_RRC_: YA > OA] ∩ [Dual_RRC_: YA +]) and R-R incongruent (i.e., [Dual_RRI_: YA > OA] ∩ [Dual_RRI_: YA +]) PPI regressors separately, we did not observe any significant results. In contrast, only for young (vs. older) adults, right SPL presented increased connectivity to bilateral supramarginal gyrus and IPS, as well as right frontal operculum and anterior insula during dual R-R congruent trials (see **Figure 8A**). Furthermore, during dual R-R incongruent trials, young (vs. older) adults showed increased task-related connectivity between right SPL and right supramarginal gyrus (see **Figure 8B**). No hyperconnectivity patterns were observed for older adults for either seed. For more details on age-related differences obtained from the gPPI analysis, please refer to **Supplementary Table S13**.

**Figure 8.**
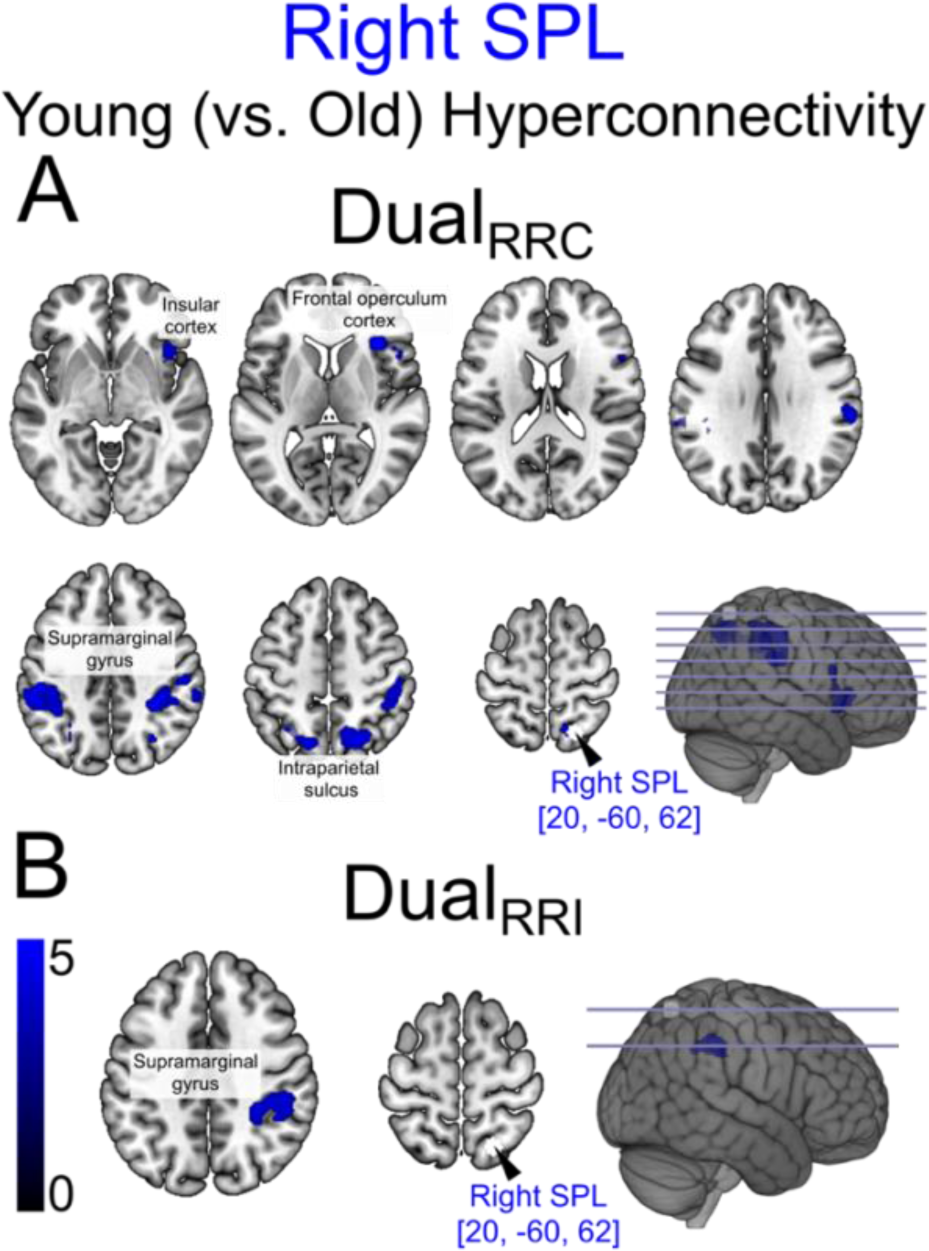
Results of the generalized PPI analysis, testing for age-related differences in the connectivity of right SPL during **(A)** Dual_RRC_ (vs. baseline) and **(B)** Dual_RRI_ (vs. baseline) PPI regressors. Each contrast was assessed in separate models with the age-specific conjunctions **(A)** [Dual_RRC_: YA > OA] ∩ [Dual_RRC_: YA +]) and **(B)** [Dual_RRI_: YA > OA] ∩ [Dual_RRI_: YA +]), respectively, and masked with dual-task conflict-related network (Dual_RRI_ + > Dual_RRC_; see **Figure 3C**). All effects were significant at cluster-level *p* < 0.05 (cFWE-corrected) with a cluster-forming threshold of *p* < 0.001 at voxel level. **Abbreviations.** OA: Older adults, SPL: Superior parietal lobule, RRC: Response–response congruent, RRI: Response–response incongruent, YA: Younger adults.

## Discussion

The present study aimed to elucidate brain activity patterns associated with processing response-code conflict in dual-tasking and their modulation by age. Young and older adults underwent fMRI while performing an auditory-manual single-stimulus onset paradigm with one versus two concurrent speeded reaction tasks (Huestegge and Koch 2009; Weller et al. 2022; Paas Oliveros et al. 2023). To manipulate response-code conflict in dual conditions, we had each hand’s response codes (i.e., response location called for by the tone’s pitch) mutually congruent or incongruent.

First, response-code conflict was associated with increased brain activity in several MDN regions (Duncan 2010; Camilleri et al. 2018). Moreover, when facing dual-task conflict, older adults engaged left superior frontal gyrus more strongly than did young adults. Second, we investigated how conflict-related brain activity and age differences therein relate to individual dual-task performance and various cognitive abilities during R-R incongruency. More efficient dual-task performance went along with lower visual cortex recruitment; and decreased subcortical, parietal, and cerebellar activity was modulated by lower crossmodal attentional or task-switching performance. Additionally, older adults showed increased engagement of left IPS and right cerebellum associated with lower attentional performance. Third, we explored whether conflict-sensitive brain regions (i.e., right dPMC and SPL) change their connectivity to other task-relevant brain areas depending on the level of dual-task difficulty and whether this is affected by age. Conflict-modulated connectivity was not sensitive to dual-task response-code conflict for either seed region or age-related effects. In contrast, when assessing connectivity commonalities between dual R-R congruency and incongruency, both seeds showed strong coupling with bilateral fronto-parieto-insular areas but with limited overlap. Only young (vs. older) adults showed increased synchronization between right SPL and parietal areas, right frontal operculum, and insular cortex during dual R-R congruency, and between right SPL and right supramarginal gyrus during dual R-R incongruency. In the following sections, we discuss our results in detail.

### Behavioral data

Our behavioral results are a within-scanner replication of our previous findings (Paas Oliveros et al. 2023). Consistent with our previous study, our dual-task paradigm elicited response-related conflict by having spatially opposing S-R mapping rules for either hand.

The detrimental effect of response-code conflict between concurrent responses was evident in all behavioral measures (dual-task costs on BIS, RT, and ER), with higher dual-task costs on R-R incongruent (vs. congruent) trials. This effect aligns with previous studies showing that dual-response conditions with a single S-R mapping rule involve a single conjoint response selection process for both responses. This efficient selection approach is reflected in relatively low dual-response performance costs overall and an equivalence in costs for the (easy) S-R compatible and the (more demanding) incompatible condition, as compared to conditions with two independent and potentially conflicting mapping rules like in our R-R incongruent condition (Fagot and Pashler 1992; Pieczykolan and Huestegge 2018; Weller et al. 2022; Paas Oliveros et al. 2023).

It is assumed that dual-response conditions with equivalent response codes, which enable conjoint response selection, restrict the costs to downstream execution-related sources, such as disinhibiting contralateral parts of the motor system or synchronizing movement kinematics. In contrast, the asymmetric cost allocation in R-R incongruent trials, in which S-R compatible responses showed higher dual-task costs than S-R incompatible ones, suggests that implementing two conflicting response codes concurrently necessitates two separate task representations in which the more demanding (S-R incompatible) task is likely to be prioritized over the easier (S-R compatible) one. We also found that participants did not only implement a response grouping strategy for shielding tasks against crosstalk, by which the first or less demanding task is processed first, while its execution gets delayed until the second or more demanding task has been processed too, so that the execution of both responses can be synchronized (Pashler 1994; Miller and Ulrich 2008). Rather, the still asymmetric costs in trials with non-grouped dual responses are consistent with the notion of a strategic prioritization of limited processing capacity, according to which the more demanding S-R incompatible response mapping would have received more processing resources than the less demanding one (Huestegge and Koch 2013; Pieczykolan and Huestegge 2014; Paas Oliveros et al. 2023). An alternative explanation to this prioritization strategy holds that instead of resulting from an active strategic decision based on perceived difficulty, the effectiveness to shield (sub)tasks is a more natural side-effect of an already existing bias in attentional resource allocation to the more difficult task.

Replicating our earlier findings (Paas Oliveros et al. 2023), we observed that response-related interference was accentuated with advanced age, as shown by the significant interaction effect between R-R congruency and age on all performance measures (dual-task costs on BIS, RT, and ER). New to this study, we employed the BIS, a recently introduced behavioral measure that jointly reflects performance speed and accuracy (Liesefeld et al. 2015; Liesefeld and Janczyk 2019), to account for potentially different speed–accuracy trade-off criteria. A three-way interaction effect on dual-task BIS costs reflected a more pronounced cost asymmetry among older (vs. young) adults in R-R incongruent (vs. congruent) conditions. Thus, older adults appear to be particularly susceptible to task confusability when two mutually incongruent response codes need to be processed concurrently, as compared to conditions with two identical response codes. From the cognitive perspective, these age-related deficits can be partially explained by generalized slowing(Salthouse 1996) or relying on response grouping (i.e., internal queuing of the central processing stages of both tasks), as the interaction between R-R congruency and age disappeared on dual-task speed costs after controlling for these two processes. Alternatively, older adults might suffer from inhibitory deficits affecting the attentional mechanisms for task processing and scheduling attention across different task sets, causing distraction between parallel processing streams (Mayr 2001; Mayr and Liebscher 2001; Hein and Schubert 2004; Paas Oliveros et al. 2023). Furthermore, older adults may voluntarily allocate more attentional resources to the more demanding task, leaving the less demanding one largely unattended, possibly explained by an over-reliance on central attention with advanced age (Maquestiaux and Ruthruff 2021). The overexerted top-down attention to one task set would then harm overall performance.

We would like to mention that the response buttons used in the MRI scanner registered the button presses in 8-ms intervals, eliciting a small binning effect on the response times. Although this constituted a negligible effect of undirected noise, it would be advisable for future studies in this domain to use a response recording setup that is able to register millisecond-level differences between responses. Finally, our analysis showed that the BIS, jointly capturing performance speed and accuracy, offers greater sensitivity than the separate analysis of dual-task speed and accuracy costs, as evidenced by the significant three-way interaction effect on the BIS only. In the same context, the BIS appears to be robust to differences in the speed–accuracy trade-off, which could otherwise reduce age differences when looking only at speed costs.

### Neuroimaging data

#### Aim 1: Brain activity correlates of response-code conflict in dual-tasking and their age-related differences

In a first step, we assessed the **neural correlates of dual-response execution** by contrasting R-R congruent vs. single-task trials (see **Figure 3A**). This revealed an activation pattern that fits well with the action-focused nature of our experimental paradigm, involving predominantly bilateral primary somato-motor (including M1, dPMC and SMA, basal ganglia, and cerebellum) and parietal areas. Compared to single unimanual conditions, the dual R-R congruent condition did not involve major parts of the MDN crucial in task-set maintenance and conflict resolution (Botvinick et al. 2004; Cieslik et al. 2015; Worringer et al. 2019). Instead, increased activity was mainly found in motor-related regions. This agrees well with our conclusion from the behavioral findings about a conjoint response selection process when congruent responses have to be executed, which would not evoke the need for controlling two different (and even conflicting) task sets in parallel via recruiting the MDN at large. Our results also corroborate earlier observations that additional motor output during bimanual response execution is mainly subserved by motor regions (Jäncke et al. 2000; Nair et al. 2003; Swinnen and Gooijers 2015). Thus, the activation of primary motor areas in our dual-response condition lends support to previous findings suggesting an upregulation of these areas when both hands are required, in contrast to single-hand reaction tasks. In this context, posterior parietal areas are thought to integrate somatosensory information, perform spatial transformations, such as spatial information implied by the auditory stimulus, and project to frontal and prefrontal regions to adjust motor responses accordingly (Rizzolatti et al. 1998; Jäncke et al. 2000; Grefkes et al. 2004; van Dun et al. 2021). Within the framework of motor control models, the cerebellar activation we observed agrees well with its implication in monitoring cortical output and giving feedback on correct motor execution and coordination (Nair et al. 2003; van Dun et al. 2021). To summarize, our results are in concordance with previous literature suggesting that the upregulation in motor, parietal, and cerebellar areas arises from higher demands for motor output and spatial transformations in dual-versus single-hand reaction tasks.

In addition, reduced activity was found in bilateral occipital and medial prefrontal areas, which are again consistent with the nature of our auditory-manual dual-task paradigm, during which participants did not engage in any visual processing besides fixating a small cross presented centrally on the screen. This focus on auditory stimuli should, in turn, lead to disengaging attention from the visual modality, which is known to go along with reduced activity in visual regions (Mozolic et al. 2008; Langner et al. 2011). This modality-specific focus and downregulation appear to have been stronger in the slightly more demanding dual-response conditions, relative to the single-response ones. Likewise, the downregulation of a region in medio-frontal cortex, an area of the default-mode network (DMN), is consistent with the nature of our externally focused task (Raichle et al. 1996; Fox et al. 2005). In line with the notion of efficient resource allocation, the not needed and to-be-deactivated DMN was more strongly suppressed during the somewhat more demanding dual-response condition.

In a second step, we tested for **brain regions related to dual-tasking at large** by contrasting R-R incongruent with single-task trials (see **Figure 3B**). Beyond primary motor areas previously observed in the dual-response network, we identified a large fronto-parieto-insular network covering regions of the extended MDN, such as dPMC, preSMA, MCC, dlPFC, anterior insula, IPS, thalamus and cerebellum (Duncan 2010; Müller et al. 2015; Camilleri et al. 2018). These regions have been consistently associated with top-down executive control, as well as dual-and multi-tasking (Stelzel et al. 2006; Szameitat et al. 2006; Stelzel et al. 2008; Stelzel et al. 2009; Deprez et al. 2013; Al-Hashimi et al. 2015; Papegaaij et al. 2017; Worringer et al. 2019). Furthermore, dual-tasking also induced suppression of right occipital and ventromedial frontal activity, consistent with the disengagement of visual processing and DMN suppression, respectively, as already discussed in the context of dual-response execution.

Third and most importantly, we contrasted R-R incongruent with congruent dual-task conditions to assess **brain activity specifically related to dual-task response-code conflict** (see **Figure 3C**), which already showed to have a distinct behavioral effect (see **Figure 2**; Paas Oliveros et al., 2022). This contrast revealed increased activity in a similar fronto-parieto-insular network as previously observed for dual-tasking at large, but no significant reductions in activity. Of note, the peak activations, indicating maximum sensitivity to response-related crosstalk, were located in right dPMC and SPL (see **Supplementary Table S5** for details), which were chosen as seed regions in the analysis of task-modulated connectivity. Right SPL has been previously associated with top-down attentional control and shifting (Corbetta and Shulman 2002; Al-Hashimi et al. 2015), but also with auditory spatial working memory (Alain et al. 2001; Alain et al. 2008) and, in particular, the spatial representation of, and transformation between various coordinate systems, such as translating visuo-spatial information into hand-centered coordinates for memory-guided finger movements (Grefkes et al. 2004; Langner et al. 2014). These processes become relevant in our task when attending to the pitch of auditory stimuli and mapping this information to the corresponding finger response, even more so when concurrently processing two opposing S-R mappings. The dPMC, in turn, is involved in cognitive action control and appears to be essential for action formulation, preparation, and execution, especially under conditions of parallel interference from concurrent movements (Abe and Hanakawa 2009; Hardwick et al. 2015; Genon et al. 2016; Worringer et al. 2019). In our study, its activity most likely reflects the preparation of the corresponding correct motor response in the demanding settings of response-related dual-task conflict.

Compared to the global dual-task network (cf. **Figure 3B**), the specifically conflict-related network is more constrained. For instance, the primary motor and somatosensory cortices were not included, which is congruent with the notion that these areas are involved in controlling bimanual action. Nonetheless, this more circumscribed response-conflict network includes key areas of the extended MDN, a domain-general network involved in executive control, including working memory, attention and action planning and inhibition (Duncan 2010; Müller et al. 2015; Camilleri et al. 2018). In part, this was expected because managing dual-task response-code conflict involves complex cognitive action control to resolve between-set interference and guide subprocesses of attention shifting and S-R mapping, depending on the location implied by the pitch (Worringer et al. 2019). Thus, the present study complements previous findings regarding our first aim and hypothesis by demonstrating that the MDN is also involved in response-related dual-task crosstalk. Still, it cannot be separated from other processes that attempt to prevent or resolve spatial dual-task interference at the effector level. As mentioned earlier, dual-tasking is a higher-order cognitive process that implicates several cognitive functions, including activating and maintaining two task sets, attentional control, and coordinating multiple actions. Given the involvement of other cognitive abilities during dual-tasking, which may vary among individuals, the covariance analysis aimed to characterize better the inter-individual differences in the activity of brain areas of the conflict-related network, as discussed later.

When assessing the impact of aging, we observed no significant **age differences in brain activity related to dual-response execution** (i.e., R-R congruent vs. single-task contrast). This agrees with the absence of age differences in dual-execution costs during R-R congruency (see **Figure 2**) but disagrees with a previous study showing that older adults more strongly recruited medial motor, somatosensory, and prefrontal areas as well as IPS when performing bimanual movements (Goble et al. 2010). However, these results were observed with hand movements that had to be performed in mutually congruent and incongruent directions, while no spatially imperative stimuli had to be processed (Goble et al. 2010). Moreover, the dual-hand motion conditions were contrasted to rest, whereas we here contrasted them to single-task trials. Thus, the neural correlates of dual-response execution have not been isolated specifically in Goble and colleagues’ (2010) study.

In contrast to dual-response execution, **dual-tasking at large** (R-R incongruent vs. single-task trials) induced stronger activation in older adults in several medial motor (SMA), cingulate and prefrontal areas (see lower section in **Figure 4A**). Previous studies manipulating the complexity of partially similar coordination tasks with dual-versus single-hand movements, such as antiphase movements, have traditionally found activations in the SMA (Debaere et al. 2004; Swinnen and Gooijers 2015). Among the elderly, SMA activity appears to have a critical compensatory role during tasks with increased coordination demands (Goble et al. 2010). Furthermore, ample research on inhibitory and attentional control demonstrated that anterior cingulate and lateral prefrontal areas become active when solving response conflict to suppress inadequate response tendencies (Corbetta and Shulman 2002; Botvinick et al. 2004; Vossel et al. 2014; Cieslik et al. 2015). More specifically, the brain regions observed here have also been shown to be upregulated among older adults when coordinating conflicting limb movements (Heuninckx et al. 2005). Thereupon, we conclude that the hyperactivation in SMA, cingulate and prefrontal areas among older (vs. young) adults may arise from older adults’ encountering increased difficulties in attending and processing incongruent dual-task sets that subsequently require the coordination and execution of spatially conflicting responses, but also inhibiting inadequate responses.

Noteworthy, our previous age-related results contrast with other dual-task studies. For instance, van Impe and colleagues (2011) found that during a drawing and addition task, older adults showed increased activity in different brain regions, including right precentral gyrus, ventrolateral prefrontal cortex, bilateral superior parietal gyrus, precuneus, and left cerebellum. However, this hyperactivation was mainly driven by the drawing task and explained by differences in goal-directed attentional processes and an age-related increase in sensory feedback reliance to guide hand movements. Another study by Papegaaij and colleagues (2017) reported hyperactivation in older adults’ motor and occipital areas, left MFG, supramarginal gyrus, and precuneus during a balance-calculation task. However, age differences analysis only considered dual-task but no single-task regressors. Consequently, the differences in results could be partially due to discrepancies in experimental paradigms and statistical contrasts, which isolated somewhat different cognitive subprocesses within dual-tasking.

In turn, young (vs. older) adults showed lateralized hyperactivation in smaller clusters in right primary motor cortex, superior frontal gyrus, and SPL (see upper section in **Figure 4A**). As most participants in this study were right-handed, this hyperactivation may reflect the increased demand for attentional and executive control processes that modulate activity associated with controlling responses using the non-dominant left hand, necessary for efficient dual-task performance. Prior research has demonstrated that when a task requires a more demanding movement by the non-dominant hand, the contralateral motor cortex becomes more active, indicating greater engagement with increasing motor task difficulty, particularly in novel tasks (Jäncke et al. 2000; Haslinger et al. 2004). Conversely, it has been suggested that in older adults, the superiority of the dominant hand diminishes (Kalisch et al. 2006), and the brain’s functional lateralization pattern attenuates (Cabeza 2002; Loibl et al. 2011; Hill et al. 2020). Therefore, the reduced motor and hand asymmetry in older adults may explain why the motor area of the non-dominant hand becomes more strongly activated in young adults. In contrast to the age differences during dual-tasking, young (vs. older) adults did not appear to recruit additional neural resources specifically related to resolving dual-task response-code conflict.

Complementing the age-related differences in brain activity observed during dual-tasking at large, we found hyperactivation in the left superior frontal gyrus among **older adults when coping with dual-task response-code conflict** (i.e., R-R incongruent vs. congruent contrast; see **Figure 4B**). Within the literature of neurocognitive aging models, this finding could be expected and is in concordance with an increased vulnerability to age-related changes in middle and superior frontal areas (Greenwood 2000), as well as with an over-recruitment of prefrontal resources with advanced age as a compensatory mechanism for cognitive action control or executive functioning (Reuter-Lorenz and Cappell 2008; Park and Reuter-Lorenz 2009; Seidler et al. 2010; Cabeza et al. 2018; Li et al. 2018; Spreng and Turner 2019). However, the specific upregulation of the left superior frontal gyrus among older adults coincides only partially with previous studies. For example, Chmielewski and colleagues (2014) identified significant age differences in MFG and SFG according to the temporal overlap of two concurrent tasks, but in this case, older participants showed functional down-regulation in these areas. In contrast, other studies have shown a stronger activation in inferior and superior frontal regions among older adults during task interference resolution (Langenecker et al. 2004; Zhu et al. 2010). Together with our behavioral observation of an increased task confusability and interference susceptibility among the elderly, at first glance, the hyperactivation of the left SFG could represent a compensatory mechanism to counteract age-related brain structural and functional decline. This would support our first hypothesis on possible compensatory mechanisms during response-code conflict in aging. However, this topic will be discussed in more detail in the following section when incorporating the findings assessing associations between conflict-related brain activity and individual levels of dual-task performance.

Overall, it appears that dual-response execution mainly requires input from motor and parietal areas and is well preserved in advanced age. It is rather in dual-tasking proper, when more demands on top-down cognitive control are put, that age differences arise, with a specific cluster in the left SFG particularly recruited when dealing with response-code conflict in dual-tasking. Thus, the observed brain activity and the age-related differences along the increased cognitive loading in our dual-task paradigm are indicative of an age-related shift along the continuum from less demanding motor control processes towards more demanding cognitive action control and processing two simultaneous tasks with mutually incongruent spatial response codes.

#### Aim 2: How brain activity is linked to individual levels of dual-task performance and related cognitive abilities

Next, we investigated how conflict-related brain activity, and age differences therein, were associated with individual dual-task performance as well as three other related cognitive abilities: (1) performance in the S-R compatible, R-R incongruent dual-task condition (i.e., the response featuring the highest dual-task costs on average), (2) crossmodal selective attention, (3) spatial working memory span, and (4) global task-switching costs. Although dual-task performance and the covariates showed phenotypical correlation, the neural effects were found to be distinct, which will be discussed below.

First, we observed that **dual-task performance** did not show positive correlations with brain activity in task-related regions but only a negative one with activity in visual cortex, highlighting the relevance of down-regulating task-irrelevant basic visual processing for keeping dual-task costs low in the most challenging condition (Hairston et al. 2008; Mozolic et al. 2008; Langner et al. 2011). Although we would have expected shared variance between dual-task performance and task-relevant areas, such as regions commonly associated with executive control, including the middle and superior frontal gyrus, medial frontal cortices, and inferior and superior parietal lobules (Saylik et al. 2022), the downregulation of visual areas is consistent with our non-visual paradigm, which requires focusing on auditory input processing and preventing distraction by task-irrelevant visual input. A previous study emphasized the significance of deactivating visual cortical areas to filter non-relevant information, particularly in challenging auditory tasks (Hairston et al. 2008). This intrinsic filtering mechanism can be an asset in suppressing visual distractions and enhancing task performance. However, the association between dual-task performance variability among individuals and brain regions beyond the dual-task network could imply that the group contrast analysis may overlook essential brain information to predict dual-task performance accurately, but this hypothesis needs further exploration. Our findings suggest that the negative correlation between medial occipital cortex activity and dual-task performance demonstrates a critical connection between visual information suppression and improved performance in auditory dual-task settings.

Next, we looked for associations between brain activity in the task-specific network and cognitive abilities related to dual-tasking. Reduced thalamic and globus pallidus activity was associated with lower **crossmodal attentional performance** (see **Figure 5B**), while lower activity in posterior parietal regions (SPL/IPS) and right cerebellum went along with lower task-switching performance (i.e., higher global task-switching costs) (see **Figure 5C**). Given that both thalamus and globus pallidus have been frequently linked to sensorimotor functions and movement regulation (Sommer 2003), as well as to the extended MDN (Camilleri et al. 2018), finding subcortical activation associated with crossmodal attention could be interpreted as reflecting an attentional requirement for executing two tasks with response-code conflict. More specifically, this agrees with the behavioral observation of a strategic prioritization of limited processing capacity or a bias in attentional resource allocation under R-R incongruency. Participants appeared to allocate more attentional resources to the more demanding S-R incompatible response mapping than to the less demanding one, which could have potentially involved the thalamus and globus pallidus.

Regarding the associations with **task-switching performance**, it is noteworthy that SPL and IPS are integral parts of the dorsal attention network, which is crucial in the top-down control of spatial attention (Corbetta and Shulman 2002; Corbetta et al. 2008). Here, we demonstrated that bilateral IPS and SPL activity exhibited shared variance with task-switching performance within the conflict-sensitive network, highlighting their relevance in this cognitive process. Consequently, these regions appear to be particularly important for the flexible allocation of attentional resources to two concurrent task sets that conflict spatially. This association agrees well with previous multi-tasking studies covering other cognitive abilities. Worringer and colleagues (2019) reported meta-analytic across-study convergence of dual-tasking and task-switching activity in the IPS but also in the dPMC and anterior insula. Closely related to this finding, previous studies have highlighted the role of the IPS during response selection and S-R mapping processes, as well as the reorientation and maintenance of spatial motor attention (Rushworth et al. 2001; Iacoboni 2006; Cieslik et al. 2010; Cieslik et al. 2015; Camilleri et al. 2018; Worringer et al. 2019; Saylik et al. 2022). Lower task-switching performance was additionally associated with decreased conflict-sensitive activity in the right cerebellum. While the cerebellum has been traditionally associated with motor functions, previous studies involving dual-tasking (Collette et al. 2005; Deprez et al. 2013; Wu et al. 2013) and task-switching settings (Dreher 2003) have also identified activity in cerebellum, reflecting its relevance in higher-order cognitive processing. This suggests that cerebellar areas play an additional role in maintaining and coordinating two task sets concurrently rather than solely reflecting their motor functions (Deprez et al. 2013). Nonetheless, it should be mentioned that the studies included in the meta-analysis (Worringer et al. 2019) mainly assessed task-switching using local switching costs and dual-task studies manipulating input-related and response-selection interference. In contrast, we analyzed global task-switching costs, and our dual-task paradigm manipulates response-code interference. Global task-switching costs stem from increased demands of maintaining two task configurations in mixed blocks compared to pure blocks. These costs arise when two task sets are mixed, inducing difficulties in maintaining each task set over time, tracking and adjusting each set’s activation level, and keeping them apart from each other. In our dual-task setting, it is conceivable that activating the associated brain regions more strongly is necessary when involving motor attention for processing two simultaneous and conflicting task sets and generating corresponding action plans. We infer that participants who experience difficulties in these processes and have lower activity in posterior parietal areas and cerebellum exhibit higher costs on performance when being faced with mixed task sets. Overall, increased activity in posterior parietal cortex and cerebellar areas, which are recruited during response-code conflict in dual-tasking, appear to be specifically associated with inter-individual differences in effectively and flexibly allocating and maintaining spatial motor attentional resources (Rushworth et al. 2001; Iacoboni 2006; Weiss et al. 2006) across two concurrent task sets with different spatial S-R mappings, and coordinating them to develop motor plans accordingly.

Another dual-task-related construct we analyzed was **working memory**. When maintaining and processing S-R mappings of two simultaneous task sets (i.e., dual-tasking), more working memory capacity should be required compared to single-task conditions. In addition, a recent study found a robust positive association between activity in multiple-demand regions and individual differences in working memory performance (Assem et al. 2020). Thus, we would have expected our task-related regions, which largely overlap with the MDN, to reveal shared variance with working memory performance. However, we did not find any significant neurofunctional correlations. The lack of any associations with working memory performance could be due to the fact that working memory abilities are material- and process-specific. For instance, the working memory task used in this study, the Corsi block-tapping task, was designed to evaluate the participants’ ability to recall the sequence in which irregularly arranged cubes were tapped, emphasizing the retention of visuo-spatial inputs and their respective order. In contrast, our dual-task paradigm did not require retention of any visuo-spatial information, and the concurrent task sets only varied in terms of the spatial S-R mappings towards the pitch. As such, the maintenance and transformation of visual input coordinates into motor coordinates required in the Corsi block-tapping task might have too little processing overlap with the concrete requirements of coping with crosstalk at the response level. Hence, it would be advisable that future studies interested in the shared variance between cognitive processes put special care on each task’s specific characteristics and requirements. In our case, an (auditory-)motor or more complex working memory task involving memoranda from two different tasks (Engle et al. 1999; Kane et al. 2004; Pläschke et al. 2020) could be considered.

The **age differences found in the cognitive modulation of conflict-related brain activity** during R-R incongruency were informative to complement the task-activation results. Although we did not find any age differences associated with dual-task performance, working memory, or task-switching, we did find them with crossmodal attention. Here, the activity within left IPS and right occipito-cerebellar areas during R-R incongruency was differently modulated by crossmodal attention abilities in both age groups. More specifically, the increased activity within these areas during R-R incongruency was significantly linked with detrimental crossmodal attentional performance (i.e., longer RT) among older adults. While the observed age differences in task-related activation pointed to a possible compensatory over-recruitment of left SFG (see the previous section of the discussion, **Aim 1**), this should be accompanied by a positive correlation between activation and dual-task behavioral performance, which we did not find. On the other hand, the dedifferentiation theory predicts more widespread activation patterns with a loss in regional specificity and without any behavioral improvement (i.e., no correlation with performance or an association between decreased/increased regional activity but with detrimental cognitive performance). Thus, the age differences in the negative correlation of left IPS and right occipito-cerebellar areas with attentional performance point towards a dedifferentiation pattern or inter-individual variability in attentional strategies among older adults (Park et al. 2001; Voss et al. 2008) and refute the hypothesis of a compensatory mechanism. As mentioned before, the IPS is a key area of the dorsal attention network (Corbetta and Shulman 2002; Corbetta et al. 2008) and is relevant for spatial motor attention (Rushworth et al. 2001; Iacoboni 2006; Weiss et al. 2006). Therefore, the dual-task age differences in this brain area’s association with attentional performance are consistent with our behavioral results of a process-specific increase in task confusability among older adults. Nonetheless, since the behavioral effects are also partially explained by an age-related generalized slowing, an additional mechanism contributing to these findings could be that neural dedifferentiation in left IPS among older adults is accompanied by a tendency to be generally slower in flexibly allocating spatial motor attentional resources between conflicting S-R mappings.

#### Aim 3: Changes in task-modulated connectivity and their age differences

Our final aim was to explore how connectivity between task-sensitive brain regions and the age differences therein differ under dual-task response-code conflict, as well as R-R congruency and incongruency. For this purpose, we investigated psychophysiological interactions, for which we defined two seed regions in right dPMC and SPL based on the peak activations during dual-task response-code conflict. These are key regions for multi-tasking (Worringer et al. 2019), but they have also been consistently associated with a broad range of motor and cognitive functions.

As mentioned above, SPL, as part of the dorsal attention network, has been associated with flexible top-down control of spatial motor attention (Rushworth et al. 2001; Iacoboni 2006; Weiss et al. 2006), as well as with coordinate transformations between visuo-spatial stimuli and hand-centered movements (Grefkes et al. 2004; Langner et al. 2014), key processes for correct performance in our dual-task paradigm. Likewise, dPMC is essential in sensorimotor integration, movement preparation, response selection, motor learning, and working memory (Abe and Hanakawa 2009; Hardwick et al. 2015; Genon et al. 2016). Our seed overlaps with the central dPMC as defined by a connectivity-based parcellation study, the core region of the dPMC, coupled to all other dPMC clusters and with solid connections to IPS and SPL (Genon et al. 2016). With respect to dual-tasking, it was assigned a crucial role in intentionally formulating and controlling the execution of actions under conditions of interference from concurrent movements (Worringer et al. 2019).

Regarding the third aim of this study, we observed that **task-modulated connectivity did not differ during dual response-code conflict** (R-R incongruent vs. congruent regressors) for both seed regions, and there were no significant age differences. Thus, while we found brain activation to be affected by response-code conflict elicited in our dual-task paradigm, the connectivity between highly conflict-sensitive seeds and the remaining conflict-related network appears to be neither sensitive to the level of conflict nor to age. Examining **task-global connectivity changes across both R-R congruent and incongruent conditions** (vs. implicit baseline, respectively), we observed commonalities and differences between the two seed regions (see **Figure 7**). Both seeds’ connectivity showed modest overlap in bilateral parietal (SPL, IPS, precuneus, and supramarginal gyrus) and frontal (dPMC, SMA, ACC, and SFG) cortex, as well as the right anterior insular cortex and frontal operculum. Although both seed regions are part of the dorsal attention network and strongly connected to each other, interestingly, their individual connectivity patterns were located just next to each other with relatively little overlap. Specifically, right SPL displayed increased connectivity with extensive regions in bilateral parietal and ipsilateral frontal cortex, along right frontal operculum, subcortical areas, and the cerebellum. In contrast, right dPMC demonstrated increased connectivity, particularly with left lateral PFC, insula, and parietal regions, as well as a frontomedial cluster.

The connectivity pattern from right SPL is reminiscent of areas within the dorsal attention network, which are thought to facilitate attentional shifting between sensory stimuli and their locations, enabling individuals to respond accordingly (Corbetta and Shulman 2002; Corbetta et al. 2008). Moreover, it is associated with regions that support top-down action control mechanisms, which mediate conflict resolution arising from spatial S-R incompatibility (Cieslik et al. 2010; Cieslik et al. 2015). Interestingly, we also found pronounced coupling with regions associated with motor functions, including parts of the subcortex such as thalamus and putamen (Caspers and Zilles 2018) as well as cerebellar regions, likely reflecting the need for basic motor coordination processes. The connectivity pattern observed for the premotor seed converges with the functional coupling described for the central dPMC cluster in a previous connectivity-based parcellation study (Genon et al. 2016). In our case, the connectivity additionally covered medial and insular regions, which are rather linked to the dorsal part of the dPMC. This premotor cluster is engaged in hand and finger movements, such as finger-tapping paradigms (Genon et al. 2016). Our results on connectivity patterns resemble regions also found in the brain activation analyses, which form part of the dorsal attention network and are known to be engaged with increasing demands for top-down control, such as allocating spatial motor attention and resolving spatial interference. However, as these results were derived from a more lenient contrast (for each seed, the minimal conjunction of each R-R congruency level contrasted with the implicit baseline) instead of the response-code conflict contrast, one could argue that the synchrony between conflict-sensitive brain areas is, in general, relevant for processing simultaneous S-R mappings, but it is rather the level of activity that becomes crucial when solving response-code conflict in dual-tasking.

Finally, the **age comparisons for task-modulated connectivity** did not reveal any hypo- or hyperconnectivity patterns for older adults with any of the seeds. In contrast, only young (vs. older) adults showed increased connectivity between right SPL and bilateral supramarginal gyrus, contralateral SPL and IPS, right frontal operculum, and insular cortex during dual R-R congruent (vs. baseline) trials (see **Figure 8A**). The connectivity between right SPL and right supramarginal gyrus in young adults appeared to be crucial during dual-tasking because this was the only pair of brain areas that were highly connected during dual R-R incongruent (vs. baseline) trials when compared to older adults (see **Figure 8B**). The connectivity agrees with the well-supported anatomical link between superior parietal cortex and supramarginal gyrus, arguably coordinating attentional networks and a top-down attentional reorientation to behaviorally relevant stimuli (Corbetta and Shulman 2002; Catani et al. 2017). Furthermore, previous research has reported age differences in connectivity within regions supporting cognitive control, with young adults exhibiting increased connectivity within attentional networks when compared to older adults (Madden et al. 2010; O’Connell and Basak 2018). The task-modulated connectivity pattern between right SPL and supramarginal gyrus in young (vs. older) adults may, thus, be linked to more efficient top-down attentional maintenance and monitoring mechanisms when task complexity increases during dual-tasking but does not appear specific for dual-task response-code conflict, as we did not see any difference when contrasting the R-R congruency conditions. Likewise, conflict-related activity appears to be sensitive to age, but conflict-related connectivity does not. Our results contrast with previous studies showing age differences in task-modulated connectivity associated with cognitive control and executive functioning (Lamichhane et al. 2018; O’Connell and Basak 2018). These differences, however, might be due to the differences in paradigms and the seeds derived from peak activations during task-specific contrasts. Nonetheless, this discrepancy remains to be further investigated in future work.

## Conclusions

Implementing a novel dual-task paradigm in the MRI scanner, we uncovered the neural correlates of response-code conflict arising between two concurrent actions. The replication of asymmetric dual-task costs occurring with response-code conflict corroborates the assumption of a flexible allocation of attentional resources and a strategic prioritization of limited processing capacity. Finding the deleterious effects of incongruent response codes exacerbated in advanced age supports the notion that, besides generalized slowing, aging may be linked to heightened response-code confusion due to deficits in shielding different task sets from each other. At the neural level, we found that conflict-free dual-response execution does not require support from large parts of the MDN but mainly relies on motor and parietal areas. In contrast, when response-related crosstalk is induced, an extensive fronto-parieto-insular network is recruited, covering key regions of the extended MDN, known to be involved in constructing mental control programs, controlling spatial motor attention, resolving spatial interference, and planning and executing motor responses accordingly. While conflict-free dual-responding did not evoke activity differences between the two age groups, older adult’s exhibited stronger engagement of bilateral motor and prefrontal regions during dual-tasking involving response-code conflict. This finding suggests a non-compensatory shift in neural recruitment along the continuum from less demanding motor control processes to more complex cognitive action control processes, indicating that older adults recruit more neural resources when faced with the same conflicting dual-task as younger adults.

However, the individual efficiency in resolving response-code conflict appears to mainly depend on effectively suppressing task-irrelevant visual cortex activity. Given the broad group-level recruitment of the MDN during conflict-laden dual-tasking, the lack of substantial associations between conflict-related activity levels and dual-task performance as well as other, somewhat related higher-order cognitive abilities can hardly be taken as evidence for the specificity of the cognitive processes taxed by our paradigm. It rather suggests that the recruitment of the MDN may be an all-or-none phenomenon, whose individual extent or intensity would not be predictive of individual performance efficiency. Still, the observed covariance between conflict-related activity in motor-parietal areas and both the ability to distribute attention across two sensory modalities and to cope with two frequently alternating task sets shows the fundamental sensitivity of our approach to detect selective associations with related constructs. It also suggests some functional specificity among the large set of regions coactivated when grappling with between-task conflict. Finally, our connectivity analyses revealed that the synchronization between our premotor or parietal seed regions and the remaining conflict-related network is neither conflict-specific nor sensitive to age. This context-insensitivity demonstrates a remarkable robustness of the coupling among MDN regions, possibly to ensure the integrity and functioning of this network across diverse task states that all require the top-down modulation of cognitive processing of some kind.

Overall, our study revealed that resolving response-code conflict in dual-tasking involves substantial parts of the domain-general MDN, a network pivotal for biasing action decisions in the service of goal-oriented coherent behavior. The breadth and stability of the MDN’s involvement, however, amplifies the question for the neural basis of intra- and inter-individual differences in the efficiency of its top-down modulatory activity and calls for future research to address this seeming discrepancy, not just in the domain of dual-tasking.

## Notes

### Competing Interest Statement

The authors have declared no competing interest.

