## SupplementaryMaterial for "Brain functional characterization of response-code conflict in dual-tasking and its modulation by age"

### Supplementary Material

#### Psychometric assessment

1. *Crossmodal attention:* Crossmodal attention was assessed via the selective and focused attention subtests (WAF-S and WAF-F, respectively) from the Perception and attention functions battery from the Schuhfried Vienna Test System (https://marketplace.schuhfried.com/en/waf; visuo-auditory crossmodal test form S1). In the *selective attention task*, participants were presented with circles, squares, high and low tones. The figures could become brighter, darker, or remain the same; the tones could become louder, softer, or remain the same. Participants were asked to respond only when a circle got brighter, or a low tone got louder; changes in squares or high-pitched tones should be ignored. In the *focused attention task*, participants were presented with black squares and auditory tones. The squares could either become brighter or keep their color; the tones got quieter or stayed with the same volume. Participants were asked to react only if a square changed its color twice in a row. Changes in the auditory signals were to be ignored. Since we aimed to assess both facets of attention jointly, we computed a compound score termed *crossmodal attention* by averaging mean reaction time in both tests per participant. Thus, higher scores reflect lower performance. Participants who missed both tasks received the sample mean compound score.
2. *Working memory*: Visuo-spatial working memory span was assessed via the computerized version of the Corsi block-tapping task (forward and backward versions) from the Schuhfried Vienna Test System (https://marketplace.schuhfried.com/en/corsi; test forms S1 and S5). Here, participants were presented with a spatial array of nine irregularly arranged cubes on the monitor and observed a cursor that tapped a sequence of cubes. After an

acoustic signal, participants were asked to re-tap the sequence either in the same (forward) or reverse (backward) order. Starting with three block taps, sequence length increased after three runs of a given length up to a maximum of nine taps. We derived a compound score by averaging the number of correctly tapped sequences in both tasks; thus, higher values reflect higher performance. Participants who missed both tasks received the sample mean compound score.

3. *Global task-switching costs:* An alternative-runs task-switching paradigm (Stoet 2010; https://www.psytoolkit.org/experiment-library/taskswitching.html) was used to assess global task-switching abilities. Participants were presented with letter and number combinations appearing sequentially in four quadrants arranged in a 2 $\times$ 2 matrix. When in the top quadrants (Task A), participants had to focus on the letter and press a key to determine whether it was a consonant (G, K, M, R) or vowel (A, E, I, U). When in the bottom quadrants (Task B), participants focused on the number and determined whether it was an odd (3, 5, 7, 9) or even (2, 4, 6, 8) number. Participants carried out “pure blocks” with just one task at a time and “mixed blocks” with both tasks (i.e., two trials of task A, followed by two trials of task B, and then back to task A). Hence, a task switch occurred every two trials, and we obtained information on “task-repeat trials” and “task-switch trials.” Here, we assessed global task-switching costs by subtracting mean RT in single-task trials from that in task-repeat trials, and thus, higher values or costs reflect lower performance. Global task-switching costs (also termed *mixing costs*) are thought to reflect increased processing demands associated with having to maintain and juggle two task configurations in mixed blocks compared to pure blocks (Rogers and Monsell 1995; Wylie and Allport 2000).

### Data analysis

#### *Behavioral data*

We implemented the recently proposed **balanced integration score** (BIS; Liesefeld et al. 2015; Liesefeld and Janczyk 2019) to assess the effects of our experimental manipulations on dual-task costs with a measure that captures both performance facets, speed and accuracy, at the same time and accounts for possible speed–accuracy trade-offs: $BIS_{i,j} = \left(z_{PC_{i,j}} - z_{\overline{RT_{i,j}}}\right) + 100$, with $z_{x_{i,j}} = \frac{x_{i,j} - \bar{x}}{SD_x}$.

The BIS is calculated by first standardizing RT and proportion of correct responses (PC = 100 – ER) to bring them to the same scale (note that the mean and standard deviation must be calculated across all observed RTs and PCs from the analyzed experiment, including all subjects and experimental conditions). In our case, we computed the mean and standard deviation across all observed RTs and PCs based on the final restricted sample of 79 participants after quality control. If values were calculated per age group, both age groups would have a mean value of zero, precluding the analysis of age effects. Therefore, we followed Liesefeld and Janczyk's (2019) suggestion of calculating the mean and sample standard deviations across all conditions and participants. Then, for each subject, the standardized mean RT was subtracted from the corresponding standardized mean PC, and 100 was added to make sure all BIS values were positive. The addition of 100 deviates from the original proposal and is an arbitrary value implemented to facilitate the interpretation of correlations between performance and brain activity.

Higher absolute BIS (> 100) values represent faster and more accurate performance. Analogous to RT, we derived dual-task performance costs by subtracting single- from dual-task BIS values according to the same S-R compatibility level.

Behavioral data

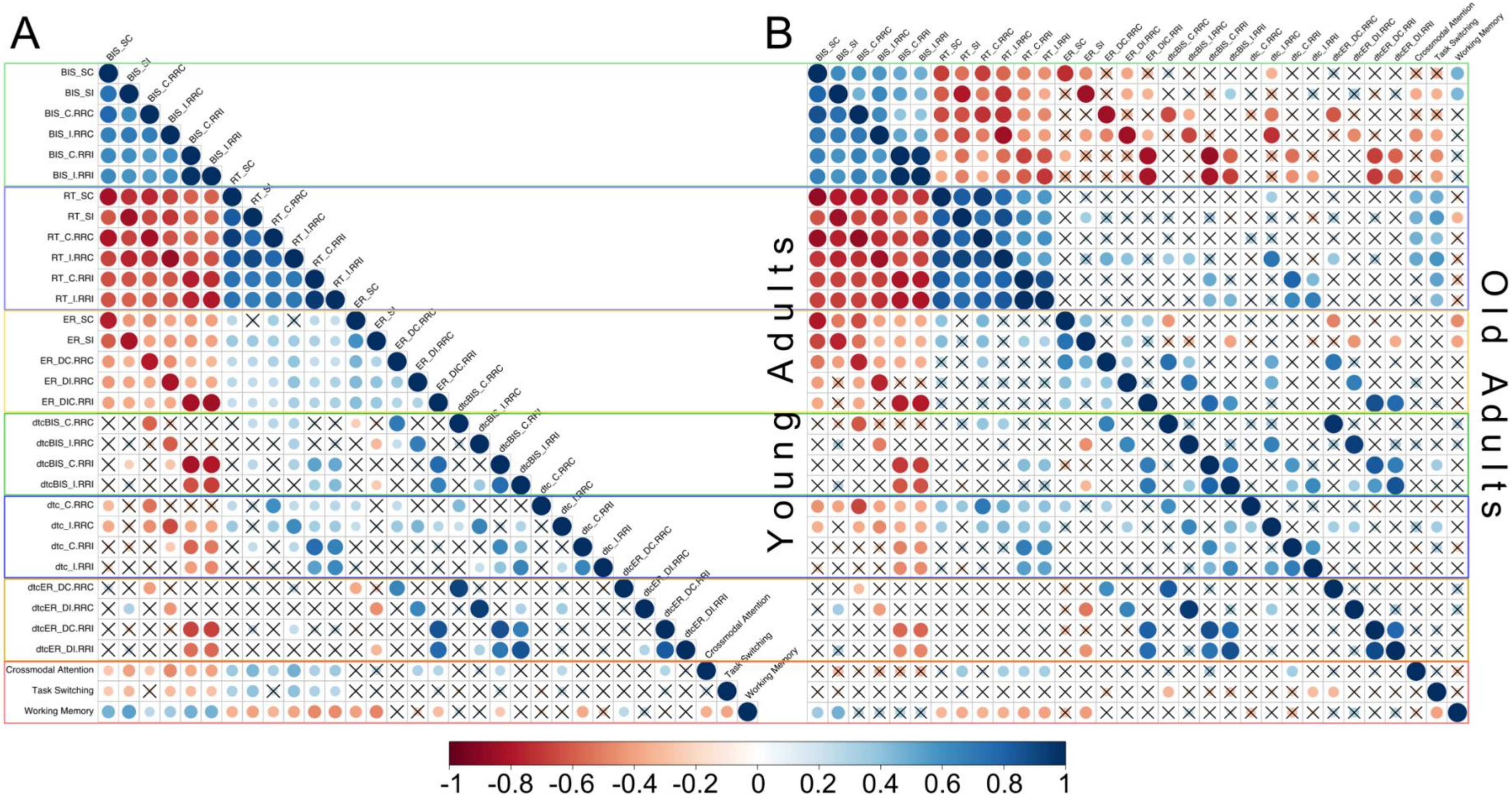

**Figure S1.**

Inter-correlation matrix between behavioral scores for **(A)** all participants and **(B)** per age group (young adults are shown in the lower triangle and older adults in the upper triangle) tested through a Pearson *r* correlation test**.** Non-significant correlations ($p > 0.05$) are crossed out.

**Abbreviations.** BIS: Balanced integration score, C: Stimulus–response compatible, DC: Dual stimulus–response compatible, DI: Dual stimulus–response incompatible, dtc: Dual-task costs, ER: Error rate, I: Stimulus–response incompatible, RRC: Response–response congruent, RRI: Response–response incongruent, RT: Reaction time, SC: Single stimulus–response compatible, SI: Single stimulus–response incompatible.

**Table S1.**

Absolute single- and dual-task performance scores for each age group.

| **Behavioral Score** | **Young (n = 43)** | | **Old (n = 36)** | |
|---|---|---|---|---|
| | ***M*** | ***SD*** | ***M*** | ***SD*** |
| *Balanced integration score* | | | | |
| Single SRC | 101.54 | 1.09 | 100.96 | 0.85 |
| Single SRI | 100.86 | 1.25 | 100.02 | 1.32 |
| Dual SRC-RRC | 100.56 | 1.37 | 100.12 | 1.10 |
| Dual SRI-RRC | 99.84 | 1.38 | 98.78 | 1.73 |
| Dual SRC-RRI | 99.26 | 1.52 | 97.68 | 1.65 |
| Dual SRI-RRI | 99.27 | 1.53 | 97.64 | 1.71 |
| *Reaction time [ms]* | | | | |
| Single SRC | 450.94 | 85.96 | 486.56 | 67.84 |
| Single SRI | 512.13 | 100.72 | 555.19 | 89.09 |
| Dual SRC-RRC | 477.79 | 112.28 | 511.96 | 71.12 |
| Dual SRI-RRC | 551.40 | 111.58 | 616.87 | 122.84 |
| Dual SRC-RRI | 583.55 | 116.79 | 666.70 | 111.93 |
| Dual SRI-RRI | 582.86 | 116.41 | 671.21 | 118.32 |
| *Error rate (%)* | | | | |
| Single SRC | 4.29 | 6.18 | 7.29 | 6.70 |
| Single SRI | 6.01 | 7.45 | 11.08 | 9.23 |
| Dual SRC-RRC | 12.40 | 7.63 | 14.00 | 8.52 |
| Dual SRI-RRC | 13.37 | 8.39 | 18.87 | 11.14 |
| Dual SRC-SRI-RRI | 16.72 | 10.19 | 26.27 | 13.28 |

**Abbreviations.** RRC: Response–response congruent, RRI: Response–response incongruent, SRC: Stimulus–response compatible, SRI: Stimulus–response incompatible**.**

**Table S2.**

Statistical results of the analyses of variance (ANOVA) for dual-task performance costs.

| **Effect/interaction** | ***F*(1,77)** | ***p*** | ***η²*** |
|---|---|---|---|
| *Dual-task costs (ΔBIS)* | | | |
| Age | 6.46 | **0.013** | 0.08 |
| S-R comp | 6.65 | **0.012** | 0.08 |
| R-R congr | 84.29 | **< 0.001** | 0.52 |
| S-R comp × R-R congr | 52.01 | **< 0.001** | 0.40 |
| Age × S-R comp | 0.13 | 0.717 | 0.00 |
| Age × R-R congr | 8.37 | **0.005** | 0.10 |
| Age × S-R comp × R-R congr | 4.31 | **0.041** | 0.05 |
| *Dual-task costs on reaction time (ms)* | | | |
| Age | 7.65 | **0.007** | 0.09 |
| S-R comp | 11.51 | **0.001** | 0.13 |
| R-R congr | 100.93 | **< 0.001** | 0.57 |
| S-R comp × R-R congr | 76.30 | **< 0.001** | 0.50 |
| Age × S-R comp | 0.90 | 0.345 | 0.01 |
| Age × R-R congr | 4.35 | **0.040** | 0.05 |
| Age × S-R comp × R-R congr | 1.70 | 0.196 | 0.02 |
| *Dual-task costs on error rate (%)* | | | |
| Age | 2.41 | 0.125 | 0.03 |
| S-R comp | 1.91 | 0.171 | 0.02 |
| R-R congr | 25.87 | **< 0.001** | 0.25 |
| S-R comp × R-R congr | 6.43 | **0.013** | 0.08 |
| Age × S-R comp | 0.00 | 0.947 | 0.00 |
| Age × R-R congr | 5.01 | **0.028** | 0.06 |
| Age × S-R comp × R-R congr | 2.86 | 0.095 | 0.04 |

**Abbreviations.** S-R comp: Stimulus–response compatibility, R-R congr: Response–response congruency.

**Note.** Significant *p*-values are shown in bold.

*Generalized slowing*

Since it has often been suggested that older adults' performance may decrease because information is processed more slowly overall, we tested whether the age-related dual-task deficits were a task-specific process or were only explained by generalized cognitive slowing (Salthouse 1996). We implemented the Brinley procedure (Brinley 1965; Madden et al. 1992; Cerella 1994), which applies a linear model (i.e., the Brinley function) and corrects for global age-related differences in RT across task conditions. The assumption is that if the interactions between experimental factors and age remain statistically significant after the procedure, there is strong evidence for domain-specific age differences independent of general age differences in processing speed.

Following the Brinley procedure (Brinley 1965; Madden et al. 1992; Cerella 1994), we first regressed the old adults' group-averaged RT values of all six single and dual experimental conditions on the corresponding mean values from younger adults (see **Figure S2**). Old adult's RTs were best represented by the following Brinley function: $RT_{OA} = 1.42 \times (RT_{YA}) - 161.26$ ($R^2 = 0.99$). Then, we implemented the Brinley function on the younger adult's RT to generate "old-like" RTs. After that, we recalculated the statistical analyses for dual-task speed costs with the resulting values (i.e., the transformed RT values for the young group and untransformed ones for the old group). As before, we computed the difference in mean RT between dual and single conditions of the same S-R compatibility to obtain the dual-task costs for each participant.

The RT transformation yielded "old-like" average dual-task speed costs of 95.46 ± 103.24 ms for young adults. As the (untransformed) average costs for older adults were 95.81 ± 95.50 ms, the age effect was not significant anymore, $F(1,77) = 0.00$, $p = 0.978$, $\eta_p^2 = 0.00$ (see **Table S3** for descriptive and statistical results after transformation). The mixed three-way ANOVA again revealed significant main effects of S-R compatibility, $F(1,77) = 12.72$, $p = 0.001$, $\eta_p^2 = 0.14$, and R-R congruency, $F(1, 77) = 98.36$, $p < 0.001$, $\eta_p^2 = 0.56$, qualified by a significant interaction between S-R compatibility and R-R congruency, $F(1,77)$

= 84.19, $p$ < 0.001, $\eta_p^2$ = 0.52, as in all performance scores. However, the previously observed interaction between R-R congruency and age turned insignificant ($p$ = 0.720).

Based on this analysis, an alternative explanation for the observed age differences in response-code conflict is that elderly suffer from global cognitive slowing; however, other mechanisms could also explain the differences (e.g., response grouping, elaborated in the next section).

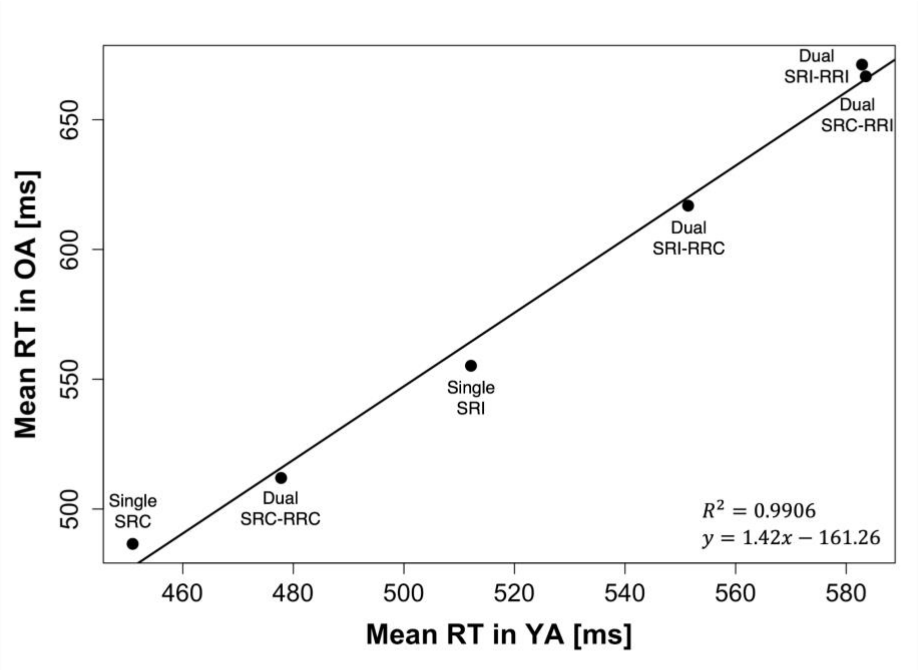


**Figure S2.**

Brinley plot illustrating single and dual mean reaction time of older adults as a function of the corresponding mean reaction time of younger adults for each experimental condition.

**Abbreviations.** SRC: Stimulus–response compatible, SRI: Stimulus–response incompatible, RRC: Response–response congruent, RRI: Response–response incongruent.

**Table S3.**

Descriptive and statistical results of the analysis of variance (ANOVA) for dual-task speed costs after Brinley procedure to assess task-dependent and generalized slowing effects.

| Behavioral Score | Young (n = 43) | | Old (n = 36) | | Effect/interaction | | | |
|---|---|---|---|---|---|---|---|---|
| | *M* | *SD* | *M* | *SD* | | $F(1,77)$ | $p$ | $\eta^2$ |
| *Dual-task costs on reaction time (ms)* | | | | | *Dual-task costs on reaction time (ms)* | | | |
| SRC-RRC | 38.04 | 65.30 | 25.39 | 26.28 | Age | 0.00 | 0.978 | 0.00 |
| SRI-RRC | 55.65 | 62.20 | 61.68 | 62.29 | S-R comp | 12.72 | **0.001** | 0.14 |
| SRC-RRI | 187.90 | 96.45 | 180.13 | 92.16 | R-R congr | 98.36 | **< 0.001** | 0.56 |
| SRI-RRI | 100.22 | 110.40 | 116.02 | 100.82 | S-R comp × R-R congr | 84.19 | **< 0.001** | 0.52 |
| | | | | | Age × S-R comp | 2.37 | 0.128 | 0.03 |
| | | | | | Age × R-R congr | 0.13 | 0.720 | 0.00 |
| | | | | | Age × S-R comp × R-R congr | 0.05 | 0.828 | 0.00 |

**Abbreviations.** RRC: Response–response congruent, RRI: Response–response incongruent, R-R congr: Response–response congruency, SRC: Stimulus–response compatible, SRI: Stimulus–response incompatible, S-R comp: Stimulus–response compatibility.

**Note.** Significant *p*-values are shown in bold.

*Response grouping*

Our experimental design might promote synchronized (or *response grouping*) because the task design involves two concurrent manual choice responses. Response grouping refers to a strategy in dual-task settings by which the faster of two reaction processes is slowed by synchronizing its motor execution with that of the second, slower reaction (Pashler 1994). This strategy would not compromise the findings of increased dual-task costs with incongruent response codes and the enhancement of this effect in advanced age, but we implemented the procedure proposed by (Miller & Ulrich, 2008) to account for possible grouping effects. The assumption was that if the observed effects remained intact after removing strongly synchronized responses, it would indicate that, at least in the remaining trials, participants engaged a mechanism different from response grouping to cope with crosstalk under conditions of response-code conflict.

The procedure involves creating a cumulative frequency distribution (CDF) of inter-response intervals (IRI) to establish an IRI cut-off value of response-grouped trials on the spot, on which a reduction in the slope of the CDF for the S-R compatible, R-R congruent trials becomes apparent. **Figure S3** shows the CDFs of IRIs categorized in deciles as a function of dual condition. The CDF shows a narrow distribution with a marked slope reduction at 90% of the trials with an IRI of 48 ms, indicating this value is an appropriate cut-off value for identifying grouped responses in our dual setting. Furthermore, approximately 80% of IRIs in R-R incongruent trials were shorter than 48 ms, and the resting trials showed longer IRIs spread out over a wide range (48 – 952 ms), consistent with the idea that our cut-off value leaves non-grouped responses. Of note, from the R-R incongruent conditions with correct trials, in 48.79 % of the trials, the S-R compatible response was given before the incompatible (47.47 % for young and 50.37 % for older adults); and in 51.17 %, the incompatible was given before the compatible one (52.47 % for young and 49.63 % for older adults).

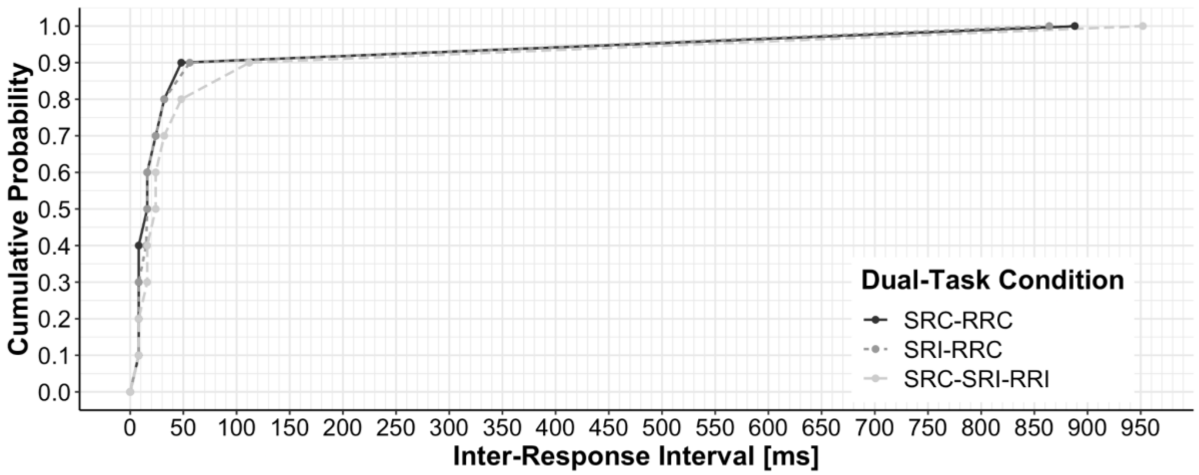


**Figure S3.**

Cumulative frequency distributions of inter-response intervals (IRIs) categorized in deciles across all subjects as a function of dual-task condition. **Abbreviations.** SRC: Stimulus–response compatible, SRI: Stimulus–response incompatible, RRC: Response–response congruent, RRI: Response–response incongruent.

After excluding all trials with grouped responses (IRI < 48 ms), we were left with 37 young and 34 older adults because not all participants had enough non-grouped responses in each dual conditions. When recalculating the dual-task costs on RT in this sample and repeating the mixed three-way ANOVA (see **Figure S4** and **Table S4** for the detailed statistical results), we obtained the same pattern of results as before, except for the age-related effects, i.e., age main effect ($p$ = 0.128) and interaction between age and R-R congruency ($p$ = 0.089), which then turned non-significant.

We found significant main effects of S-R compatibility ($F$(1, 69) = 4.79, $p$ = 0.032, $\eta_p^2$ = 0.06) and R-R congruency ($F$(1, 69) = 34.22, $p$ < 0.001, $\eta_p^2$ = 0.33) on dual-task speed costs. These main effects were again overruled by a significant interaction between S-R compatibility and R-R congruency ($F$(1, 69) = 8.49, $p$ = 0.005, $\eta_p^2$ = 0.11). In more detail, the S-R compatibility reversal effect with R-R incongruency survived as well: Dual-task speed costs did not differ between S-R compatible and incompatible responses with R-R congruency ($p$ = 0.603), but when response codes for either hand were incongruent to each other, dual-task costs were significantly higher for S-R *compatible* responses (225.25 ± 111.81 ms) than for *incompatible* ones (160.69 ± 123.75 ms, $p$ < 0.001). Finally, as before, we neither found a significant interaction between S-R compatibility and age nor a three-way interaction between S-R compatibility, R-R congruency, and age.

Our results support the idea that participants engaged another mechanism different from response grouping to cope with crosstalk under conditions of response-code conflict. However, it is possible that older adults, besides suffering from generalized cognitive slowing, additionally implemented the alternative strategy of response grouping to cope with conflicting response codes.

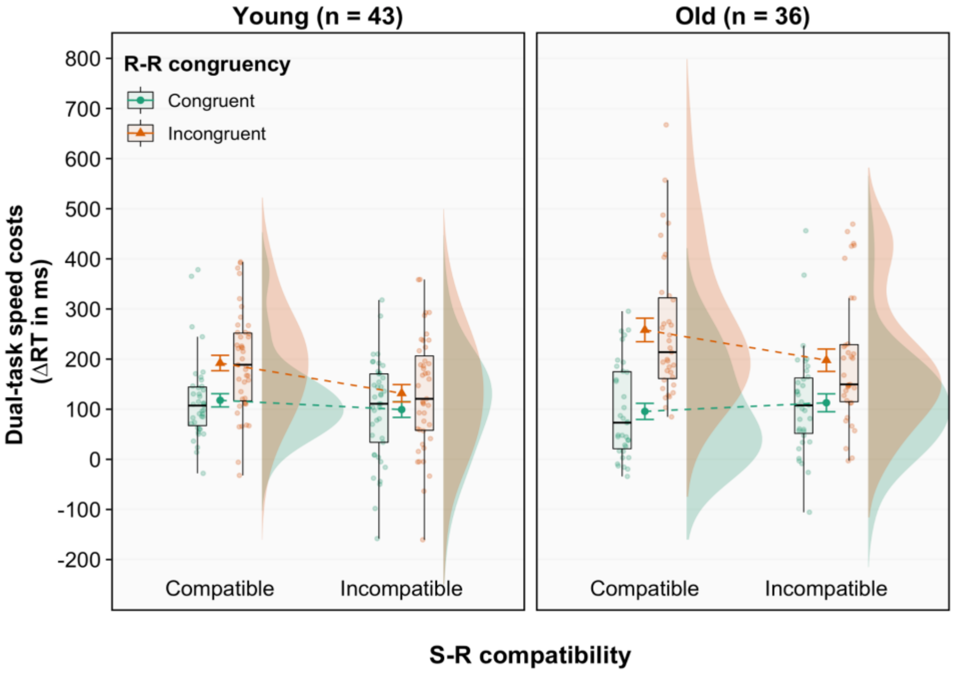


**Figure S4.**

Dual-task costs on reaction time excluding response-grouped trials (cut-off value of inter-response interval ≥ 48 ms) according to age group, stimulus–response compatibility, and response–response congruency. Dual-task costs were obtained through the difference in mean RT between analogous dual- and single-task conditions. Error bars represent S.E.M.

**Abbreviations.** RT: Reaction time, S-R: Stimulus–response, R-R: Response–response.

**Table S4.**

Descriptive and statistical results of the analysis of variance (ANOVA) for dual-task speed costs after excluding response-grouped trials (cut-off value of inter-response interval ≥ 48 ms).

| Behavioral Score | Young (n = 37) | | Old (n = 34) | | Effect/interaction | | | |
|---|---|---|---|---|---|---|---|---|
| | *M* | *SD* | *M* | *SD* | | $F(1,69)$ | $p$ | $\eta^2$ |
| *Dual-task costs on reaction time (ms)* | | | | | *Dual-task costs on reaction time (ms)* | | | |
| SRC-RRC | 117.78 | 84.91 | 91.09 | 92.60 | Age | 2.38 | 0.128 | 0.03 |
| SRI-RRC | 99.78 | 102.57 | 130.67 | 187.65 | S-R comp | 4.79 | **0.032** | 0.06 |
| SRC-RRI | 204.44 | 102.67 | 246.06 | 120.95 | R-R congr | 34.22 | **< 0.001** | 0.33 |
| SRI-RRI | 130.39 | 119.57 | 190.99 | 127.93 | S-R comp × R-R congr | 8.49 | **0.005** | 0.11 |
| | | | | | Age × S-R comp | 2.43 | 0.124 | 0.03 |
| | | | | | Age × R-R congr | 2.97 | 0.089 | 0.04 |
| | | | | | Age × S-R comp × R-R congr | 0.56 | 0.458 | 0.01 |

**Abbreviations.** RRC: Response–response congruent, RRI: Response–response incongruent, R-R congr: Response–response congruency, SRC: Stimulus–response compatible, SRI: Stimulus–response incompatible, S-R comp: Stimulus–response compatibility.

**Note.** Significant *p*-values are shown in bold.

### Neuroimaging data

##### Table S5.

Brain regions showing significant effects for each task-specific contrast across both age groups.

| ***Contrast /*** **Macroanatomical structure** | **H** | **Cluster extent ($k_E$)** | **Cluster *p(cFWE)*** | **MNI coordinates** *x* | *y* | *z* | **Cytoarchit. assignm. (Overlap in %)** | **Peak *Z* value** |
|---|---|---|---|---|---|---|---|---|
| ***Dual$_{RRC}$ + > Single*** | | | | | | | | |
| Precentral gyrus | R | 13223 | < 0.001 | 40 | -14 | 56 | Area 4a (15.2) | > 8 |
| | L | | | -40 | -18 | 60 | Area 4a (17.8) | > 8 |
| Postcentral gyrus | R | | | 52 | -18 | 48 | Area 2 (49.0) | > 8 |
| | L | | | -50 | -22 | 52 | Area 2 (22.1) | > 8 |
| | R | | | 44 | -32 | 60 | Area 1 (45.5) | > 8 |
| Supplementary motor area | R | | | 2 | -8 | 52 | Area 6mc / SMA (53.6) | > 8 |
| | L | | | -2 | -10 | 52 | Area 6mc / SMA (57.5) | > 8 |
| Cerebellum VI | L | 3150 | < 0.001 | -18 | -54 | -22 | – | > 8 |
| | R | | | 22 | -52 | -24 | – | > 8 |
| Cerebellum V | L | | | -4 | -60 | -14 | – | > 8 |
| Thalamus | R | 1711 | < 0.001 | 16 | -14 | 4 | – | 7.81 |
| Putamen | R | | | 26 | -8 | 0 | – | 6.38 |
| Pallidum | R | | | 22 | 0 | -6 | – | 6.00 |
| Central opercular cortex | R | | | 46 | 4 | 8 | Area OP8 (14.1) | 5.85 |
| Inferior frontal gyrus, pars op. | R | | | 48 | 6 | 10 | Area 44 (10.7) | 5.82 |
| Central insula | R | | | 34 | 4 | 4 | – | 5.09 |
| Thalamus | L | 855 | < 0.001 | -14 | -18 | 4 | – | > 8 |
| Putamen | L | | | -28 | -6 | -2 | – | 5.71 |
| ***Dual$_{RRC}$ – < Single*** | | | | | | | | |
| Occipital pole | R | 2902 | < 0.001 | 18 | -96 | 0 | Area hOc1 (V1, 75.1) | > 8 |
| | R | | | 26 | -96 | 18 | Area hOc3d (V3d, 54.7) | 7.38 |
| Lateral occipital cortex, inf. div. | R | | | 44 | -74 | -6 | Area hOc4la (74.5) | 7.22 |
| | R | | | 38 | -90 | 6 | Area hOc4lp (79.6) | 6.30 |
| Occipital fusiform gyrus | R | | | 30 | -80 | -6 | Area hOc4v (V4v, 62.5) | 5.53 |
| Frontal pole | L/R | 1884 | < 0.001 | 0 | 60 | 12 | – | 5.46 |

| | | | | | | | | |
|---|---|---|---|---|---|---|---|---|
| Cingulate gyrus, ant. div. | L/R | | | 0 | 42 | 2 | – | 4.68 |
| Frontal pole | R | | | 2 | 58 | -2 | – | 4.44 |
| Paracingulate gyrus | R | | | 2 | 44 | 22 | – | 4.38 |
| | L | | | -8 | 40 | -10 | Area s32 (45.4) | 4.32 |
| Frontal medial cortex | R | | | 2 | 40 | -18 | Area s32 (39.4) | 4.23 |
| | L | | | -4 | 48 | -20 | Area Fp2 (39.7) | 3.89 |
| Occipital pole | L | 1876 | < 0.001 | -12 | -92 | -2 | Area hOc1 (V1, 62.7) | 7.06 |
| | L | | | -16 | -100 | 14 | Area hOc2 (V2, 48.3) | 6.49 |
| Occipital fusiform gyrus | L | | | -38 | -76 | -10 | Area FG1 (38.8) | 5.31 |
| | L | | | -32 | -80 | -8 | Area hOc4v (V4v, 42.4) | 5.00 |
| Occipital pole | L | | | -28 | -94 | 22 | Area hOc4d (V3A, 39.7) | 4.92 |
| Lateral occipital cortex, inf. div. | L | | | -28 | -90 | 4 | Area hOc4lp (67.9) | 4.68 |
| Occipital pole | L | | | -10 | -96 | 28 | Area hOc3d (V3d, 62.4) | 4.30 |
| Hippocampus | R | 424 | 0.003 | 34 | -30 | -10 | CA1 (Hippocampus, 60.7) | 4.69 |
| | R | | | 30 | -14 | -20 | DG (Hippocampus, 47.0) | 4.43 |
| Superior frontal gyrus | R | 384 | 0.006 | 2 | 24 | 60 | Area 6mr / preSMA (2.7) | 5.12 |
| | L | | | -2 | 36 | 56 | – | 4.97 |
| Frontal pole | R | | | 8 | 42 | 52 | – | 4.36 |
| Superior frontal gyrus | L | | | -14 | 32 | 54 | – | 3.54 |
| Frontal pole | R | | | 18 | 38 | 54 | – | 3.50 |
| Angular gyrus | L | 229 | 0.045 | -48 | -62 | 50 | Area PFm (IPL, 40.7) | 5.38 |
| Lateral occipital cortex, sup. div. | L | | | -42 | -78 | 38 | Area PGp (IPL, 55.1) | 4.08 |
| | L | | | -32 | -80 | 44 | Area hIP5 (IPS, 45.4) | 3.86 |
| | L | | | -42 | -64 | 54 | Area PGa (IPL, 39.0) | 3.64 |
| Supramarginal gyrus, post. div. | L | | | -60 | -50 | 42 | Area PFm (IPL, 51.0) | 3.25 |
| ***Dual$_{RRI}$ + > Single*** | | | | | | | | |
| Precentral gyrus | R | 52398 | < 0.001 | 30 | -10 | 60 | Area 6dl (31.7) | > 8 |
| | R | | | 40 | -16 | 54 | Area 3b (19.3) | > 8 |
| Cerebellum VI | L | | | -20 | -54 | -22 | – | > 8 |
| Postcentral gyrus | R | | | 44 | -36 | 58 | – | > 8 |
| Superior parietal lobule | R | | | 42 | -38 | 56 | Area 2 (25.0) | > 8 |
| *This cluster covers large portions of frontal, motor, somatosensory, cingulate, insular, and parietal cortices on both hemispheres, as well as basal ganglia and cerebellum. However, due to its size, peaks do not become significant.* | | | | | | | | |
| Middle temporal gyrus, TO part | R | 316 | 0.014 | 54 | -52 | 0 | – | 4.79 |
| Inferior temporal gyrus, TO part | R | | | 60 | -54 | -14 | – | 4.37 |

| | | | | | | | | |
|---|---|---|---|---|---|---|---|---|
| ***$Dual_{RRI}$ – < Single*** | | | | | | | | |
| Occipital pole | R | 278 | 0.023 | 24 | -98 | 16 | Area hOC2 (V2, 49.0) | 6.37 |
| | R | | | 28 | -98 | 10 | Area hOc3d (V3d, 40.8) | 5.79 |
| Lateral occipital cortex, inf. div. | R | | | 44 | -76 | -8 | Area hOc4la (64.2) | 4.45 |
| Occipital pole | R | | | 42 | -90 | 4 | Area hOc4lp (82.9) | 4.14 |
| Frontal medial cortex | L | 251 | 0.033 | -6 | 46 | -18 | Area Fp2 (19.0) | 5.87 |
| | R | | | 4 | 42 | -20 | Area s32 (3.4) | 5.21 |
| | L | | | -6 | 42 | -16 | Area s32 (44.2) | 5.14 |
| Paracingulate gyrus | R | | | 2 | 32 | -12 | Area s24 (40.3) | 4.21 |
| | L | | | -8 | 32 | -10 | Area s24 (47.9) | 3.56 |
| ***$Dual_{RRI}$ + > $Dual_{RRC}$*** | | | | | | | | |
| Superior frontal gyrus | R | 37326 | < 0.001 | 24 | -6 | 54 | Area 6d3 (70.5) | > 8 |
| Superior parietal lobule | R | | | 20 | -60 | 62 | Area 7A (SPL; 68.6) | > 8 |
| | R | | | 32 | -42 | 40 | Area hIP1 (IPS; 38.4) | > 8 |
| Superior frontal gyrus | L | | | -24 | -2 | 58 | Area 6d3 (41.3) | > 8 |
| Insular cortex | R | | | 32 | 22 | -8 | – | > 8 |
| | L | | | -32 | 22 | -6 | Area Id7 (3.8) | > 8 |
| Thalamus | R | | | 18 | -22 | 8 | – | > 8 |
| Cerebellum | L | | | -12 | -72 | -24 | – | 7.61 |
| Superior parietal lobule | L | 5255 | < 0.001 | -20 | -62 | 60 | Area 7A (SPL; 56.2) | > 8 |
| | L | | | -36 | -44 | 46 | Area hIP3 (IPS, 20.8) | > 8 |
| Supramarginal gyrus, ant. div. | L | | | -56 | -30 | 34 | Area PFt (IPL; 62.2) | 4.38 |
| Middle temporal gyrus, TO part | R | 375 | 0.006 | 54 | -52 | 0 | – | 4.79 |
| Inferior temporal gyrus, TO part | R | | | 60 | -54 | -14 | – | 4.37 |
| ***$Dual_{RRI}$ – < $Dual_{RRC}$*** | | | | | | | | |
| *nothing significant* | | | | | | | | |

**Abbreviations.** Ant. div.: Anterior division, CA: Cornu ammonis, Cytoarchit. assignm.: Cytoarchitectonic assignment, DG: Dentate gyrus, FG: Fusiform gyrus, FP: Frontal pole, inf. div.: Inferior division, IPL: Inferior parietal lobule, IPS: Intraparietal sulcus, Oc: Occipital cortex, OP: Opercular cortex, PFm: Parietal area F, part m, PFt: Parietal area F, part t, PGp: Parietal area G, posterior, post. div.: Posterior division, preSMA: Presupplementary motor area, RRC: Response–response congruent, RRI: Response–response incongruent, SFG: Superior frontal gyrus, SMA: Supplementary motor area, SPL: Superior parietal lobule, sup. div.: Superior division, TO: Temporooccipital.

**Note.** Selection of peaks based on macroanatomical and cytoarchitectonic representative regions.

The contrasts were estimated via a conjunction combining the contrast and the regressor's main effect, marked with + or –.

All activations are reported at a cluster-forming threshold with voxel-level uncorrected at $p < 0.001$, and cluster-level family-wise error (cFWE) corrected at $p < 0.05$.

Detailed information on the cytoarchitectonic maps for all regions included in all supplementary tables can be found in the respective publications: Areas 4a and 4p (Geyer et al. 1996); Area 2 (Grefkes et al. 2001); Areas 1, 3a, 3b (Geyer et al. 1999; Geyer et al. 2000); Area 6mc / SMA and 6mr /preSMA (Ruan et al. 2018); Areas 44 and 45 (Amunts et al. 1999; Amunts et al. 2004); Areas hOc1 (V1) and hOc2 (V2) (Amunts et al. 2000); Areas hOc3d (V3d) and hOc4d (V3a) (Kujovic et al. 2013); Areas hOc4la and hOc4lp (Malikovic et al. 2016); Areas hOc3v (V3v) and hOc4v (V4v) (Rottschy et al. 2007), Areas 33, s24 and s32 (Palomero-Gallagher et al. 2015); Areas Fp1 and Fp2 (Bludau et al. 2014); Area FG1 (Caspers et al. 2013); Areas CA1 and DG (Hippocampus) (Amunts et al. 2005); Areas PF, PFcm, PFm, PFop, PFt, PGa and PGp (IPL) (Caspers et al. 2006; Caspers et al. 2008); Areas hIP5, hIP6 and hIP8 (IPS) (Richter et al. 2019); Area 6 (Geyer 2004); Areas 5Ci, 5L, 5M, 7A, 7P and 7PC (SPL) and hIP3 (IPS) (Scheperjans, Hermann, et al. 2008; Scheperjans, Eickhoff, et al. 2008); Areas hIP1 and hIP2 (IPS) (Choi et al. 2006); Areas OP3 (VS) (Eickhoff, Amunts, et al. 2006; Eickhoff, Schleicher, et al. 2006); Area Ig1 (Kurth et al. 2010); Areas p24ab, p24c and p32 (Palomero-Gallagher et al. 2019); Area Fo3 (Henssen et al. 2016); Area TE 3 (Morosan et al. 2005); Areas FG3 and FG4 (Lorenz et al. 2015); Ventral dentate nucleus and interposed nucleus (Tellmann et al. 2015).

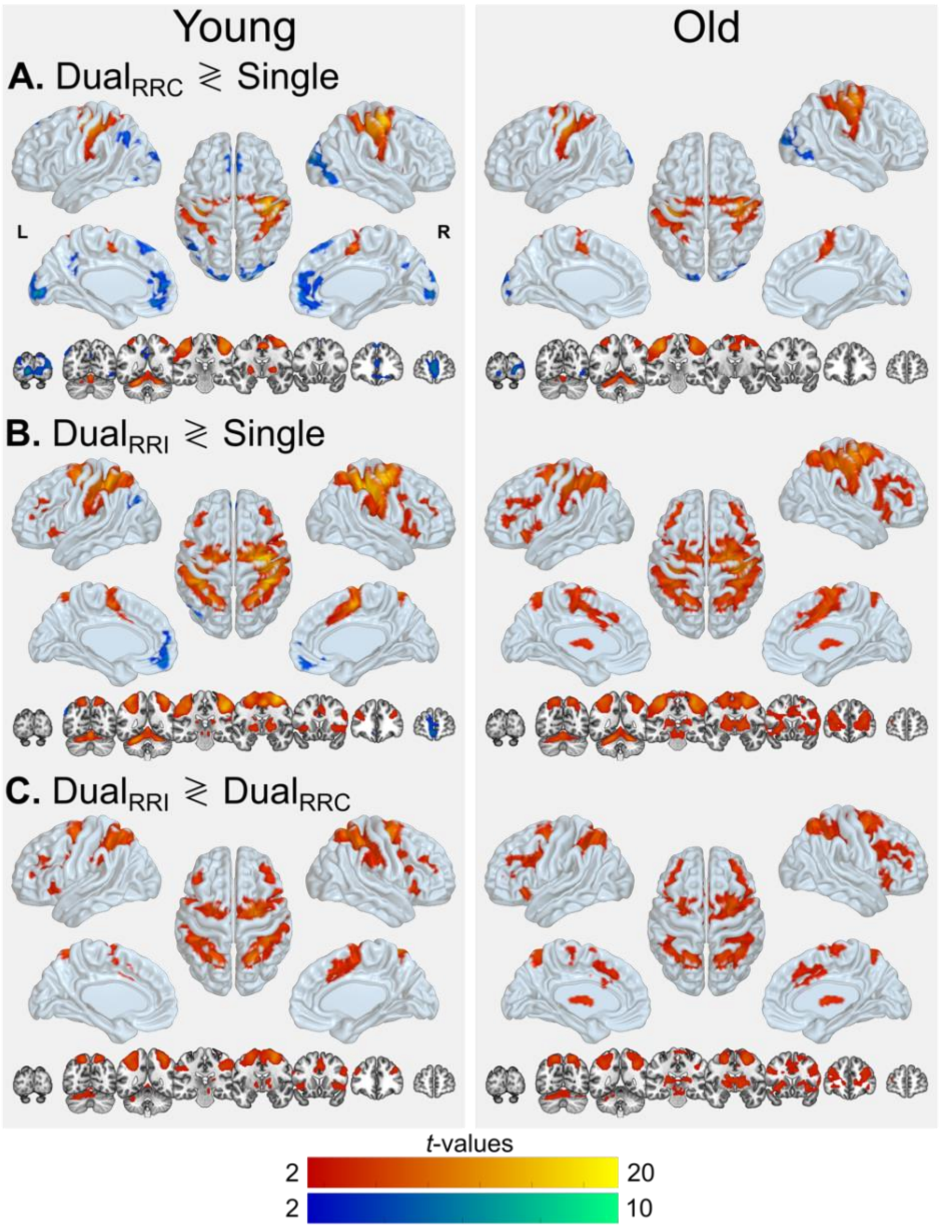


**Figure S5.**

Task brain activity for young and older adults separately (left and right column, respectively). **(A)** Dual-response execution network obtained by contrasting dual R-R congruent vs. single-task conditions (Dual$_{RRC}$ ≷ Single). **(B)** Dual-task network obtained by contrasting dual R-R incongruent vs. single-task conditions (Dual$_{RRI}$ ≷ Single). **(C)** Dual-task response-code conflict network computed by contrasting dual-task R-R incongruent vs. dual R-R congruent conditions (Dual$_{RRI}$ ≷ Dual$_{RRC}$). Statistical interaction between the dual-task network and age. Hot colors represent activations (i.e., “greater than” contrasts), and cool colors represent deactivations (i.e., “smaller than” contrasts). All effects were significant at cluster-level $p < 0.05$ (cFWE-corrected) with a cluster-forming threshold of $p < 0.001$ at voxel level. **Abbreviations.** RRC: Response–response congruent, RRI: Response–response incongruent.

Examining each age group separately for the effect of dual-response execution versus single-hand responding (see **Supplementary Figure S5A**), young adults showed pretty much the same activation and deactivation patterns as were observed for the entire sample (cf. **Figure 3A**), but some deactivations were more pronounced in bilateral medial prefrontal cortex and ACC. Older adults, in turn, showed a similar but overall weaker activation pattern and some sparse deactivation in occipital cortex.

The conceptually wide contrast of incongruent dual-tasking (vs. single trials; see **Supplementary Figure S5B**) also covered almost the same brain regions as for the entire sample (cf. **Figure 3B**), but only the young adults presented reduced activity in bilateral medial prefrontal cortex and paracingulate gyrus, as well as left IPL, while the older group did not present any deactivations. Older adults, in contrast, showed again overall weaker activity but covering wider areas of bilateral prefrontal cortex, SMA, preSMA and basal ganglia.

The activation pattern linked to dual-task response-code conflict also looked rather similar when separately analyzed in either age group (see **Supplementary Figure S5C**); older adults only appeared to present wider activations in the basal ganglia. No significant deactivations were found. The **Supplementary Table S6** lists the local maxima of clusters showing increased and decreased brain activity for each task-specific contrast, separately for young and older adults.

**Table S6.**

Brain regions showing significant effects for each task-specific contrast in each age group.

| ***Contrast /*** **Macroanatomical structure** | **H** | **Cluster extent ($k_E$)** | **Cluster *p(cFWE)*** | **MNI coordinates x** | **y** | **z** | **Cytoarchit. assignm. (Overlap in %)** | **Peak *Z* value** |
|---|---|---|---|---|---|---|---|---|
| ***YA: Dual$_{RRC}$ + > Single*** | | | | | | | | |
| Precentral gyrus | R | 9340 | < 0.001 | 40 | -14 | 58 | Area 3b (10.6) | > 8 |
| Postcentral gyrus | R | | | 50 | -18 | 48 | Area 2 (51.1) | > 8 |
| Precentral gyrus | L | | | -38 | -16 | 60 | Area 6d1 (20.7) | > 8 |
| Postcentral gyrus | R | | | 46 | -34 | 58 | Area 1 (51.3) | > 8 |
| | L | | | -50 | -22 | 52 | Area 2 (22.1) | > 8 |
| | L | | | -38 | -28 | 48 | Area 4p (47.1) | > 8 |
| Supplementary motor area | L/R | | | 4 | -6 | 52 | Area 6mc / SMA (46.3) | > 8 |
| Supramarginal gyrus, ant. div. | R | | | 54 | -18 | 24 | Area PFop (IPL, 31.9) | 5.91 |
| Superior parietal lobule | L | | | -28 | -54 | 64 | Area 7PC (SPL, 42.3) | 4.81 |
| Central opercular cortex | L | | | -48 | -18 | 16 | Area OP3 (VS, 36.2) | 4.80 |
| Cerebellum VI | L | 2673 | < 0.001 | -22 | -54 | -24 | – | > 8 |
| | R | | | 22 | -54 | -24 | – | > 8 |
| Cerebellum V | L | | | -2 | -62 | -16 | – | > 8 |
| Thalamus | R | 1004 | < 0.001 | 16 | -14 | 6 | – | 7.06 |
| Pallidum | R | | | 26 | -4 | 0 | – | 6.79 |
| Central opercular cortex | R | | | 44 | 2 | 10 | Area OP3 (VS, 3.6) | 4.04 |
| Precentral gyrus | R | | | 50 | 6 | 10 | Area 44 (25.7) | 4.04 |
| Insular cortex | R | | | 32 | -22 | 4 | Area Ig1 (47.2) | 3.93 |
| Central opercular cortex | R | | | 48 | 4 | 2 | Area OP8 (20.2) | 3.57 |
| Putamen | L | 641 | < 0.001 | -26 | -6 | 0 | – | 6.30 |
| Thalamus | L | | | -14 | -18 | 6 | – | 6.21 |
| Putamen | L | | | -26 | -18 | 4 | – | 3.34 |
| ***OA: Dual$_{RRC}$ + > Single*** | | | | | | | | |
| Precentral gyrus | L | 11355 | < 0.001 | -36 | -24 | 50 | Area 4p (64.6) | > 8 |
| | R | | | 40 | -14 | 54 | Area 3b (18.0) | > 8 |
| | L | | | -42 | -20 | 60 | Area 4a (26.5) | > 8 |
| | R | | | 38 | -20 | 64 | Area 4a (8.3) | > 8 |
| | R | | | 34 | -24 | 56 | Area 4p (57.7) | > 8 |

| | | | | | | | | |
|---|---|---|---|---|---|---|---|---|
| | L | | | -34 | -20 | 66 | Area 6d1 (18.1) | > 8 |
| Postcentral gyrus | L | | | -50 | -20 | 50 | Area 3b (37.5) | > 8 |
| | R | | | 54 | -18 | 46 | Area 2 (44.8) | > 8 |
| Supplementary motor area | L/R | | | -4 | -10 | 50 | Area 6mc / SMA (33.8) | > 8 |
| Cerebellum V | L | 2283 | < 0.001 | -16 | -54 | -20 | – | > 8 |
| | R | | | 16 | -52 | -20 | – | > 8 |
| | L | | | -6 | -56 | -12 | – | > 8 |
| ***YA: Dual$_{RRC}$ – < Single*** | | | | | | | | |
| Occipital pole | L | 3489 | < 0.001 | -12 | -92 | 0 | Area hOc1 (V1, 72.7) | 6.88 |
| | R | | | 16 | -98 | 2 | Area hOc1 (V1, 88.4) | 6.31 |
| Lateral occipital cortex, inf. div. | R | | | 38 | -88 | 8 | Area hOc4lp (92.6) | 5.88 |
| Occipital pole | R | | | 20 | -98 | 18 | Area hOc2 (V2, 52.5) | 5.72 |
| | R | | | 26 | -96 | 16 | Area hOc3d (V3d, 55.6) | 5.44 |
| Lateral occipital cortex, inf. div. | R | | | 46 | -78 | -4 | Area hOc4la (92.0) | 5.33 |
| Occipital fusiform gyrus | R | | | 30 | -80 | -6 | Area hOc4v (V4v, 62.5) | 5.24 |
| Occipital pole | L | | | -14 | -100 | 16 | Area hOc2 (V2, 47.9) | 4.94 |
| | L | | | -28 | -94 | 22 | Area hOc4d (V3A, 39.7) | 4.36 |
| Frontal pole | L/R | 2590 | < 0.001 | 0 | 60 | 10 | – | 5.81 |
| | R | | | 4 | 56 | -2 | Area Fp2 (58.9) | 5.67 |
| Paracingulate gyrus | L | | | -6 | 42 | -10 | Area p32 (43.1) | 5.43 |
| Cingulate gyrus, ant. div. | L/R | | | 0 | 42 | 2 | – | 4.97 |
| | R | | | 2 | 36 | 16 | Area p24ab (54.4) | 4.39 |
| Frontal medial cortex | R | | | 2 | 40 | -18 | Area s32 (39.4) | 4.32 |
| Frontal pole | R | | | 22 | 36 | -14 | Area Fo3 (75.8) | 4.03 |
| Superior frontal gyrus | L/R | 404 | 0.004 | 0 | 38 | 54 | – | 5.50 |
| | R | | | 2 | 24 | 60 | Area 6mr / preSMA (2.7) | 5.03 |
| | L | | | -2 | 28 | 50 | – | 4.07 |
| Inferior parietal lobule | L | 311 | 0.014 | -50 | -64 | 46 | Area PGa (IPL, 52.3) | 4.91 |
| | L | | | -40 | -78 | 40 | Area PGp (IPL, 48.5) | 3.96 |
| Inferior parietal sulcus | L | | | -32 | -78 | 46 | Area hIP5 (IPS, 53.1) | 3.47 |
| Supramarginal gyrus, post. div. | L | | | -60 | -52 | 40 | Area PFm (IPL, 54.5) | 3.38 |
| ***OA: Dual$_{RRC}$ – < Single*** | | | | | | | | |
| Occipital pole | R | 1083 | < 0.001 | 20 | -92 | 2 | Area hOc1 (V1, 36.6) | 6.07 |
| | R | | | 26 | -96 | 18 | Area hOc3d (V3d, 54.7) | 5.25 |
| Lateral occipital cortex, inf. div. | R | | | 42 | -72 | -2 | Area hOc4la (61.3) | 4.93 |

| | | | | | | | | |
|---|---|---|---|---|---|---|---|---|
| Lateral occipital cortex, sup. div. | R | | | 38 | -88 | 18 | Area hOc4lp (52.1) | 4.00 |
| Lateral occipital cortex, inf. div. | R | | | 38 | -66 | -6 | Area FG1 (8.2) | 3.98 |
| Occipital pole | L | 391 | 0.005 | -16 | -100 | 10 | Area hOc2 (V2, 45.2) | 4.95 |
| Occipital fusiform gyrus | L | | | -18 | -88 | -8 | Area hOc3v (V3v, 69.8) | 4.43 |
| Occipital pole | L | | | -24 | -94 | 24 | Area hOc4d (V3A, 43.9) | 3.36 |
| ***YA: Dual$_{RRI}$ + > Single*** | | | | | | | | |
| Precentral gyrus | R | 24983 | < 0.001 | 32 | -10 | 60 | Area 6d1 (20.5) | > 8 |
| Postcentral gyrus | R | | | 44 | -36 | 56 | Area 1 (39.4) | > 8 |
| Precentral gyrus | R | | | 40 | -16 | 56 | Area 4a (17.9) | > 8 |
| Postcentral gyrus | R | | | 42 | -26 | 42 | Area 2 (50.9) | > 8 |
| Supplementary motor area | L/R | | | 4 | -6 | 52 | Area 6mc / SMA (46.3) | > 8 |
| Superior parietal lobule | L | | | -38 | -44 | 60 | Area 7PC (SPL, 41.0) | > 8 |
| Precentral gyrus | L | | | -38 | -18 | 60 | Area 6d1 (15.6) | > 8 |
| *This cluster is too large for the peaks to represent all covered macrostructural areas, but it also includes right basal ganglia and insular cortex.* | | | | | | | | |
| Cerebellum VI | L | 4023 | < 0.001 | -20 | -54 | -22 | – | > 8 |
| Cerebellum V | L/R | | | -4 | -62 | -16 | – | > 8 |
| Cerebellum VI | R | | | 20 | 52 | -22 | – | > 8 |
| Cerebellum I-IV | L/R | | | 0 | -50 | -4 | – | > 8 |
| Thalamus | L | 1160 | < 0.001 | -14 | -18 | 6 | – | > 8 |
| Pallidum | L | | | -16 | -8 | 0 | – | 5.97 |
| Middle frontal gyrus | L | 664 | < 0.001 | -40 | 32 | 30 | – | 4.90 |
| Frontal pole | L | | | -32 | 36 | 22 | – | 4.60 |
| Frontal pole | R | 547 | 0.001 | 36 | 42 | 28 | – | 5.61 |
| Precentral gyrus | L | 544 | 0.001 | -54 | 6 | 34 | Area 44 (31.8) | 5.66 |
| Insular cortex | L | 489 | 0.002 | -32 | 22 | -4 | Area Id7 (32.8) | 5.63 |
| ***OA: Dual$_{RRI}$ + > Single*** | | | | | | | | |
| Precentral gyrus | R | 47967 | < 0.001 | 34 | -24 | 56 | Area 4p (57.7) | > 8 |
| | R | | | 36 | -20 | 64 | Area 4a (10.6) | > 8 |
| | R | | | 42 | -16 | 54 | Area 3b (19.8) | > 8 |
| Cerebellum V | L | | | -16 | -52 | -20 | – | > 8 |
| Precentral gyrus | R | | | 28 | -8 | 64 | Area 6d1 (30.9) | > 8 |
| | L | | | -32 | -26 | 50 | Area 4p (41.9) | > 8 |
| Cerebellum V | R | | | 16 | -52 | -18 | – | > 8 |
| Precentral gyrus | L | | | -32 | -24 | 64 | Area 4a (49.1) | > 8 |

*This cluster is too large for the peaks to represent all covered macrostructural areas, but it also includes bilateral SPL, SMA, preSMA, cingulate gyrus, ant. div., frontal pole, insular cortex, and basal ganglia.*

| | | | | | | | | |
|---|---|---|---|---|---|---|---|---|
| ***YA: $Dual_{RRI}$ – < Single*** | | | | | | | | |
| Frontal medial cortex | L | 892 | < 0.001 | -8 | 48 | -18 | Area Fp2 (21.6) | 5.61 |
| Frontal pole | L/R | | | 0 | 60 | -8 | – | 5.43 |
| | L | | | -10 | 56 | 6 | Area p32 (58.3) | 4.70 |
| Frontal medial cortex | R | | | 6 | 52 | -16 | Area Fp2 (25.2) | 4.59 |
| Paracingulate gyrus | L | | | -8 | 36 | -12 | Area s32 (70.8) | 4.21 |
| Subcallosal cortex | R | | | 2 | 30 | -10 | Area s24 (62.2) | 3.73 |
| Frontal medial cortex | R | | | 4 | 38 | -18 | Area s32 (56.2) | 3.61 |
| Inferior parietal lobule | L | 250 | 0.034 | -52 | -72 | 34 | Area PGp (IPL, 52.4) | 5.73 |
| Angular gyrus | L | | | -62 | -60 | 22 | Area PGa (IPL, 42.4) | 3.64 |
| Inferior parietal lobule | L | | | -52 | -60 | 48 | Area PFm (IPL, 52.1) | 3.38 |
| ***OA: $Dual_{RRI}$ – < Single*** | | | | | | | | |
| *nothing significant* | | | | | | | | |
| ***YA: $Dual_{RRI}$ + > $Dual_{RRC}$*** | | | | | | | | |
| Precentral gyrus | R | 7540 | < 0.001 | 26 | -8 | 58 | Area 6d3 (42.2) | > 8 |
| Superior frontal gyrus | L | | | -22 | -6 | 60 | Area 6d1 (29.2) | 7.69 |
| Paracingulate gyrus | L/R | | | 2 | 18 | 40 | – | 6.12 |
| Precentral gyrus | L | | | -48 | 0 | 34 | Area 44 (2.6) | 5.57 |
| Inferior frontal gyrus, pars op. | R | | | 50 | 14 | 26 | Area 44 (36.0) | 5.48 |
| Cingulate gyrus, ant. div. | L/R | | | 6 | 4 | 44 | Area 6mr / preSMA (16.6) | 5.33 |
| Superior parietal lobule | R | 5261 | < 0.001 | 22 | -58 | 62 | Area 7A (SPL, 53.4) | > 8 |
| | R | | | 36 | -44 | 52 | Area hIP3 (IPS, 63.0) | > 8 |
| Supramarginal gyrus, ant. div. | R | | | 54 | -22 | 40 | Area PFt (IPL, 47.2) | 5.99 |
| | R | | | 60 | -34 | 32 | Area PF (IPL, 46.8) | 5.30 |
| Angular gyrus | R | | | 36 | -56 | 38 | Area hIP1 (IPS, 40.3) | 4.55 |
| Postcentral gyrus | R | | | 60 | -12 | 32 | Area 1 (38.4) | 4.01 |
| Superior parietal lobule | L | 4459 | < 0.001 | -22 | -60 | 60 | Area 7A (SPL, 48.3) | > 8 |
| | L | | | -42 | -44 | 52 | Area 7PC (SPL, 21.4) | > 8 |
| Supramarginal gyrus, ant. div. | L | | | -38 | -38 | 38 | Area hIP1 (IPS, 43.4) | > 8 |
| Intraparietal sulcus | L | | | -30 | -52 | 48 | Area hIP3 (IPS, 57.4) | 6.31 |
| Supramarginal gyrus, ant. div. | L | | | -58 | -30 | 34 | Area PFt (IPL, 65.1) | 4.93 |
| Cerebellum VI | L | 1647 | < 0.001 | -20 | -72 | -24 | – | 5.85 |
| Cerebellum Crus I | L | | | -48 | -58 | -34 | – | 4.56 |

| | | | | | | | | |
|---|---|---|---|---|---|---|---|---|
| Cerebellum Vermis VI | L/R | | | 0 | -76 | -16 | – | 4.54 |
| | R | | | 2 | -66 | -28 | – | 4.11 |
| Insular cortex | R | 901 | < 0.001 | 34 | 22 | -6 | Area Id7 (13.9) | 6.45 |
| Inferior frontal gyrus, pars op. | R | | | 60 | 14 | 2 | – | 3.52 |
| Thalamus | R | 781 | < 0.001 | 18 | -22 | 8 | – | 5.68 |
| Pallidum | R | | | 18 | -8 | -4 | – | 4.10 |
| Frontal pole | R | 532 | 0.001 | 40 | 38 | 32 | – | 5.21 |
| Middle frontal gyrus | R | | | 34 | 30 | 24 | – | 4.09 |
| Insular cortex | L | 431 | 0.003 | -32 | 22 | -2 | Area Id7 (67.0) | 4.67 |
| Middle frontal gyrus | L | 415 | 0.004 | -42 | 30 | 34 | – | 5.23 |
| Frontal pole | L | | | -38 | 48 | 12 | – | 3.66 |
| Thalamus | L | 253 | 0.032 | -12 | -20 | 8 | – | 4.14 |
| ***OA: $Dual_{RRI}$ + > $Dual_{RRC}$*** | | | | | | | | |
| Superior frontal gyrus | R | 23654 | < 0.001 | 22 | -4 | 54 | Area 6d3 (84.9) | > 8 |
| | L | | | -20 | 0 | 50 | Area 6d3 (69.2) | 7.56 |
| Middle frontal gyrus | R | | | 32 | 4 | 60 | – | 7.25 |
| Pallidum | R | | | 16 | -2 | 2 | – | 7.23 |
| Insular cortex | L | | | -32 | 22 | -6 | Area Id7 (3.8) | 6.92 |
| Superior frontal gyrus | R | | | 24 | -2 | 66 | Area 6d2 (34.6) | 6.77 |
| Thalamus | L | | | -18 | -22 | 10 | – | 6.40 |
| Cerebellum VI | L | | | -10 | -72 | -26 | – | 6.31 |
| Thalamus | R | | | 12 | -18 | 8 | – | 6.24 |
| Superior parietal lobule | R | 4022 | < 0.001 | 20 | -66 | 58 | Area 7A (SPL, 75.9) | > 8 |
| | R | | | 34 | -50 | 44 | Area hIP3 (IPS, 53.3) | 7.76 |
| Angular gyrus | R | | | 38 | -54 | 50 | Area hIP3 (SPL, 12.7) | 7.36 |
| Intraparietal sulcus | R | | | 32 | -68 | 44 | Area hIP6 (IPS, 49.8) | 5.04 |
| Postcentral gyrus | R | | | 26 | -42 | 60 | Area 2 (51.5) | 4.21 |
| | R | | | 42 | -22 | 34 | Area 3b (15.6) | 3.88 |
| Superior parietal lobule | L | 3554 | < 0.001 | -18 | -64 | 58 | Area 7A (SPL, 61.8) | > 8 |
| | L | | | -34 | -44 | 44 | Area hIP1 (IPS, 20.0) | 6.57 |
| Intraparietal sulcus | L | | | -28 | -62 | 44 | Area hIP3 (IPS, 54.4) | 6.55 |
| ***YA: $Dual_{RRI}$ – < $Dual_{RRC}$*** | | | | | | | | |
| *nothing significant* | | | | | | | | |
| ***OA: $Dual_{RRI}$ – < $Dual_{RRC}$*** | | | | | | | | |

*nothing significant*

---

**Abbreviations.** Ant. div.: Anterior division, Cytoarchit. assignm.: Cytoarchitectonic assignment, FG: Fusiform gyrus, FP: Frontal pole, inf. div.: Inferior division, IPL: Inferior parietal lobule, IPS: Intraparietal sulcus, OA, Old adults, Oc: Occipital cortex, OP: Opercular cortex, PFm: Parietal area F, part m, PFop: Parietal area, opercular subregion, PFt: Parietal area F, part t, PGa: Parietal area G, anterior, PGp: Parietal area G, posterior, post. div.: Posterior division, preSMA: Presupplementary motor area, RRC: Response–response congruent, RRI: Response–response incongruent, SMA: Supplementary motor area, SPL: Superior parietal lobule, sup. div.: Superior division, VS: Ventral somatosensory, YA: Young adults.

**Note.** Selection of peaks based on macroanatomical and cytoarchitectonic representative regions.

The contrasts were estimated via a conjunction combining the contrast and the regressor's main effect, marked with + or – for each group separately. All activations are reported at a cluster-forming threshold with voxel-level uncorrected at $p < 0.001$, and cluster-level family-wise error (cFWE) corrected at $p < 0.05$.

**Table S7.**

Brain regions showing significant interactions between each task-specific contrast and age.

| ***Contrast /*** **Macroanatomical structure** | **H** | **Cluster extent ($k_E$)** | **Cluster *p(cFWE)*** | **MNI coordinates x** | **y** | **z** | **Cytoarchit. assignm. (Overlap in %)** | **Peak *Z* value** |
|---|---|---|---|---|---|---|---|---|
| ***(Dual$_{RRC}$ vs. Single) × Age 1 ∩ OA: Dual$_{RRC}$ > Single*** | | | | | | | | |
| *not significant* | | | | | | | | |
| ***(Dual$_{RRC}$ vs. Single) × Age 2 ∩ YA: Dual$_{RRC}$ > Single*** | | | | | | | | |
| *not significant* | | | | | | | | |
| ***(Dual$_{RRI}$ vs. Single) × Age 1 ∩ OA: Dual$_{RRI}$ > Single*** | | | | | | | | |
| Precentral gyrus | R | 878 | < 0.001 | 6 | -26 | 68 | Area 4a (45.6) | 5.56 |
| | L | | | -2 | -24 | 66 | – | 5.39 |
| Postcentral gyrus | L | | | -16 | -36 | 66 | Area 3b (32.2) | 3.95 |
| Frontal pole | L | 690 | < 0.001 | -18 | 54 | -2 | – | 5.00 |
| Cingulate gyrus, ant. div. | L | | | -6 | 36 | 0 | Area p24ab (72.0) | 4.41 |
| Frontal pole | L | | | -30 | 38 | 0 | – | 4.07 |
| Paracingulate gyrus | L | | | -12 | 42 | -6 | Area p32 (43.5) | 3.82 |
| Frontal pole | L | | | -26 | 48 | 22 | – | 3.73 |
| FP / Middle frontal gyrus | L | 500 | 0.001 | -24 | 24 | 28 | – | 5.07 |
| FP / Superior frontal gyrus | L | | | -24 | 40 | 36 | – | 3.99 |
| Frontal pole | R | 394 | 0.005 | 18 | 52 | 2 | – | 4.28 |
| | R | | | 32 | 34 | 10 | – | 4.21 |
| | R | | | 20 | 54 | 14 | Area Fp1 (3.1) | 4.00 |
| Cingulate gyrus, ant. div. | R | | | 10 | 38 | 6 | Area p24ab (6.9) | 3.55 |
| Paracingulate gyrus | R | | | 16 | 44 | -2 | Area p32 (3.4) | 3.49 |
| FP / Middle frontal gyrus | R | | | 44 | 40 | 14 | Area 45 (9.3) | 3.35 |
| Cingulate gyrus, ant. div. | R | | | 8 | 38 | -4 | Area p24c (28.5) | 3.34 |
| Pallidum | L | 276 | 0.023 | -18 | -2 | 14 | – | 3.92 |
| Putamen | L | | | -22 | 22 | -2 | – | 3.71 |
| | L | | | -14 | 8 | -4 | – | 3.70 |
| Insular cortex | L | | | -28 | 30 | 8 | Area Id7 (3.3) | 3.47 |
| Inferior frontal gyrus, pars triang. | L | | | -40 | 34 | 8 | Area 45 (1.4) | 3.32 |
| | L | | | -38 | 30 | 10 | Area OP9 (4.9) | 3.30 |
| ***(Dual$_{RRI}$ vs. Single) × Age 2 ∩ YA: Dual$_{RRI}$ > Single*** | | | | | | | | |
| Postcentral gyrus | R | 657 | < 0.001 | 42 | -36 | 52 | Area 2 (51.1) | 5.59 |
| Superior parietal lobule | R | | | 24 | -42 | 72 | – | 5.51 |
| | R | | | 28 | -42 | 70 | Area 1 (38.3) | 5.51 |

| | | | | | | | | |
|---|---|---|---|---|---|---|---|---|
| | R | | | 20 | -60 | 66 | Area 7A (SPL, 80.4) | 4.06 |
| Supramarginal gyrus, ant. div. | R | | | 36 | -32 | 34 | – | 3.58 |
| Precentral gyrus | R | 348 | 0.009 | 28 | -12 | 58 | Area 6d1 (39.2) | 4.91 |
| | R | | | 38 | -10 | 60 | Area 4a (5.4) | 4.67 |
| Superior frontal gyrus | R | | | 16 | -6 | 74 | Area 6d1 (33.0) | 3.45 |
| ***(Dual$_{RRI}$ vs. Dual$_{RRC}$) × Age 1 ∩ OA: Dual$_{RRI}$ > Dual$_{RRC}$*** | | | | | | | | |
| Superior frontal gyrus | L | 299 | 0.017 | -16 | 28 | 34 | – | 4.35 |
| | L | | | -24 | 24 | 30 | – | 3.91 |
| Frontal pole | L | | | -22 | 38 | 38 | – | 3.61 |
| ***(Dual$_{RRI}$ vs. Dual$_{RRC}$) × Age 2 ∩ YA: Dual$_{RRI}$ > Dual$_{RRC}$*** | | | | | | | | |
| *not significant* | | | | | | | | |

**Abbreviations.** Ant. div.: Anterior division, Cytoarchit. assignm.: Cytoarchitectonic assignment, FP: Frontal pole, OA: Old adults, OP: Opercular cortex, pars triang.: pars triangularis, RRC: Response–response congruent, RRI: Response–response incongruent, YA: Young adults.

**Note.** Selection of peaks based on macroanatomical and cytoarchitectonic representative regions.

The interactions were estimated via a conjunction combining the interaction and the age-specific task contrast. All activations are reported at a cluster-forming threshold with voxel-level uncorrected at $p < 0.001$, and cluster-level family-wise error (cFWE) corrected at $p < 0.05$.

*Response grouping*

Besides exploring whether participants engaged an alternative mechanism of response grouping during dual-task response-code conflict that could explain the asymmetric dual-task costs under R-R incongruency (see section: Supplementary Material – Behavioral data – Response grouping), we additionally tested whether there were any brain regions associated with non-synchronous responding during dual-task crosstalk. As mentioned before, non-synchronous responses were defined via the IRI (≥ 48 ms).

For this purpose, we implemented the same single-subject level model for assessing the task-related brain effects (see section: Methods – Data analysis – Task-related brain activity). The only difference was that we entered a second parametric modulator (besides hand(s) used for responding) for response grouping based on IRI (non-grouped trials = -1, single trials = 0, or grouped trials = 1). For each participant, we created difference contrasts assessing dual-task response-code conflict [$Dual_{RRI}$ > ($Dual_{SRC-RRC}$ & $Dual_{SRI-RRC}$)] using the task regressors and parametric regressors separately. We entered these two regressors ($Dual_{RRI>RRC}$ and $Dual_{RRI>RRC}$ × IRI) into a group-level between-subject model for each age group, assuming unequal variance between subjects. This resulted in a design matrix with four regressors at the group level.

Our results showed that the positive effect of $Dual_{RRI>RRC}$ × IRI (vs. baseline) across both age groups revealed increased brain activity in bilateral occipital pole, TPS, angular gyrus, and posterior cingular cortex, as well as right dPMC and middle and superior temporal gyrus (see **Figure S6** and **Table S8**). In other words, under response-code conflict, these areas showed a parametric increase of activation with increasing response grouping, on average, across both age groups. We did not find any significant negative effects or age differences. One consideration to keep in mind is that participants executed non-grouped responses only in a small portion of trials (< 10%). Thus, the imbalanced number of trials might have a tentative impact on the power of this supplementary analysis.

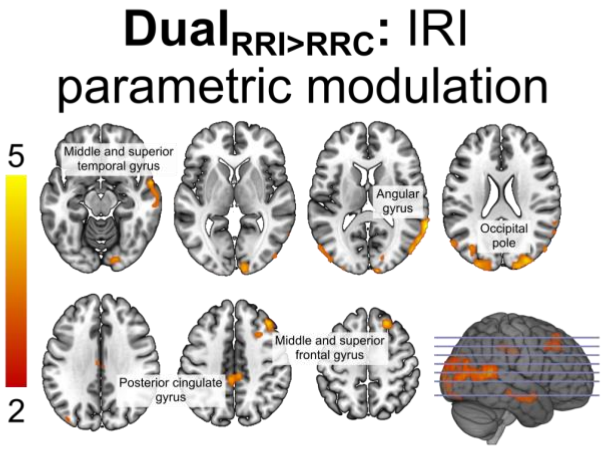


**Figure S6.**

Brain areas showing parametric modulation of inter-response interval under response-code conflict across both age groups (estimated using the $Dual_{RRI}$ > $Dual_{RRC}$ contrast from the first-level analysis). All effects were significant at cluster-level $p$ < 0.05 (cFWE-corrected) with a cluster-forming threshold of $p$ < 0.001 at voxel level. **Abbreviations.** IRI: Inter-response interval, RRC: Response–response congruent, RRI: Response–response incongruent.

**Table S8.**

Brain regions showing significant inter-response interval parametric modulation during response-code conflict (Dual$_{RRI}$ > Dual$_{RRC}$ at single-subject level).

| *Contrast /* Macroanatomical structure | H | Cluster extent ($k_E$) | Cluster *p(cFWE)* | MNI coordinates x | y | z | Cytoarchit. assignm. (Overlap in %) | Peak *Z* value |
|---|---|---|---|---|---|---|---|---|
| ***Dual$_{RRI>RRC}$ × IRI +*** | | | | | | | | |
| Occpital pole | R | 653 | < 0.001 | 10 | -100 | 14 | Area hOc2 (V2, 62.6) | 4.54 |
| Lateral occipital cortex, sup. div. | R | | | 30 | -88 | 24 | Area hOc4d (V3A, 44.9) | 4.52 |
| Occipital pole | R | | | 20 | -94 | 20 | Area hOc3d (V3d, 43.1) | 4.27 |
| | R | | | 12 | -100 | 2 | Area hOc1 (V1, 83.1) | 3.97 |
| Occipital fusiform gyrus | R | | | 18 | -88 | -14 | Area hOc3v (V3v, 59.8) | 3.91 |
| Occpital pole | L | 648 | < 0.001 | -26 | -94 | 22 | Area hOc4d (V3A, 38.9) | 4.02 |
| Lateral occipital cortex, sup. div. | L | | | -54 | -74 | 16 | Area PGp (IPL, 40.2) | 3.94 |
| Intraparietal sulcus | L | | | -34 | -74 | 24 | Area hIP5 (IPS, 47.9) | 3.88 |
| Occpital pole | L | | | -12 | -96 | 24 | Area hOc3d (V3d, 64.7) | 3.58 |
| Lateral occipital cortex, sup. div. | L | | | -38 | -86 | 16 | Area hOc4lp (56.7) | 3.49 |
| Angular gyrus | R | 594 | 0.001 | 68 | -48 | 12 | – | 4.85 |
| Lateral occipital cortex, inf. div. | R | | | 48 | -82 | 6 | Area hOc4la (73.7) | 4.61 |
| Middle temporal gyrus, TO part | R | | | 64 | -56 | 8 | Area PGa (IPL, 12.2) | 4.38 |
| Lateral occipital cortex, inf. div. | R | | | 60 | -62 | 10 | Area hOc4la (7.5) | 4.20 |
| Inferior parietal lobule | R | | | 52 | -76 | 24 | Area PGp (IPL, 61.8) | 3.82 |
| Middle frontal gyrus | R | 292 | 0.021 | 40 | 30 | 42 | – | 4.38 |
| Superior frontal gyrus | R | | | 22 | 30 | 52 | – | 4.35 |
| Middle frontal gyrus | R | | | 28 | 16 | 42 | Area 6d3 (2.4) | 3.49 |
| Cingulate gyrus, post. div. | R | 283 | 0.023 | 2 | -22 | 36 | – | 4.10 |
| | L | | | -4 | -34 | 40 | – | 4.03 |
| | R | | | 6 | -32 | 42 | Area 5Ci (SPL, 46.1) | 3.80 |
| Superior temporal gyrus, ant. div. | R | 245 | 0.039 | 58 | 2 | -14 | Area TE 3 (49.6) | 4.25 |
| Middle temporal gyrus, ant. div. | R | | | 60 | -4 | -18 | Area TE 3 (4.2) | 4.24 |
| Middle temporal gyrus, post. div. | R | | | 64 | -24 | -12 | – | 3.70 |
| ***Dual$_{RRI>RRC}$ × IRI –*** | | | | | | | | |
| *not significant* | | | | | | | | |

| ***$Dual_{RRI>RRC}$ × IRI: OA > YA*** |
| --- |
| *not significant* |
| ***$Dual_{RRI>RRC}$ × IRI: YA > OA*** |
| *not significant* |

**Abbreviations.** Ant. div.: Anterior division, Cytoarchit. assignm.: Cytoarchitectonic assignment, inf. div.: Inferior division, IPL: Inferior parietal lobule, IPS: Intraparietal sulcus, IRI: Inter-response interval, OA: Old adults, Oc: Occipital cortex, PGa: Parietal area G, anterior, PGp: Parietal area G, posterior, post. div.: posterior division, RRC: Response–response congruent, RRI: Response–response incongruent, SPL: Superior parietal lobule, sup. div.: Superior division, TO: Temporooccipital, YA: Young adults.

**Note.** Selection of peaks based on macroanatomical and cytoarchitectonic representative regions.

The model included two regressors: The task regressor ($Dual_{RRI}$ > $Dual_{RRC}$) estimated at the single-subject level and its parametric modulation (PM) through IRI. The contrasts estimated the PM-regressor main effect (+ or –) or its age contrast. All activations are reported at a cluster-forming threshold with voxel-level uncorrected at $p < 0.001$, and cluster-level family-wise error (cFWE) corrected at $p < 0.05$.

**Table S9.**

Brain areas from the task-specific response-code conflict network ($Dual_{RRI}$ > $Dual_{RRC}$) showing a significant negative correlation with dual-task performance, as represented by the BIS in S-R compatible and R-R incongruent condition, crossmodal attention, working memory, and global task-switching costs during the R-R incongruent (vs. baseline) condition.

| *Contrast /* **Macroanatomical structure** | H | Cluster extent ($k_E$) | Cluster *p(cFWE)* | MNI coordinates x | y | z | Cytoarchit. assignm. (Overlap in %) | Peak *Z* value |
|---|---|---|---|---|---|---|---|---|
| ***Dual-task performance*** | | | | | | | | |
| ***$Dual_{RRI}$ × $BIS_{SRC-RRI}$ – (Mask: $Dual_{RRI}$ > $Dual_{RRC}$)*** | | | | | | | | |
| Cuneal cortex | R | 2762 | < 0.001 | 2 | -78 | 22 | – | 4.79 |
| Lingual gyrus | L | | | -18 | -72 | -4 | Area hOc3v (V3v, 53.6) | 4.74 |
| Cuneal cortex | L | | | -4 | -82 | 26 | Area hOc3d (V3d, 45.4) | 4.60 |
| Lingual gyrus | L | | | -2 | -68 | 6 | Area hOc1 (V1, 40.5) | 4.39 |
| Cuneal cortex | R | | | 26 | -68 | 22 | – | 4.29 |
| Intracalcarine cortex | L | | | -4 | -80 | 4 | Area hOc1 (V1, 75.8) | 4.27 |
| | R | | | 16 | -76 | 16 | Area hOc3d (V3d, 8.1) | 4.19 |
| Lateral occipital cortex, sup. div. | R | | | 18 | -88 | 30 | Area hOc4d (V3A, 59.5) | 4.02 |
| Lingual gyrus | R | | | 24 | -64 | 0 | Area hOc1 (V1, 69.1) | 3.92 |
| ***Crossmodal attention*** | | | | | | | | |
| ***$Dual_{RRI}$ × Att – (Mask: $Dual_{RRI}$ > $Dual_{RRC}$)*** | | | | | | | | |
| Brainstem | R | 903 | < 0.001 | 16 | -20 | -8 | – | 5.75 |
| | L | | | -8 | -10 | -8 | – | 4.70 |
| | R | | | 8 | -28 | -16 | – | 4.53 |
| | R | | | 8 | -8 | -10 | – | 4.43 |
| Thalamus | R | | | 18 | -14 | 2 | – | 3.99 |
| Pallidum | R | | | 18 | -4 | 6 | – | 3.67 |
| Thalamus | R | | | 22 | -20 | 8 | – | 3.46 |
| ***Working memory*** | | | | | | | | |
| ***$Dual_{RRI}$ × WM – (Mask: $Dual_{RRI}$ > $Dual_{RRC}$)*** | | | | | | | | |
| *not significant* | | | | | | | | |
| ***Task switching*** | | | | | | | | |
| ***$Dual_{RRI}$ × gTSC – (Mask: $Dual_{RRI}$ > $Dual_{RRC}$)*** | | | | | | | | |

| | | | | | | | | |
|---|---|---|---|---|---|---|---|---|
| Intraparietal sulcus | R | 488 | 0.002 | 30 | -68 | 44 | Area hIP6 (IPS, 43.1) | 4.90 |
| Superior parietal lobule | R | | | 22 | -72 | 62 | Area 7P (SPL, 28.6) | 4.43 |
| | R | | | 18 | -66 | 48 | Area 7A (SPL, 19.4) | 4.23 |
| Intraparietal sulcus | L | 279 | 0.023 | -32 | -38 | 38 | Area hIP1 (IPS, 4.7) | 4.56 |
| Superior parietal lobule | L | | | -46 | -42 | 54 | Area 2 (16.6) | 3.72 |
| Cerebellum VI | R | 257 | 0.031 | 22 | -74 | -20 | – | 4.35 |
| Cerebellum Crus I | R | | | 40 | -66 | -24 | – | 4.35 |
| Inferior temporal gyrus, TO part | R | | | 48 | -56 | -20 | Area FG4 (50.8) | 3.42 |

**Abbreviations.** BIS: Balanced integration score, Cytoarchit. assignm.: Cytoarchitectonic assignment, gTSC: Global task-switching costs, IPS: Intraparietal sulcus, Oc: Occipital cortex, RRC: Response–response congruent, RRI: Response–response incongruent, Att: Crossmodal attention, SPL: Superior parietal lobule, SRC: Stimulus–response compatible, sup. div.: Superior division, TO: Temporooccipital, WM: Working memory.

**Note.** Selection of peaks based on macroanatomical and cytoarchitectonic representative regions.

The contrasts for each separate covariance model included the negative main effect (–) of the covariance regressor for $Dual_{RRI}$. The contrasts were masked with the response-code conflict network ($Dual_{RRI}$ > $Dual_{RRC}$ contrast at $p < 0.001$). All activations are reported at a cluster-forming threshold with voxel-level uncorrected at $p < 0.001$, and cluster-level family-wise error (cFWE) corrected at $p < 0.05$.

**Table S10.**

Brain areas from the task-specific response-code conflict network ($Dual_{RRI}$ > $Dual_{RRC}$) showing significant age difference (OA > YA) in dual-task performance, as represented by the BIS in S-R compatible and R-R incongruent condition, crossmodal attention, working memory, and global task-switching costs during the R-R incongruent (vs. baseline) condition.

| ***Contrast /*** **Macroanatomical structure** | **H** | **Cluster extent ($k_E$)** | **Cluster *p(cFWE)*** | **MNI coordinates** | | | **Cytoarchit. assignm. (Overlap in %)** | **Peak *Z* value** |
|---|---|---|---|---|---|---|---|---|
| | | | | **x** | **y** | **z** | | |
| ***Dual-task performance*** | | | | | | | | |
| ***$Dual_{RRI}$ × $BIS_{SRC-RRI}$ : OA > YA (Mask: $Dual_{RRI}$ > $Dual_{RRC}$)*** | | | | | | | | |
| *not significant* | | | | | | | | |
| ***Crossmodal attention*** | | | | | | | | |
| ***$Dual_{RRI}$ × Att : OA > YA (Mask: $Dual_{RRI}$ > $Dual_{RRC}$)*** | | | | | | | | |
| Occipital fusiform gyrus | R | 380 | 0.006 | 20 | -76 | -18 | Area hOc3v (V3v, 44.3) | 5.34 |
| Lingual gyrus | R | | | 10 | -82 | -14 | Area hOc2 (V2, 51.4) | 4.69 |
| Occipital fusiform gyrus | R | | | 34 | -72 | -18 | Area FG1 (38.9) | 4.40 |
| Occipital pole | R | | | 14 | -96 | -10 | Area hOc2 (V2, 45.0) | 3.61 |
| Intraparietal sulcus | L | 244 | 0.035 | -32 | -64 | 46 | Area hIP6 (IPS, 35.9) | 3.72 |
| | L | | | -30 | -58 | 36 | Area hIP1 (IPS, 48.6) | 3.64 |
| Superior parietal lobule | L | | | -28 | -64 | 56 | Area 7A (SPL, 49.4) | 3.32 |
| ***Working memory*** | | | | | | | | |
| ***$Dual_{RRI}$ × WM : OA > YA (Mask: $Dual_{RRI}$ > $Dual_{RRC}$)*** | | | | | | | | |
| *not significant* | | | | | | | | |
| ***Task switching*** | | | | | | | | |
| ***$Dual_{RRI}$ × gTSC : OA > YA (Mask: $Dual_{RRI}$ > $Dual_{RRC}$)*** | | | | | | | | |
| *not significant* | | | | | | | | |

**Abbreviations.** BIS: Balanced integration score, Cytoarchit. assignm.: Cytoarchitectonic assignment, FG: Fusiform gyrus, gTSC: Global task-switching costs, IPS: Intraparietal sulcus, OA: Old adults, Oc: Occipital cortex, RRC: Response–response congruent, RRI: Response–response incongruent, Att: Crossmodal attention, SPL: Superior parietal lobule, SRC: Stimulus–response compatible, WM: Working memory, YA: Young adults.

**Note.** Selection of peaks based on macroanatomical and cytoarchitectonic representative regions.

The contrasts for each separate covariance model were estimated by contrasting the age groups (OA > YA) for the covariance regressor for $Dual_{RRI}$. The contrasts were masked with the response-code conflict network ($Dual_{RRI}$ > $Dual_{RRC}$ contrast at $p < 0.001$). All activations are reported at a cluster-forming threshold with voxel-level uncorrected at $p < 0.001$, and cluster-level family-wise error (cFWE) corrected at $p < 0.05$.

**Table S11.**

Common brain areas in which the connectivity with the right dPMC and SPL was significantly modulated as a function of $Dual_{RRI}$ and $Dual_{RRC}$, assessed through a minimal conjunction of both experimental conditions.

| Seed | *Contrast /* Macroanatomical structure | H | Cluster extent ($k_E$) | MNI coordinates x | y | z | Ciytoarchit. assignm. (Overlap in %) | Peak *T* value |
|---|---|---|---|---|---|---|---|---|
| **Right dPMC** | ***min((YA: $Dual_{RRC}$ + ∩ OA: $Dual_{RRC}$ +), (YA: $Dual_{RRI}$ + ∩ OA: $Dual_{RRI}$ +))*** | | | | | | ***(Mask: $Dual_{RRI}$ + > $Dual_{RRC}$)*** | |
| | Dorsal premotor cortex | R | 2202 | 24 | -6 | 54 | Area 6d3 (70.5) | 10.50 |
| | Middle frontal gyrus | R | | 36 | -2 | 56 | – | 6.01 |
| | Inferior parietal cortex | R | | 50 | -32 | 26 | Area PF (IPL, 43.2) | 4.64 |
| | Precentral gyrus | R | | 26 | -28 | 54 | Area 4p (35.0) | 4.40 |
| | Intraparietal sulcus | R | | 42 | -48 | 52 | Area hIP3 (IPS, 49.8) | 4.39 |
| | Postcentral gyrus | R | | 36 | -30 | 56 | Area 3b (57.2) | 4.34 |
| | Superior parietal lobule | R | | 34 | -50 | 58 | Area 7PC (SPL, 32.3) | 4.31 |
| | Postcentral gyrus | L | 1125 | -48 | -34 | 46 | Area 2 (41.8) | 4.97 |
| | Intraparietal sulcus | L | | -32 | -56 | 48 | Area hIP3 (IPS, 78.7) | 4.85 |
| | Superior parietal lobule | L | | -16 | -70 | 54 | Area 7A (SPL, 45.4) | 4.06 |
| | Supramarginal gyrus, ant. div. | L | | -58 | -32 | 38 | Area PFt (IPL, 61.0) | 3.83 |
| | Precuneus cortex | L | | -10 | -64 | 56 | Area 7A (SPL, 54.1) | 3.80 |
| | Superior parietal lobule | L | | -24 | -42 | 56 | Area 5L (SPL, 42.4) | 3.63 |
| | Precentral gyrus | L | 1114 | -30 | -10 | 54 | Area 6d1 (19.4) | 4.73 |
| | Inferior frontal gyrus, pars op. | L | | -52 | 12 | 22 | Area 44 (62.2) | 4.27 |
| | Middle frontal gyrus | L | | -50 | 8 | 36 | Area 44 (20.8) | 4.12 |
| | Superior frontal gyrus | L | | -22 | 2 | 54 | Area 6d3 (84.4) | 3.90 |
| | Precentral gyrus | L | | -40 | 2 | 28 | Area 44 (14.7) | 3.71 |
| | Supplementary motor area | L/R | 506 | 4 | -4 | 52 | Area 6mc / SMA (24.8) | 4.72 |
| | Precentral gyrus | R | | 8 | -14 | 48 | – | 4.54 |
| | Paracingulate gyrus | R | | 2 | 6 | 4 | – | 4.02 |
| | Cingulate gyrus, ant. div. | R | | 4 | 10 | 32 | Area 33 (48.2) | 3.91 |
| | Paracingulate gyrus | R | | 6 | 16 | 38 | – | 3.82 |
| | Superior frontal gyrus | R | | 2 | 20 | 52 | – | 3.35 |
| | Inferior frontal gyrus, pars op. | R | 443 | 46 | 8 | 20 | Area 44 (50.7) | 4.15 |
| | Insular cortex | R | | 36 | 18 | 0 | Area Id7 (32.8) | 4.06 |
| | Opercular cortex | R | | 38 | 24 | 8 | Area OP9 (46.4) | 3.72 |
| | Insular cortex | L | 306 | -28 | 16 | -8 | – | 4.38 |
| | Frontal orbital cortex | L | | -30 | 22 | -8 | – | 4.30 |

| | | | | | | | | |
|---|---|---|---|---|---|---|---|---|
| | Putamen | L | | -28 | 12 | 6 | – | 3.99 |
| | Middle frontal gyrus | L | 265 | -40 | 28 | 34 | – | 4.47 |
| | Frontal pole | L | | -34 | 40 | 28 | – | 3.60 |
| | Precentral gyrus | L | 1 | -20 | -12 | 70 | Area 6d1 (60.9) | 3.28 |
| **Right SPL** | ***min((YA: Dual$_{RRC}$ + ∩ OA: Dual$_{RRC}$ +), (YA: Dual$_{RRI}$ + ∩ OA: Dual$_{RRI}$ +))*** | | | | | | ***(Mask: Dual$_{RRI}$ + > Dual$_{RRC}$)*** | |
| | Cerebellum VI | L | 1724 | -32 | -56 | -32 | – | 5.48 |
| | Cerebellum VI | R | | 22 | -62 | -28 | – | 4.77 |
| | Cerebellum V | L/R | | 0 | -58 | -24 | – | 4.54 |
| | Cerebellum I-IV | L/R | | 2 | -56 | -22 | – | 4.46 |
| | Cerebellum VI | R | | 8 | -62 | -28 | Ventral Dentate Ncl. (6.2) | 4.43 |
| | Cerebellum V | L | | -24 | -40 | -30 | – | 4.20 |
| | Superior parietal lobule | R | 1055 | 18 | -60 | 62 | Area 7A (SPL, 70.3) | 7.87 |
| | Intraparietal sulcus | R | | 34 | -70 | 48 | Area hIP6 (IPS; 41.4) | 4.50 |
| | Postcentral gyrus | R | | 48 | -34 | 60 | Area 1 (41.7) | 4.06 |
| | Supramarginal gyrus, post. div. | R | | 48 | -44 | 56 | Area PFm (IPL, 43.6) | 3.82 |
| | Angular gyrus | R | | 44 | -54 | 54 | Area PGa (IPL, 34.3) | 3.81 |
| | Superior parietal lobule | R | | 36 | -52 | 54 | Area hIP3 (IPS, 39.0) | 3.55 |
| | Cingulate gyrus, ant. div. | L | 1011 | -4 | 4 | 34 | – | 4.51 |
| | | R | | 10 | 20 | 28 | – | 4.31 |
| | Paracingulate gyrus | L | | -4 | 10 | 50 | Area 6mr / preSMA (47.4) | 4.29 |
| | Supplementary motor area | L/R | | 2 | -4 | 62 | Area 6mc / SMA (34.1) | 4.16 |
| | Presupplementary motor area | R | | 6 | 6 | 52 | Area 6mr / preSMA (58.5) | 4.02 |
| | Cingulate gyrus, post. div. | R | | 6 | -16 | 46 | Area 6mc / SMA (39.1) | 4.00 |
| | Precentral gyrus | R | | 2 | -16 | 56 | – | 3.99 |
| | Thalamus | R | 878 | 14 | -14 | 6 | – | 4.89 |
| | Frontal orbital cortex | R | | 34 | 30 | -4 | Area Id7 (38.8) | 4.57 |
| | Putamen | R | | 24 | 12 | 2 | – | 4.32 |
| | Frontal pole | R | | 44 | 36 | -2 | Area OP9 (21.3) | 4.26 |
| | Inferior frontal gyrus, pars op. | R | | 56 | 10 | 4 | Area 44 (47.4) | 4.13 |
| | Temporal pole | R | | 52 | 12 | -6 | Area TE 3 (50.8) | 3.94 |
| | Frontal pole | R | | 40 | 46 | 0 | – | 3.82 |
| | Thalamus | L | 596 | -14 | -16 | 6 | – | 4.70 |
| | Putamen | L | | -26 | 4 | 4 | – | 4.44 |
| | Pallidum | L | | -14 | 0 | 6 | – | 4.20 |
| | Insular cortex | L | | -34 | 8 | 0 | – | 3.95 |
| | Superior frontal gyrus | R | 499 | 28 | -4 | 66 | Area 6d1 (28.7) | 4.83 |
| | Middle frontal gyrus | R | | 44 | 2 | 54 | – | 4.62 |
| | Precentral gyrus | R | | 32 | -24 | 66 | Area 4a (28.1) | 3.80 |
| | | R | | 30 | -16 | 66 | Area 6d1 (31.1) | 3.74 |
| | Precuneus cortex | L | 391 | -8 | -58 | 58 | Area 7A (SPL, 18.7) | 4.02 |

| | | | | | | | |
|---|---|---|---|---|---|---|---|
| Intraparietal sulcus | L | | -40 | -50 | 56 | Area hIP3 (IPS, 38.4) | 3.97 |
| Superior parietal lobule | L | | -32 | -42 | 60 | Area 2 (34.7) | 3.95 |
| Supramarginal gyrus, post. div. | L | | -50 | -44 | 52 | Area hIP2 (IPS, 27.9) | 3.92 |
| Postcentral gyrus | L | | -46 | -38 | 58 | Area 2 (52.5) | 3.78 |
| Postcentral gyrus | R | 183 | 62 | -16 | 38 | Area 1 (39.4) | 4.32 |
| Supramarginal gyrus, ant. div. | R | | 60 | -28 | 46 | Area PF (IPL, 53.8) | 4.08 |
| | | | 56 | -30 | 50 | Area PFt (IPL, 40.3) | 4.00 |
| | | | 62 | -20 | 26 | Area PFop (IPL, 35.8) | 3.70 |
| Parietal operculum cortex | R | 20 | 56 | -34 | 26 | Area PFcm (IPL, 44.7) | 4.04 |
| Precentral gyrus | L | 14 | -24 | -24 | 70 | Area 6d1 (24.2) | 3.47 |
| Superior frontal gyrus | L | 1 | -24 | 6 | 68 | Area 6d1 (36.5) | 3.38 |

**Abbreviations.** Ant. div.: Anterior division, Cytoarchit. assignm.: Cytoarchitectonic assignment, dPMC: Dorsal premotor cortex, IPL: Inferior parietal lobule, IPS: Intraparietal sulcus, OA: Old adults, OP: Opercular cortex, PFm: Parietal area F, caudal part m, PFm: Parietal area F, part m, PFop: Parietal area, opercular subregion, PFt: Parietal area F, part t, PGa: Parietal area G, anterior, post. div.: posterior division, preSMA: Presupplementary motor area, RRC: Response–response congruent, RRI: Response–response incongruent, SMA: Supplementary motor area, SPL: Superior parietal lobule, YA: Young adults.

**Note.** Selection of peaks based on macroanatomical and cytoarchitectonic representative regions.

A different model was created for each seed (right IPS or dPMC) and each regressor ($Dual_{RRC}$ or $Dual_{RRI}$). Each connectivity contrast was estimated through the age-specific conjunction of the task regressor (+ or –). The contrasts were masked with the response-code conflict network ($Dual_{RRI}$ + > $Dual_{RRC}$ conjunction from the task-related GLM, inclusive mask at $p < 0.001$). All activations are reported at a cluster forming threshold with voxel-level uncorrected at $p < 0.001$, and cluster-level family-wise error (cFWE) corrected at $p < 0.05$. We used ImCalc to create a minimal conjunction of both main-effect results from the $Dual_{RRC}$ or $Dual_{RRI}$ regressors (Expression: min($Dual_{RRC}$, $Dual_{RRI}$)) for each seed separately.

**Table S12.**

Brain areas in which the connectivity with right dPMC and SPL was significantly modulated as a function of $Dual_{RRI}$ (vs. baseline) and $Dual_{RRC}$ (vs. baseline).

| Seed | *Contrast /* Macroanatomical structure | H | Cluster extent ($k_E$) | Cluster *p(cFWE)* | MNI coordinates x | y | z | Ciytoarchit. assignm. (Overlap in %) | Peak *Z* value |
|---|---|---|---|---|---|---|---|---|---|
| Right dPMC | ***YA: $Dual_{RRI>RRC}$ + ∩ OA: $Dual_{RRI>RRC}$ + (Mask: $Dual_{RRI}$ + > $Dual_{RRC}$)*** | | | | | | | | |
| | *not significant* | | | | | | | | |
| | ***YA: $Dual_{RRI>RRC}$ – ∩ OA: $Dual_{RRI>RRC}$ – (Mask: $Dual_{RRI}$ + > $Dual_{RRC}$)*** | | | | | | | | |
| | *not significant* | | | | | | | | |
| | ***YA: $Dual_{RRC}$ + ∩ OA: $Dual_{RRC}$ + (Mask: $Dual_{RRI}$ + > $Dual_{RRC}$)*** | | | | | | | | |
| | Superior frontal gyrus | R | 25407 | 0.000 | 24 | -6 | 54 | Area 6d3 (70.5) | > 8 |
| | | L | | | -22 | 4 | 52 | Area 6d3 (73.2) | 6.34 |
| | Precuneus cortex | R | | | 10 | -60 | 58 | Area 7A (SPL, 35.7) | 6.31 |
| | Supplementary motor area | R | | | 4 | -16 | 56 | Area 6mc / SMA (49.8) | 6.08 |
| | Insular cortex | L | | | -36 | 26 | -4 | Area Id7 (32.0) | 6.04 |
| | | R | | | 32 | 26 | 2 | Area Id7 (81.0) | 6.02 |
| | Middle frontal gyrus | R | | | 42 | 22 | 36 | – | 6.01 |
| | Cingulate gyrus, ant. div. | L/R | | | 0 | -4 | 48 | – | 5.98 |
| | Precuneus cortex | L | 3590 | 0.000 | -14 | -68 | 44 | Area hIP8 (IPS, 17.6) | 5.85 |
| | Superior parietal lobule | L | | | -18 | -58 | 56 | Area 7A (SPL, 40.1) | 5.28 |
| | Postcentral gyrus | L | | | -38 | -38 | 48 | Area 2 (33.1) | 5.14 |
| | Supramarginal gyrus, ant. div. | L | | | -48 | -32 | 42 | Area PFt (IPL, 48.4) | 5.11 |
| | Precuneus cortex | L | | | -14 | -52 | 52 | Area 5M (SPL, 12.8) | 5.09 |
| | Superior parietal lobule | L | | | -22 | -58 | 48 | Area hIP3 (IPS, 23.1) | 4.95 |
| | Middle temporal gyrus, TO part | R | 292 | 0.029 | 54 | -46 | 6 | Area PGa (IPL, 1.2) | 4.97 |
| | | R | | | 60 | -52 | -8 | – | 4.13 |
| | ***YA: $Dual_{RRC}$ – ∩ OA: $Dual_{RRC}$ – (Mask: $Dual_{RRI}$ + > $Dual_{RRC}$)*** | | | | | | | | |
| | *not significant* | | | | | | | | |
| | ***YA: $Dual_{RRI}$ + ∩ OA: $Dual_{RRI}$ + (Mask: $Dual_{RRI}$ + > $Dual_{RRC}$)*** | | | | | | | | |
| | Superior frontal gyrus | R | 2268 | 0.000 | 24 | -6 | 54 | Area 6d3 (70.5) | > 8 |
| | Precentral gyrus | R | | | 34 | -4 | 58 | Area 6d1 (5.2) | 5.46 |
| | Middle frontal gyrus | R | | | 36 | -2 | 56 | – | 5.40 |
| | Parietal operculum cortex | R | | | 60 | -32 | 26 | Area PF (IPL, 43.2) | 4.35 |
| | Postcentral gyrus | R | | | 38 | -30 | 56 | Area 3b (45.4) | 4.34 |
| | Superior parietal lobule | R | | | 34 | -50 | 60 | Area 7PC (SPL, 40.3) | 4.28 |
| | Supramarginal gyrus, ant. div. | R | | | 58 | -24 | 28 | Area PFcm (IPL, 32.4) | 4.20 |

| | | | | | | | | | |
|---|---|---|---|---|---|---|---|---|---|
| | Precentral gyrus | R | | | 26 | -28 | 54 | Area 4p (35.0) | 4.14 |
| | Angular gyrus | R | | | 42 | -50 | 54 | Area hIP3 (IPS, 45.8) | 4.14 |
| | Superior parietal lobule | L | 1225 | 0.000 | -32 | -58 | 50 | Area hIP3 (IPS, 59.5) | 4.93 |
| | Postcentral gyrus | L | | | -40 | -40 | 56 | Area 2 (60.2) | 4.78 |
| | Supramarginal gyrus, ant. div. | L | | | -48 | -34 | 46 | Area 2 (41.8) | 4.62 |
| | Superior parietal lobule | L | | | -16 | -70 | 54 | Area 7A (SPL, 45.4) | 3.85 |
| | Supramarginal gyrus, ant. div. | L | | | -58 | -32 | 38 | Area PFt (IPL, 61.0) | 3.66 |
| | Precuneus cortex | L | | | -10 | -64 | 56 | Area 7A (SPL, 54.1) | 3.62 |
| | Superior parietal lobule | L | | | -24 | -42 | 56 | Area 5L (SPL, 42.4) | 3.48 |
| | Precentral gyrus | L | 1169 | 0.000 | -28 | -10 | 56 | Area 6d1 (26.6) | 4.82 |
| | Inferior frontal gyrus, pars opercularis | L | | | -54 | 14 | 20 | Area 44 (65.9) | 4.07 |
| | Middle frontal gyrus | L | | | -50 | 8 | 36 | Area 44 (20.8) | 3.91 |
| | Superior frontal gyrus | L | | | -26 | -4 | 66 | Area 6d2 (39.4) | 3.90 |
| | | L | | | -22 | 2 | 54 | Area 6d3 (84.4) | 3.71 |
| | Supplementary motor area | R | 506 | 0.002 | 4 | -4 | 52 | Area 6mc / SMA (24.8) | 4.41 |
| | Juxtapositional lobule cortex | R | | | 8 | -14 | 48 | – | 4.26 |
| | Cingulate gyrus, ant. div. | R | | | 4 | 10 | 32 | Area 33 (48.2) | 3.73 |
| | Paracingulate gyrus | R | | | 6 | 16 | 38 | – | 3.65 |
| | Superior frontal gyrus | R | | | 2 | 20 | 52 | – | 3.22 |
| | Inferior frontal gyrus, pars opercularis | R | 454 | 0.003 | 46 | 8 | 18 | Area 44 (48.6) | 4.01 |
| | Insular cortex | R | | | 36 | 18 | 0 | Area Id7 (32.8) | 3.85 |
| | Frontal operculum cortex | R | | | 38 | 24 | 8 | Area OP9 (46.4) | 3.56 |
| | Insular cortex | L | 306 | 0.018 | -28 | 16 | -8 | – | 4.12 |
| | Frontal orbital cortex | L | | | -30 | 22 | -8 | – | 4.06 |
| | Putamen | L | | | -28 | 12 | 6 | – | 3.79 |
| | Middle frontal gyrus | L | 265 | 0.031 | -40 | 28 | 34 | – | 4.21 |
| | Frontal pole | L | | | -34 | 40 | 28 | – | 3.64 |
| | ***YA: $Dual_{RRI}$ – ∩ OA: $Dual_{RRI}$ – (Mask: $Dual_{RRI}$ + > $Dual_{RRC}$)*** | | | | | | | | |
| | *not significant* | | | | | | | | |
| **Right SPL** | ***YA: $Dual_{RRI>RRC}$ + ∩ OA: $Dual_{RRI>RRC}$ + (Mask: $Dual_{RRI}$ + > $Dual_{RRC}$)*** | | | | | | | | |
| | *not significant* | | | | | | | | |
| | ***YA: $Dual_{RRI>RRC}$ – ∩ OA: $Dual_{RRI>RRC}$ – (Mask: $Dual_{RRI}$ + > $Dual_{RRC}$)*** | | | | | | | | |
| | *not significant* | | | | | | | | |
| | ***YA: $Dual_{RRC}$ + ∩ OA: $Dual_{RRC}$ + (Mask: $Dual_{RRI}$ + > $Dual_{RRC}$)*** | | | | | | | | |
| | Superior parietal lobule | R | 16624 | 0.000 | 22 | -60 | 62 | Area 7A (SPL) | > 8 |
| | Thalamus | L | | | -8 | -20 | 0 | – | 6.73 |
| | | R | | | 12 | -16 | -2 | – | 6.62 |

| | | | | | | | | |
|---|---|---|---|---|---|---|---|---|
| Putamen | L | | | -26 | 14 | -2 | – | 6.36 |
| Pallidum | R | | | 18 | -4 | 6 | – | 5.95 |
| | L | | | -12 | 6 | -2 | – | 5.88 |
| Temporal pole | R | | | 48 | 16 | -18 | – | 5.85 |
| Caudate | R | | | 12 | 10 | -2 | – | 5.72 |
| Cerebellum VI | L | 2977 | 0.000 | -16 | -58 | -20 | – | 6.09 |
| Cerebellum Crus I | L | | | -28 | -66 | -32 | – | 5.73 |
| Cerebellum V | L | | | -24 | -40 | -26 | – | 5.02 |
| Cerebellum VI | R | | | 24 | -62 | -28 | – | 4.94 |
| Cerebellum VI | L | | | -38 | -50 | -26 | Area FG4 (32.8) | 4.91 |
| Temporal occipital fusiform cortex | L | | | -32 | -48 | -22 | Area FG3 (33.4) | 4.88 |
| Frontal pole | L | 1880 | 0.000 | -36 | 48 | 16 | – | 5.94 |
| Middle frontal gyrus | L | | | -42 | 36 | 20 | – | 5.15 |
| Inferior frontal gyrus, pars opercularis | L | | | -46 | 10 | 18 | Area 44 (46.5) | 4.42 |
| Superior parietal lobule | L | 1576 | 0.000 | -14 | -60 | 62 | Area 7A (SPL, 66.1) | 5.85 |
| | L | | | -12 | -66 | 60 | Area 7P (SPL, 42.0) | 5.76 |
| | L | | | -36 | -48 | 58 | Area 7PC (SPL, 46.3) | 4.32 |
| Supramarginal gyrus, post. div. | L | | | -46 | -46 | 54 | Area hIP2 (IPS, 29.8) | 4.12 |
| Precuneus cortex | L | | | -12 | -64 | 46 | Area 7A (SPL, 16.5) | 3.90 |
| Superior parietal lobule | L | | | -28 | -40 | 54 | Area 5L (SPL, 50.5) | 3.66 |
| Middle temporal gyrus, TO part | R | 337 | 0.026 | 62 | -54 | -8 | – | 5.22 |
| Inferior temporal gyrus, TO part | R | | | 58 | -54 | -16 | – | 5.14 |
| Middle temporal gyrus, TO part | R | | | 56 | -44 | 0 | – | 4.70 |
| Precentral gyrus | L | 303 | 0.037 | -36 | -6 | 60 | Area 6d2 (12.1) | 4.16 |
| Superior frontal gyrus | L | | | -24 | -8 | 72 | Area 6d1 (45.8) | 4.14 |
| Precentral gyrus | L | | | -24 | -24 | 70 | Area 6d1 (24.2) | 3.83 |
| Middle frontal gyrus | L | | | -32 | 4 | 62 | Area 6d3 (7.0) | 3.65 |
| ***YA: $Dual_{RRC}$ – ∩ OA: $Dual_{RRC}$ – (Mask: $Dual_{RRI}$ + > $Dual_{RRC}$)*** | | | | | | | | |
| *not significant* | | | | | | | | |
| ***YA: $Dual_{RRI}$ + ∩ OA: $Dual_{RRI}$ + (Mask: $Dual_{RRI}$ + > $Dual_{RRC}$)*** | | | | | | | | |
| Superior parietal lobule | R | 4065 | 0.000 | 18 | -60 | 62 | Area 7A (SPL, 70.3) | 6.72 |
| Inferior parietal lobule | R | | | 58 | -30 | 26 | Area PFcm (IPL, 48.3) | 4.89 |
| Superior frontal gyrus | R | | | 30 | -2 | 64 | Area 6d1 (8.0) | 4.77 |
| Precentral gyrus | R | | | 46 | 2 | 52 | – | 4.65 |
| Postcentral gyrus | R | | | 44 | -36 | 60 | – | 4.29 |
| Supramarginal gyrus, ant. div. | R | | | 58 | -24 | 46 | Area PFt (IPL, 51.4) | 4.29 |

| | | | | | | | | |
|---|---|---|---|---|---|---|---|---|
| Cingulate gyrus, ant. div. | L | | | -4 | 4 | 34 | – | 4.24 |
| Cerebellum VI | L | 1739 | 0.000 | -32 | -56 | -32 | – | 5.05 |
| | L | | | -20 | -60 | -26 | – | 5.00 |
| | L | | | -8 | -68 | -28 | – | 4.75 |
| Cerebellum Vermis VI | R | | | 2 | -62 | -28 | Interposed Nucleus (17.9) | 4.48 |
| Cerebellum VI | R | | | 22 | -62 | -28 | – | 4.45 |
| Cerebellum I-IV | L/R | | | 0 | -56 | -22 | – | 4.27 |
| Cerebellum V | L | | | -16 | -50 | -24 | – | 4.17 |
| Cerebellum Crus I | L | | | -48 | -62 | -34 | – | 3.49 |
| Thalamus | R | 919 | 0.000 | 14 | -14 | 6 | – | 4.55 |
| Frontal orbital cortex | R | | | 34 | 30 | -4 | Area Id7 (38.8) | 4.39 |
| Putamen | R | | | 24 | 12 | 2 | – | 4.07 |
| Temporal pole | R | | | 52 | 12 | -4 | Area 44 (40.9) | 4.04 |
| Frontal pole | R | | | 44 | 36 | -2 | Area OP9 (21.3) | 4.03 |
| Inferior frontal gyrus, pars opercularis | R | | | 56 | 10 | 4 | Area 44 (47.4) | 3.92 |
| Superior parietal lobule | L | 694 | 0.000 | -22 | -48 | 62 | Area 5L (SPL, 55.6) | 4.16 |
| Postcentral gyrus | L | | | -32 | -40 | 62 | Area 2 (39.3) | 4.09 |
| Intraparietal sulcus | L | | | -40 | -50 | 54 | Area hIP3 (IPS, 36.6) | 3.85 |
| Postcentral gyrus | L | | | -44 | -38 | 58 | Area 2 (56.7) | 3.85 |
| Precuneus cortex | L | | | -8 | -58 | 58 | Area 7A (SPL, 18.7) | 3.82 |
| Supramarginal gyrus, post. div. | L | | | -54 | -44 | 48 | Area hIP2 (IPS, 51.3) | 3.82 |
| Thalamus | L | 598 | 0.001 | -14 | -16 | 6 | – | 4.39 |
| Putamen | L | | | -26 | 4 | 4 | – | 4.17 |
| Pallidum | L | | | -14 | 0 | 6 | – | 3.98 |
| Insular cortex | L | | | -34 | 8 | 0 | – | 3.76 |
| ***YA: $Dual_{RRI}$ – ∩ OA: $Dual_{RRI}$ – (Mask: $Dual_{RRI}$ + > $Dual_{RRC}$)*** | | | | | | | | |
| *not significant* | | | | | | | | |

**Abbreviations.** Ant. div.: Anterior division, Cytoarchit. assignm.: Cytoarchitectonic assignment, dPMC: Dorsal premotor cortex, IPL: Inferior parietal lobule, IPS: Intraparietal sulcus, OA: Old adults, OP: Opercular cortex, PFcm: Parietal area F, part cm, PFt: Parietal area F, part t, PGa: Parietal area G, anterior, post. div.: posterior division, RRC: Response–response congruent, RRI: Response–response incongruent, SPL: Superior parietal lobule, TO: Temporooccipital, YA: Young adults.

**Note.** Selection of peaks based on macroanatomical and cytoarchitectonic representative regions.

A different model was created for each seed (right IPS or dPMC) and each regressor ($Dual_{RRC}$ or $Dual_{RRI}$). Each connectivity contrast was estimated through the age-specific conjunction of the task regressor (+ or –). The contrasts were masked with the response-code conflict network ($Dual_{RRI}$ + > $Dual_{RRC}$ conjunction from the task-related GLM, inclusive mask at $p < 0.001$). All activations are reported at a cluster forming threshold with voxel-level uncorrected at $p < 0.001$, and cluster-level family-wise error (cFWE) corrected at $p < 0.05$.

**Table S13.**

Age differences in the brain areas in which the connectivity with right dPMC and SPL was significantly modulated as a function of Dual$_{RRI}$ (vs. baseline) and Dual$_{RRC}$ (vs. baseline).

| Seed | *Contrast /* Macroanatomical structure | H | Cluster extent ($k_E$) | Cluster *p(cFWE)* | MNI coordinates | | | Ciytoarchit. assignm. (Overlap in %) | Peak *Z* value |
|---|---|---|---|---|---|---|---|---|---|
| | | | | | x | y | z | | |
| **Right dPMC** | ***Dual$_{RRI>RRC}$: OA + > YA (Mask: Dual$_{RRI}$ + > Dual$_{RRC}$)*** | | | | | | | | |
| | *not significant* | | | | | | | | |
| | ***Dual$_{RRI>RRC}$: YA + > OA (Mask: Dual$_{RRI}$ + > Dual$_{RRC}$)*** | | | | | | | | |
| | *not significant* | | | | | | | | |
| | ***Dual$_{RRC}$: OA + > YA (Mask: Dual$_{RRI}$ + > Dual$_{RRC}$)*** | | | | | | | | |
| | *not significant* | | | | | | | | |
| | ***Dual$_{RRC}$: YA + > OA (Mask: Dual$_{RRI}$ + > Dual$_{RRC}$)*** | | | | | | | | |
| | *not significant* | | | | | | | | |
| | ***Dual$_{RRI}$: OA + > YA (Mask: Dual$_{RRI}$ + > Dual$_{RRC}$)*** | | | | | | | | |
| | *not significant* | | | | | | | | |
| | ***Dual$_{RRI}$: YA + > OA (Mask: Dual$_{RRI}$ + > Dual$_{RRC}$)*** | | | | | | | | |
| | *not significant* | | | | | | | | |
| **Right SPL** | ***Dual$_{RRI>RRC}$: OA + > YA (Mask: Dual$_{RRI}$ + > Dual$_{RRC}$)*** | | | | | | | | |
| | *not significant* | | | | | | | | |
| | ***Dual$_{RRI>RRC}$: YA + > OA (Mask: Dual$_{RRI}$ + > Dual$_{RRC}$)*** | | | | | | | | |
| | *not significant* | | | | | | | | |
| | ***Dual$_{RRC}$: OA + > YA (Mask: Dual$_{RRI}$ + > Dual$_{RRC}$)*** | | | | | | | | |
| | *not significant* | | | | | | | | |
| | ***Dual$_{RRC}$: YA + > OA (Mask: Dual$_{RRI}$ + > Dual$_{RRC}$)*** | | | | | | | | |
| | Supramarginal gyrus, ant. div. | L | 1149 | 0.000 | -42 | -28 | 34 | Area 3a (12.3) | 5.12 |
| | | L | | | -54 | -36 | 42 | Area PFt (IPL, 47.3) | 4.53 |
| | | L | | | -64 | 30 | 32 | Area PFop (IPL, 52.5) | 4.42 |
| | Precuneus cortex | L | | | -14 | -68 | 48 | Area hIP8 (IPS, 26.4) | 4.31 |
| | Intraperietal sulcus | L | | | -26 | -56 | 46 | Area hIP3 (IPS, 33.6) | 4.30 |
| | Supramarginal gyrus, ant. div. | L | | | -38 | -40 | 36 | Area hIP1 (IPS, 61.5) | 3.89 |
| | Postcentral gyrus | L | | | -44 | -34 | 48 | Area 2 (80.6) | 3.13 |
| | Supramarginal gyrus, post. div. | R | 808 | 0.000 | 34 | -38 | 40 | Area hIP3 (IPS, 21.5) | 4.32 |
| | Supramarginal gyrus, ant. div. | R | | | 60 | -28 | 30 | Area PF (IPL, 43.2) | 3.94 |
| | Postcentral gyrus | R | | | 46 | -32 | 52 | Area 2 (63.9) | 3.88 |
| | Supramarginal gyrus, post. div. | R | | | 44 | -38 | 50 | Area hIP2 (IPS, 79.2) | 3.73 |

| | | | | | | | | |
|---|---|---|---|---|---|---|---|---|
| | R | | | 52 | -24 | 40 | Area PFt (IPL, 57.2) | 3.62 |
| Intraperietal sulcus | R | 458 | 0.008 | 22 | -68 | 46 | Area hIP8 (IPS, 11.2) | 4.48 |
| Superior parietal lobule | R | | | 14 | -62 | 58 | Area 7A (SPL, 51.3) | 4.14 |
| Frontal operculum cortex | R | 369 | 0.018 | 36 | 22 | 6 | Area OP9 (26.0) | 5.01 |
| Inferior frontal gyrus, pars opercularis | R | | | 48 | 18 | 6 | Area 44 (34.4) | 3.74 |
| Putamen | R | | | 26 | 14 | 0 | – | 3.36 |
| ***$Dual_{RRI}$: OA + > YA (Mask: $Dual_{RRI}$ + > $Dual_{RRC}$)*** | | | | | | | | |
| *not significant* | | | | | | | | |
| ***$Dual_{RRI}$: YA + > OA (Mask: $Dual_{RRI}$ + > $Dual_{RRC}$)*** | | | | | | | | |
| Superior parietal lobule | R | 354 | 0.014 | 30 | -38 | 40 | Area hIP3 (IPS, 34.2) | 4.12 |
| Postcentral gyrus | R | | | 38 | -32 | 40 | Area 2 (50.9) | 4.00 |
| Supramarginal gyrus, post. div. | R | | | 46 | -40 | 40 | Area hIP2 (IPS, 32.3) | 3.82 |
| Supramarginal gyrus, ant. div. | R | | | 46 | -30 | 40 | Area PFt (IPL, 37.3) | 3.82 |
| | R | | | 56 | -30 | 36 | Area PFcm (IPL, 32.8) | 3.50 |

**Abbreviations.** Ant. div.: Anterior division, Cytoarchit. assignm.: Cytoarchitectonic assignment, dPMC: Dorsal premotor cortex, IPL: Inferior parietal lobule, IPS: Intraparietal sulcus, OA: Old adults, PFcm: Parietal area F, part cm, PFop: Parietal area, opercular subregion, PFt: Parietal area F, part t, RRC: Response–response congruent, RRI: Response–response incongruent, SPL: Superior parietal lobule, YA: Young adults.

**Note.** Selection of peaks based on macroanatomical and cytoarchitectonic representative regions.

A different model was created for each seed (right IPS or dPMC) and each regressor ($Dual_{RRC}$ or $Dual_{RRI}$). Each connectivity contrast included the age contrast in conjunction with the regressor's main effect for the age group marked by +. The contrasts were masked with the response-code conflict network ($Dual_{RRI}$ + > $Dual_{RRC}$ conjunction from the task-related GLM, inclusive mask at $p < 0.001$). All activations are reported at a cluster forming threshold with voxel-level uncorrected at $p < 0.001$, and cluster-level family-wise error (cFWE) corrected at $p < 0.05$.
